# Identification and classification of cis-regulatory elements in the amphipod crustacean *Parhyale hawaiensis*

**DOI:** 10.1101/2021.09.16.460328

**Authors:** Dennis A Sun, Jessen V Bredeson, Heather S Bruce, Nipam H Patel

## Abstract

Emerging research organisms enable the study of biology that cannot be addressed using classical “model” organisms. The development of new data resources can accelerate research in such animals. Here, we present new functional genomic resources for the amphipod crustacean *Parhyale hawaiensis*, facilitating the exploration of gene regulatory evolution using this emerging research organism. We use Omni-ATAC-Seq, an improved form of the Assay for Transposase-Accessible Chromatin coupled with next-generation sequencing (ATAC-Seq), to identify accessible chromatin genome-wide across a broad time course of *Parhyale* embryonic development. This time course encompasses many major morphological events, including segmentation, body regionalization, gut morphogenesis, and limb development. In addition, we use short- and long-read RNA-Seq to generate an improved *Parhyale* genome annotation, enabling deeper classification of identified regulatory elements. We discover differential accessibility, predict nucleosome positioning, infer transcription factor binding, cluster peaks based on accessibility dynamics, classify biological functions, and correlate gene expression with accessibility. Using a Minos transposase reporter system, we demonstrate the potential to identify novel regulatory elements using this approach, including distal regulatory elements. This work provides a platform for the identification of novel developmental regulatory elements in *Parhyale*, and offers a framework for performing such experiments in other emerging research organisms.

**Primary Findings:**

– Omni-ATAC-Seq identifies cis-regulatory elements genome-wide during crustacean embryogenesis
– Combined short- and long-read RNA-Seq improves the *Parhyale* genome annotation
– ImpulseDE2 analysis identifies dynamically regulated candidate regulatory elements
– NucleoATAC and HINT-ATAC enable inference of nucleosome occupancy and transcription factor binding
– Fuzzy clustering reveals peaks with distinct accessibility and chromatin dynamics
– Integration of accessibility and gene expression reveals possible enhancers and repressors
– Omni-ATAC can identify known and novel regulatory elements

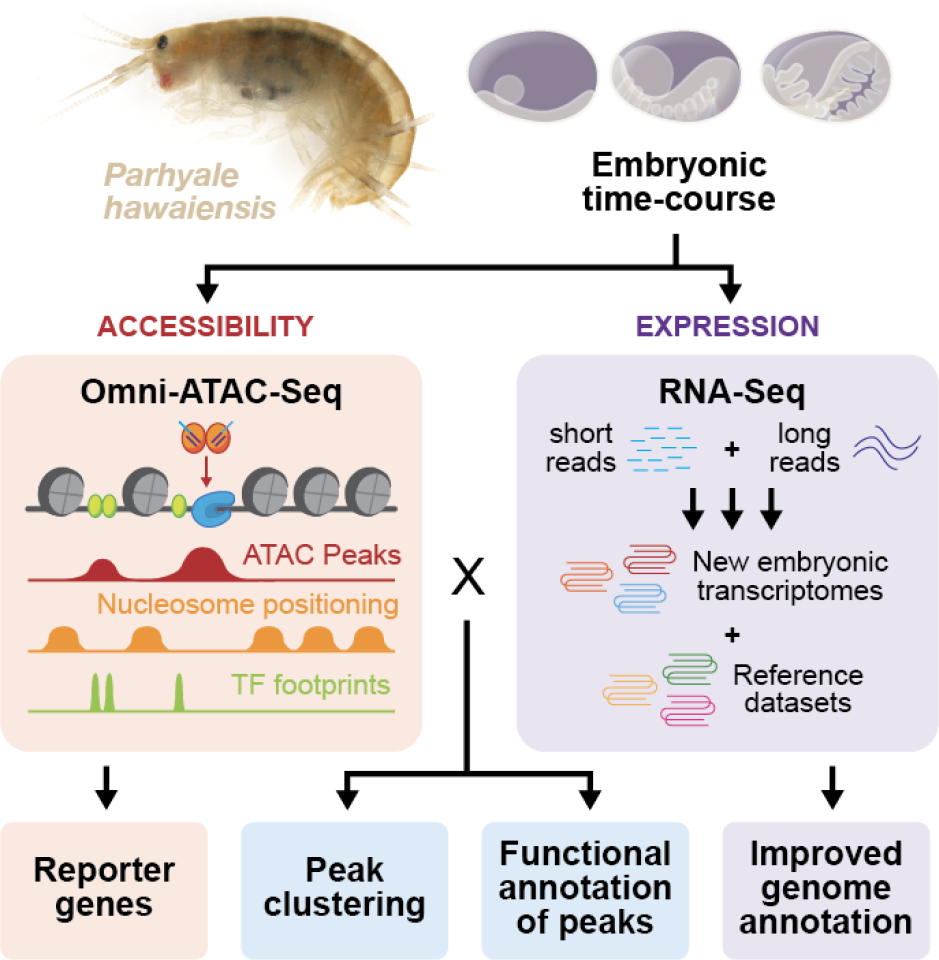

## Introduction

Advances in genomic techniques have facilitated genetic research in emerging research organisms. As sequencing tools become more sophisticated and less expensive over time, understanding what genes an organism expresses at any time in development becomes more straightforward. The expansion of genome sequencing has accelerated to such a degree that cost per gigabase of genome sequence has decreased at a rate faster than predicted by Moore’s Law (Wetterstramd). Moreover, the establishment of techniques, including RNAi and CRISPR-Cas9 mutagenesis, has made genetic ablation available to researchers working in a range of different research organisms, enabling the rapid characterization of gene function. RNAi knockdown has been applied to various insects (Mito et al., 2011; Schmitt-Engel et al., 2015), chelicerates (Sharma et al., 2013), flatworms (Rouhana et al., 2013; Srivastava et al., 2014), and numerous other organisms to study gene function. More recently, CRISPR-Cas9 mutagenesis has enabled both targeted ablation and targeted transgenesis in organisms as diverse as cephalopods (Crawford et al., 2020) and reptiles (Rasys et al., 2019). Using these tools, it is now possible to understand how genes and genomes evolve across the tree of life.

While changes to gene function have been shown to play some role in evolutionary processes, changes to gene regulation are thought to be a more important force in shaping the diversity of cell types, morphologies, and populations across clades (Mattioli et al., 2020; Signor and Nuzhdin, 2018; Wittkopp et al., 2004). The evolution of gene regulation is governed by changes in the function of two gene-regulatory modes: changes to trans-regulatory factors and changes to cis-regulatory elements. Trans-regulatory factors may include transcription factors, noncoding RNAs, and other diffusible elements that modify the expression of distant genes, while cis-regulatory elements are considered regions of the genome that locally mediate gene regulation.

The identification and characterization of possible trans-regulatory factors in a genome can be accomplished with relative ease using genome sequencing. For example, functional annotation of transcriptomes is able to easily identify transcription factors based on highly conserved protein domains (Ofran et al., 2007; Wang and Brown, 2006). Many bioinformatic tools also exist to identify microRNAs and long noncoding RNAs through sequence-based approaches (Duan et al., 2020; Liu et al., 2014).

RNA-Seq approaches can also enable the identification of differentially expressed transcripts which might be important for biological function (Costa-Silva et al., 2017). Thus, it is possible to identify many of the trans-regulatory factors within an organism’s genome with straightforward RNA sequencing approaches.

Cis-regulatory elements (CREs), on the other hand, have historically proven difficult to identify even in model organisms. Unlike protein-coding sequences, cis-regulatory elements are composed of complex assemblies of transcription factor binding sites, which can be computationally challenging to identify (Li et al., 2015; Wasserman and Sandelin, 2004). These sites can change dramatically over short periods of time; for example, in the context of the *even-skipped* stripe 2 enhancer across several *Drosophila* species, the position of critical transcription factor binding sites shifted in as short a span as 1–2 million years divergence (Ludwig et al., 1998). Such rapid evolutionary flexibility, while important for driving short- and long-term evolutionary processes, makes the identification of such elements in novel research organisms a considerable challenge.

In recent years, novel techniques for identifying CREs have been developed, which rely on the fact that CREs often occur in regions of accessible chromatin. The earliest, including DNAse I hypersensitivity and Formaldehyde-Assisted Isolation of Regulatory Elements (FAIRE-Seq) enabled researchers to locate accessible chromatin regions and discover novel regulatory elements in a range of research organisms (Giresi et al., 2007; Lai et al., 2018; Pérez-Zamorano et al., 2017). However, such techniques were hampered by low signal-to-noise ratios, the requirement for very large amounts of tissue, and technically challenging protocols. In 2013, Buenrostro et al. published papers describing the Assay for Transposase-Accessible Chromatin with next-generation sequencing (ATAC-Seq), a novel technique using a hyperactive version of the Tn5 transposase, as a way of identifying accessible chromatin regions genome-wide (Buenrostro et al., 2015). In this paper, they demonstrated that ATAC-Seq performed comparably to previous accessibility techniques with reduced tissue requirements and increased convenience and speed.

ATAC-Seq has proved to be an effective technique for traditional model systems, and has offered a true revolution for emerging model systems. The low material input requirements of ATAC-Seq are of particular use in emerging research organisms, for which generating the millions of cells required for previous techniques would prove challenging. Researchers have applied ATAC-Seq to gar embryos in order to study the evolution of limb development (JL and Shubin, 2015) and regenerating acoel flatworms and hydra to study regeneration (Cazet et al., 2021; Gehrke et al., 2019), among numerous other examples. Thus, ATAC-Seq is now established as a generalizable tool that is particularly well-suited to accelerating work in emerging research organisms.

In 2017, Corces et al. reported a further-improved version of ATAC-Seq, Omni-ATAC-Seq, which provides several improvements to the standard ATAC-Seq protocol (Corces et al., 2017). In particular, it has been reported to have a greater signal-to-noise ratio, decreased mitochondrial read contamination, and applicability to fixed and frozen tissues, with only minor modifications to the buffers and detergents used in the standard ATAC-Seq protocol.

In this paper, we apply the Omni-ATAC-Seq protocol to embryos of the amphipod crustacean *Parhyale hawaiensis*, an emerging research organism for the study of arthropod development, evolution, and regeneration (Paris et al., 2021; Stamataki and Pavlopoulos, 2016; Sun and Patel, 2019). We identify dynamic regions of open chromatin (“peaks”) across a broad swath of *Parhyale* developmental time. We comprehensively analyze our Omni-ATAC-Seq data to predict the position of nucleosomes along the genome and infer the footprints of transcription factors bound to peaks. Using fuzzy clustering (Kumar and E Futschik, 2007), we partition our peaks into groups based on similar accessibility trajectories, revealing groups of peaks with different transcription factor footprint enrichment and nucleosome occupancy that are active at different points in development. In addition, we use short- and long-read RNA-Seq to improve the *Parhyale* genome annotation and investigate the relationship between accessibility and gene expression over time during development.

*Parhyale* has served as a platform for foundational discoveries about such processes as body plan evolution (Martin et al., 2015), the evolution of arthropod limbs (Bruce and Patel, 2020), and the evolution of regeneration (Konstantinides and Averof, 2014). By facilitating the identification of regulatory elements in this research organism, our work will enable other researchers to investigate the complexities of gene regulatory evolution in these processes and others. Furthermore, by enabling the assessment of cis-regulatory elements in *Parhyale*, we open up avenues to investigate fundamental mechanisms of gene regulation in the understudied non-insect crustacean clade.

While this work primarily focuses on *Parhyale*, our methods can provide examples of how one can perform thorough analyses of ATAC-Seq and RNA-Seq data using existing tools to generate hypotheses about gene expression dynamics and regulatory element function. Such approaches can be applied to a diverse range of organisms, and will facilitate deeper understanding of gene regulation across the tree of life.

## Results

### Omni-ATAC-Seq identifies open chromatin across *Parhyale* developmental stages

To identify developmental regulatory elements, we performed Omni-ATAC-Seq on 15 stages of *Parhyale* embryonic development (Fig. 1). For each developmental stage, we generated two Omni-ATAC-Seq libraries using five embryos each, and fixed and DAPI stained sibling embryos to confirm staging (Fig. 1A). The developmental stages catalogued in our dataset are illustrated in Fig. 1B, and representative fixed, DAPI-stained sibling embryos are illustrated in Fig. 1C. Additional brightfield images of embryos prior to tagmentation can be found in Supp. Fig. 1A.

**Fig. 1:**
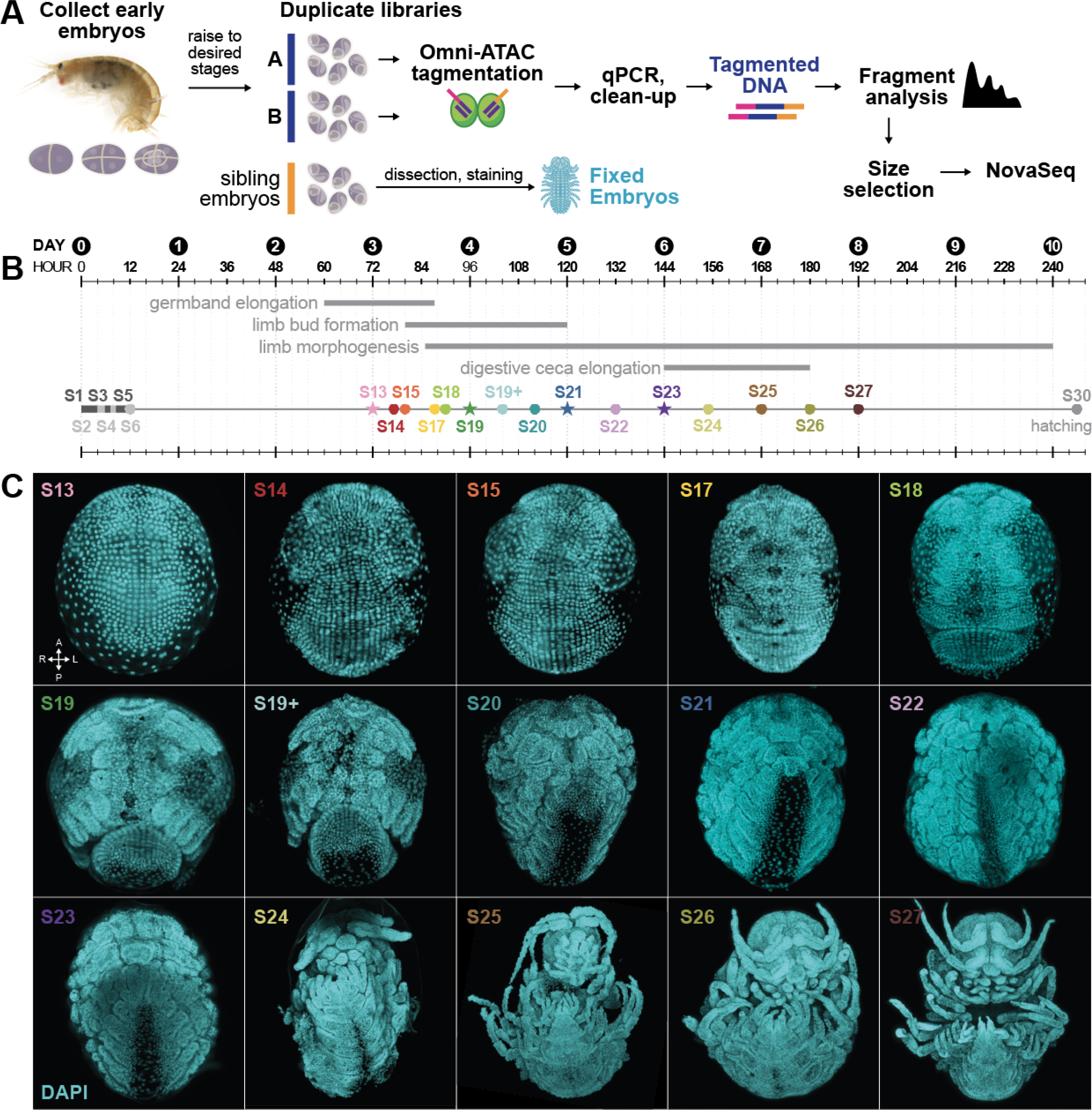
Time-course ATAC-Seq in *Parhyale hawaiensis* embryos. A) Overview of ATAC-Seq protocol. Embryos were collected at early developmental stages and raised to specific developmental timepoints. Duplicate libraries were generated by tagmenting 5 embryos each from a single clutch of sibling embryos. qPCR analysis was used to determine the optimum number of additional cycles of PCR. Tagmented DNA was cleaned and fragment analysis was performed to assess library quality. Final libraries were pooled in equal concentrations, size-selected, and sequence on an Illumina NovaSeq short read sequencer. B) Timeline of developmental stages. RNA-Seq libraries were also generated for timepoints marked with a star. C) Representative embryo images from clutches used for Omni-ATAC-Seq. Embryos are stained with DAPI and mounted ventral-side up. (A: anterior, P: posterior, L: left, R: right).

To assess the quality of our libraries, we employed a battery of standard tests used to assess the quality of ATAC-Seq data (Supp Fig. 2A). Omni-ATAC-Seq libraries were also compared to previous ATAC-Seq libraries (see Methods), and had greater library size, higher numbers of non-duplicated reads, and lower amounts of mitochondrial DNA contamination (Supp. Fig. 2B, C). Fragment size analysis of Omni-ATAC-Seq reads after mapping revealed distinct 1-nucleosome and 2-nucleosome peaks, indicating that tagmentation was efficient (Supp Fig. 2E, F).

**Fig. 2:**
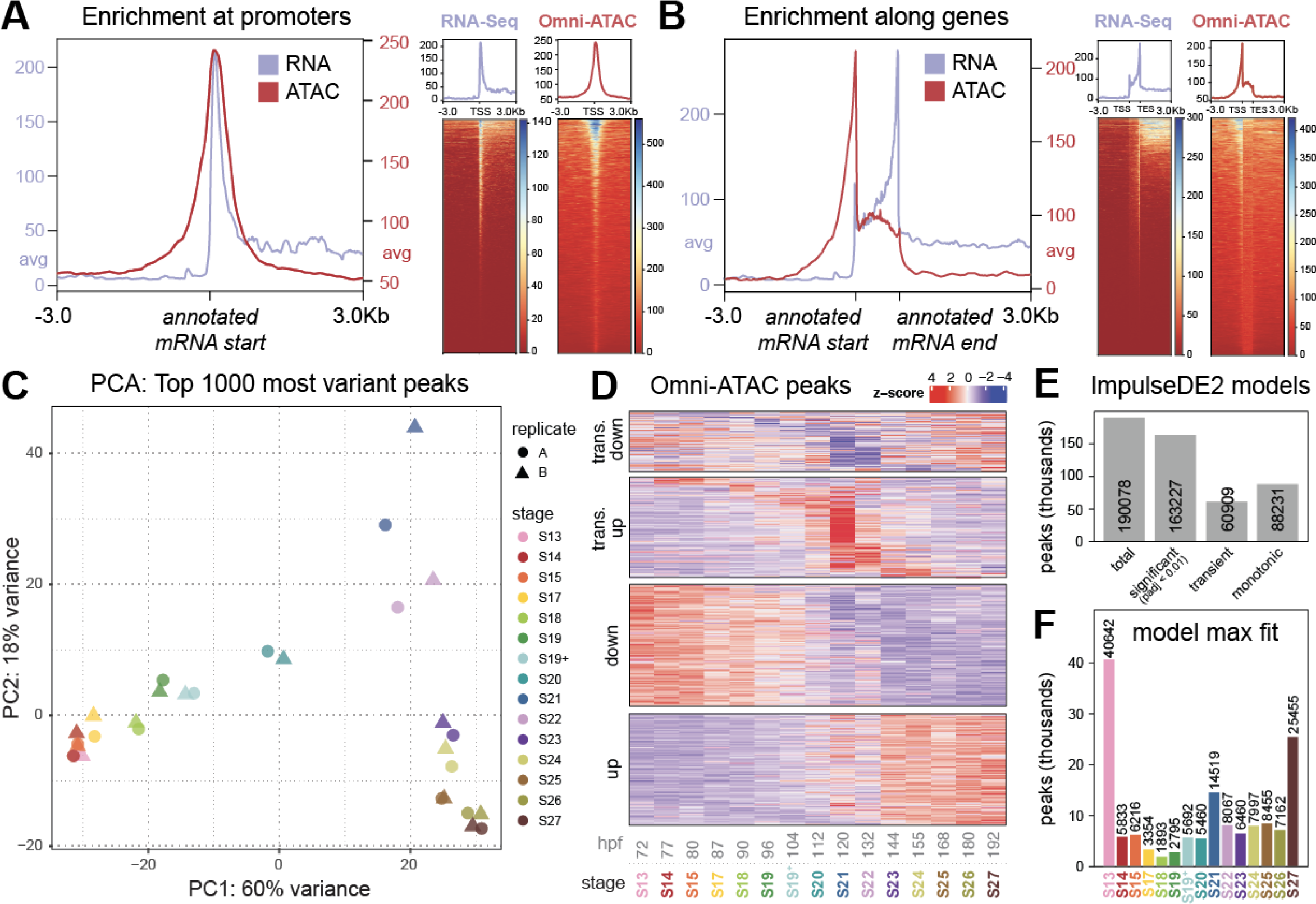
Whole-genome analysis of ATAC-Seq reveals enrichment of promoter signals, dynamic accessibility over developmental time. A) Omni-ATAC and RNA-Seq signal enrichment at promoters. Annotated mRNA start positions are aligned, and average signal across genomic positions is plotted for either Omni-ATAC or RNA-Seq signal. Axes reflect the average signal of RNA-Seq or Omni-ATAC reads across all mRNA start positions in the genome. A strong enrichment of RNA-Seq signal is observed 3’ of mRNA start sites, while enrichment of Omni-ATAC-Seq signal appears symmetric around mRNA start sites. B) Omni-ATAC and RNA-Seq signal enrichment across gene bodies. Omni-ATAC signal is greatly enriched 5’ of gene bodies and slightly enriched across genes, while RNA-Seq reads are enriched across gene bodies, with an increase in enrichment towards the 3’ end of gene bodies. C) PCA of Omni-ATAC libraries, generated using DESeq2. PCA loadings were assigned to the top 1000 variant peaks, where variance was measured as row variance across all libraries. PC1 (60% of variance) appears to correlate with developmental time. PC2 (18% variance) appears to be associated with middle developmental timepoints. D) Heatmap of Omni-ATAC accessibility dynamics, generated from ImpulseDE2 model fits. Colors reflect a Z-score calculated from ImpulseDE2, where red (high Z) indicates increase in accessibility and blue (low Z) indicates decrease in accessibility. E) Bar chart indicating number of peaks classified as significant, transient, and monotonic by ImpulseDE2. F) Bar chart quantifying stage of maximum accessibility for significant ImpulseDE2 model fits. Peaks predominantly achieved maximum accessibility at the start and end of the time course.

To determine whether our Omni-ATAC-Seq data represented genuine enrichment in accessible chromatin regions, we examined promoter accessibility genome-wide. A hallmark of successful ATAC-Seq experiments is strong enrichment of reads in gene promoters (Yan et al., 2020). We evaluated our ATAC-Seq using the most recent phaw_5.0 gene annotations (obtained from Leo Blondel, hereafter referred to as the “MAKER annotation”) at the start of annotated mRNAs and across mRNA lengths, and compared our results to RNA-Seq read pileups (Fig. 2A–B). As expected, Omni-ATAC-Seq signal is enriched symmetrically at mRNA starts, whereas RNA-Seq reads are enriched 3’ of mRNA starts. In addition, Omni-ATAC-Seq data shows greater enrichment at promoters than over annotated mRNA regions, and decreased enrichment outside of annotated mRNAs. These data suggest that our Omni-ATAC-Seq performed as expected in identifying promoter regions genome-wide.

To assess whether our libraries were capable of capturing significant variation over time, we performed Principal Component Analysis (PCA) on our libraries (Fig. 2C). PCA revealed that our libraries were primarily separated along two principal components (PCs), PC1 (60% of variation) and PC2 (18% of variation), with a considerable drop in variation explained in other PCs (Supp. Fig. 2D). PC1 appeared to be associated with developmental time, with earlier developmental stage libraries showing a negative loading, and later developmental stages showing a positive loading.

To identify regions of dynamically accessible chromatin, we used the ImpulseDE2 (IDE2) pipeline (Fischer et al., 2018). IDE2 differs from other software for differential expression analysis in that it allows the investigation of trajectories of dynamic expression over large numbers of timepoints. Identifying differential expression is difficult for more categorical differential expression software such as edgeR and DESeq2, which use pairwise comparisons between timepoints to assess change over time (Love et al., 2014; Robinson et al., 2010). In addition, IDE2 enables the detection of transient changes in gene expression or accessibility during a time course.

We used Genrich (Gaspar) to call significant peaks (q < 0.05) in each of our 15 timepoints independently, and merged overlapping peaks across timepoints using bedtools (Quinlan and Hall, 2010), yielding 190,078 genomic regions which we used as our “peaks” in downstream analyses, including IDE2. The dynamic accessibility of these peaks is illustrated in a heatmap in Fig. 2D. Of these peaks, 163,227 (85.87%) were classified as having statistically significant variation (padj < 0.01) over our time-course; 60,909 (37.32%) were classified as having transient expression dynamics; and 88,231 (54.05%) were classified as showing positively or negatively monotonic expression dynamics. These results indicate that we were able to identify many dynamically accessible regions across the *Parhyale* genome.

IDE2 produces a fitted model of accessibility defined as the product of two sigmoid functions for each of the peaks used in the analysis (Fischer et al., 2018). We used these model fits to summarize the global properties of the dynamic peaks we identified. First, we estimated the number of models that achieved their maximum accessibility during each of the 15 timepoints sampled (Fig. 2F). The timepoints during which models most frequently showed their highest accessibility were S13, S21, and S27, representing approximately the earliest, middle, and latest timepoints.

We then used these IDE2 model fits to understand how the variance of our libraries captured by the PCs was related to accessibility dynamics (Supp. Fig. 2G). We extracted the 100 peaks with the most positive and negative loadings in PC1 and PC2 and plotted their IDE2 model fits. Based on these plots, we inferred that PC1 captured variation along developmental time, with positively-loaded peaks showing a “low early, high late” accessibility trajectory, and negatively-loaded peaks showing a “high early, low late” accessibility trajectory. Peaks with high positive loading in PC2 appeared to be those with highest accessibility in the middle developmental stages.

These global analysis results indicate that our Omni-ATAC-Seq experiments captured information normally found in ATAC-Seq data, including strong enrichment at promoters. Overall, the low mitochondrial read contamination, large library size, and low fraction of duplicated reads suggest that our Omni-ATAC-Seq data are of high quality. In addition, the results of the IDE2 differential accessibility analyses indicate that the vast majority of accessible regions in the *Parhyale* genome show dynamic accessibility over developmental time.

### Improving the *Parhyale* genome annotation using short- and long-read RNA-Sequencing

While Omni-ATAC-Seq signal showed enrichment across annotated mRNA starts in a genome-wide analysis, careful examination of individual genes and gene models indicates that many of the MAKER gene annotations are fragmented. For example, among the *Parhyale Hox* genes, 7 of 9 the genes showed discrepancies in annotation compared to previously-published Rapid Amplification of cDNA Ends (RACE) data (Serano et al., 2016), with *lab, pb*, *Ubx*, and *abd-A* showing fragmented gene models (Supp. Fig. 3.1A, B).

**Fig. 3:**
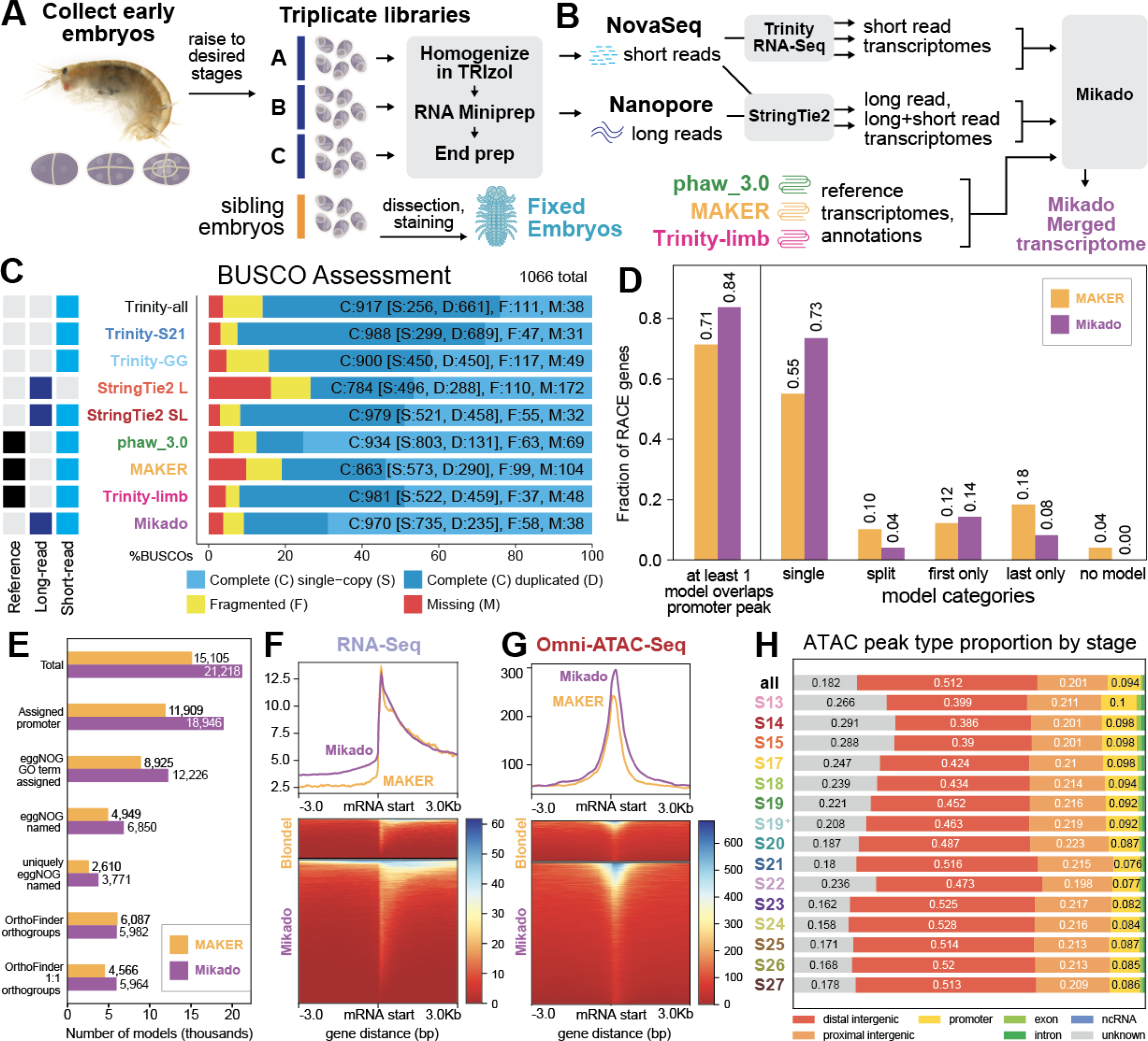
Time-course RNA-Seq of Parhyale embryos. A) RNA-Seq protocol overview. Triplicate libraries were generated for four developmental stages. B) RNA-Seq transcriptome assembly and merging pipeline. Short-read, long-read, and long- + short-read transcriptomes were generated for pooled developmental stages matched to Omni-ATAC timepoints (S13, S19, S21, S23) using Trinity and StringTie2. Transcriptomes were also assembled using Trinity for short reads from a limb developmental time course (see Methods). Assembled transcriptomes were then merged with additional transcripts from previous publications into a Mikado transcriptome. C) BUSCO scores for representative datasets and Mikado transcriptome. Overall, the StringTie2 short + long -read transcriptome (StringTie2 SL) performed comparably to the best Trinity transcriptomes. The Mikado transcriptome appeared to have a comparable BUSCO complement to the other datasets with high BUSCO completeness. D) Gene model completeness for Mikado transcriptome vs. MAKER genome annotation based on a dataset of 49 RACE genes. The Mikado transcriptome appears to have more complete gene models and less fragmentation. E) Comparison of number of gene models of different functional annotation categories between MAKER genome annotation and Mikado transcriptome. The Mikado transcriptome appears to produce more annotated genes with a higher annotation quality. F) Plot comparing RNA-Seq signal pileups between MAKER mRNA starts and Mikado mRNA starts. Axis represents mean signal at mRNA start sites for each dataset. G) Plot comparing Omni-ATAC-Seq signal pileups between MAKER mRNA starts and Mikado mRNA starts. H) Categorization of Omni-ATAC-Seq peaks in each stage-specific ATAC-Seq library by their position relative to Mikado gene models. Distal intergenic peaks are defined as those greater than 10kb away from the nearest gene, whereas unknown peaks were peaks that were located on a contig that did not have a gene model. A large majority of peaks (>71%) were intergenic peaks, and a majority of peaks (>51%) were distal intergenic peaks. The proportion of peaks of each type appeared relatively stable across developmental stages.

A more accurate genome annotation would improve both genome-wide analyses and enable the precise classification of candidate regulatory elements as promoters, exonic and intronic regulatory elements, and intergenic regulatory elements. To generate a more complete genome annotation, we performed RNA-Seq using two approaches: short-read sequencing using the Illumina NovaSeq platform and long-read sequencing using Oxford Nanopore technology. For four developmental stages (S13, S19, S21, and S23), we generated triplicate RNA-Seq libraries, which were sequenced using a short-read approach (Illumina NovaSeq) and a long-read approach (Oxford Nanopore) (Fig. 3A, B; representative embryo images in Supp. Fig. 1B).

We assembled multiple transcriptomes from each of the sequencing approaches (Fig. 3B). Using the Trinity (Haas et al., 2013) pipeline to assemble the short-read sequences, we generated four developmental stage-specific *de novo* transcriptomes (Trinity-S13 to Trinity-S23) and two merged transcriptomes, one *de novo* (Trinity-all) and one genome-guided (Trinity-GG). Additionally, we generated two transcriptomes with reads from several developmental stages covering the time course of limb development (Trinity-limb (new) and Trinity-limb (old), see Methods). Although the merged transcriptomes were generated from all four developmental stages (Trinity-all, Trinity-GG), they scored lower compared to stage-specific transcriptomes when CDS completeness was assessed using BUSCO (Fig 3D). One possible explanation for this could be an increase in transcript assembly fragmentation from de Bruijn graph assembly caused by the high heterozygosity previously described of the *Parhyale* genome (Kao et al., 2016). For long-read sequencing, we mapped the reads to the phaw_5.0 genome using minimap2 and assembled a transcriptome using StringTie2 (Kovaka et al., 2019; Li, 2018). We also used StringTie2 to generate a combined transcriptome containing both short and long reads. While the long-read transcriptome (StringTie2 L) yielded a low BUSCO score, the short- + long-read transcriptome (StringTie2 SL) scored comparably to our other transcriptomes (Fig. 3D).

To generate a more complete genome annotation, we used the Mikado pipeline to merge our assembled transcriptomes with the previously-generated Kao et al. 2016 transcriptome and the MAKER genome annotation. The Mikado pipeline and others (e.g. EvidentialGene) take advantage of the previously observed variable ability of transcriptome assemblers to produce complete gene models for different types of genes (Gilbert, 2019; Venturini et al., 2018). We evaluated the Mikado-merged transcriptome (hereafter “Mikado transcriptome”) using BUSCO, and observed a comparable score to the best of the individual transcriptomes (90.9% complete), and a marked improvement compared to the MAKER annotation (80.9% complete).

We first examined the Mikado transcriptome models representing the *Parhyale Hox* cluster genes and compared those to the RACE data and the MAKER annotation (Supp. Fig. 3 C–E). The gene models from the Mikado transcriptome showed an improvement relative to the MAKER models overall, with complete models for *pb, Hox3, Dfd* and *Scr*, which were fragmented in the MAKER annotation. While the Mikado pipeline was only able to recover one transcript isoform each for the three posterior *Hox* genes, for *abd-A* and *Ubx*, the isoforms identified were more complete than those the MAKER annotation.

Moreover, the Mikado-merged transcriptome was able to produce a more complete gene model for *Hox3* than the previous RACE data or MAKER annotation. Combining the RACE data with our Mikado transcriptome, we have assembled the most complete annotation of the *Parhyale Hox* cluster to date.

To quantify the level of gene model fragmentation in our dataset, we generated a series of manual gene annotations based on RACE sequences. Among 143 previously-generated RACE transcripts, we selected 49 multi-exonic transcripts that appeared to have a single promoter based on the Omni-ATAC-Seq data (Supp. Fig. 3D). For each of these RACE transcripts, we manually annotated the extent of the first and last exon by comparing RNA-Seq read pileups to RACE data and the current genome annotation. We also identified the Omni-ATAC-Seq peak most likely to capture the gene promoter, based on the strength of the peak (as evaluated by the number of timepoints over which we observed a statistically significant peak), as well as overlap to RNA-Seq read data. We used these manual annotations (see Supp. Table 2) to evaluate the gene models from each of the different transcript sources (transcriptomes or gene annotations).

For each of the 49 genes in our dataset, we evaluated whether any models in each of the transcript sources in our dataset overlapped the promoter peak (Supp. Fig. 3E). The ability to unambiguously identify promoters is essential for downstream analyses, including building reporter constructs or targeting CRISPR guides to the 5’-most end of genes. Among the transcript sources, the Trinity *de novo* transcriptomes had the highest proportion (0.96) of genes where at least one model overlapped with the promoter peak. However, the Trinity models we observed often (0.98, 48/49 RACE genes examined for the Trinity-limb transcriptome) contained numerous spurious transcript fragments in introns, exons, and 3’UTR regions (see Supp. Fig. 3.2 for summary statistics and examples). Among the remaining transcript sources, the Mikado transcriptome had the highest fraction of genes where at least one model overlapped with the promoter peak (0.84).

In addition, we assessed the degree of fragmentation of gene models across the 49 RACE transcripts (Supp. Fig. 3.1D, Methods). For each gene, we determined first whether any single gene model spanned the first and last exon, representing a complete “single” transcript. If there was not a single model, we next assessed whether two separate models overlapped with the first and last exon, or if all models only overlapped either the first or last exon. These results together formed a measure of transcript completeness. Comparing the different transcript sources, the StringTie2 SL transcriptome had the highest fraction (0.76) of complete “single” transcripts, with the Mikado transcriptome having a slightly lower fraction of “single” transcripts (0.73)(Supp. Fig. 3E).

When we directly compared the completeness measures between the Mikado transcriptome and the MAKER genome annotation, we observed a marked improvement. Notably, the Mikado transcriptome had a greater proportion of RACE genes with promoter-peak overlap (Mikado 0.84 > MAKER 0.71), as well as a greater fraction of “single” transcripts (Mikado 0.73 > MAKER 0.55)(Fig. 3D).

To further evaluate the quality of the Mikado transcriptome as a reference, we compared its performance to the MAKER annotation using a variety of metrics (Fig. 3E). The Mikado transcriptome produced a larger number of total gene models (n = 21,218) than are found in the MAKER annotation (n = 15,105). Using bedtools, we attempted to assign a candidate promoter peak to each transcript in each genome annotation (see Methods), and observed that the Mikado transcriptome had a larger number of transcripts with an assigned promoter (Mikado n = 18,946; MAKER n = 11,909), further suggesting improved genome-wide gene completeness.

We further examined the quality of the Mikado and MAKER transcriptomes by evaluating Omni-ATAC-Seq signal and RNA-Seq signal genome-wide at mRNA starts (Fig. 3F, G). We observed a comparable RNA-Seq signal, but a higher Omni-ATAC-Seq signal for the Mikado transcriptome as compared to the MAKER annotation, suggesting a stronger enrichment of promoters in our dataset.

These results together suggest that the Mikado transcriptome produces a more complete genome annotation, capturing more of the *bona fide* genes in the *Parhyale* genome than previously available.

To further assess the quality of the two genome annotations, we performed automated functional annotations to assign gene names and functions. We used two approaches: eggNOG and OrthoFinder (Emms and Kelly, 2019; Huerta-Cepas et al., 2019). eggNOG is a rapid and lightweight genome annotation software, and assigns gene names, KEGG pathway information, and GO terms, among numerous other metrics, to gene models. Trinotate is a more comprehensive transcript annotation pipeline, while OrthoFinder facilitates the identification of orthogroups between provided peptide libraries.

We observed that the Mikado transcriptome included a greater number of GO-term assigned genes, eggNOG named genes, and eggNOG uniquely-named genes than found in the MAKER annotation (Fig. 3E). We used OrthoFinder to assign orthogroups between the Mikado and MAKER transcriptomes and the list of all *Drosophila melanogaster* peptides from the UNIPROT-SWISSPROT database. The Mikado and MAKER transcriptomes produced similar numbers of orthogroups; however, the Mikado transcriptome produced a greater number of 1:1 orthogroups, which proved useful for downstream GO term enrichment analysis (summarized in Fig. 3E, see Supp. Fig. 3.3 for further explanation of orthogroup size comparisons).

Overall, we observed an improvement across nearly all metrics we used to compare the Mikado transcriptome to the MAKER annotation. The Mikado transcriptome appeared to produce more complete gene models than in the MAKER annotation, and more of these models could be annotated using automated methods such as eggNOG and OrthoFinder. Moreover, the Mikado transcriptome draws from a diverse range of tissue sources, as the previous Kao transcriptome and MAKER annotations that contribute to the Mikado transcriptome also incorporated data from adult tissues and other embryonic stages (See Supp. Fig. 3.1G for contributions to Mikado from each transcript source). Given these results, we used the Mikado transcriptome as our reference genome annotation for downstream analyses.

Using the Mikado transcriptome as a reference, we assigned Omni-ATAC-Seq peaks to a number of different spatial categories along the genome (Fig. 3J, see Methods). We observed that intergenic peaks make up the vast majority (71.3%) of Omni-ATAC peaks. We further partitioned the intergenic peaks into proximal and distal segments, with distal intergenic peaks representing those peaks >10kb away from the nearest gene. The majority of peaks in our dataset (51%) were distal intergenic peaks, indicating that much of the *Parhyale* regulatory landscape is composed of distal regulatory elements.

Nearly a fifth of regulatory elements (18.2%) were also located on contigs that did not contain genes, and were classified as “unknown”.

As our peaks represent the merged combination of peaks across individual timepoints, we also evaluated the proportion of peaks from each time-point belonging to each spatial category (Fig. 3J). Early developmental stages appeared to have a greater proportion of peaks in “unknown” regions, while later developmental stages appeared to have an increased number of peaks in intergenic regions, with the number of promoter, exonic, and intronic peaks remaining similar over time.

### Inferring nucleosome positioning and transcription factor footprints from Omni-ATAC-Seq

ATAC-Seq and other accessibility approaches primarily identify regions of open chromatin in the genome. However, with sequencing performed at sufficient depth, one can infer additional information from the pattern of insertions of ATAC-Seq adapters into the genome. Using our Omni-ATAC data, we predicted the temporal positioning of nucleosomes near Omni-ATACpeaks using NucleoATAC and inferred the footprints of transcription factors bound to accessible peaks using HINT-ATAC.

NucleoATAC enables the prediction of nucleosome positioning from ATAC-Seq data (Schep et al., 2015). We applied NucleoATAC to a window of +/- 500bp around our Omni-ATAC-Seq peaks at each developmental stage. NucleoATAC depends on a stereotypical signal of Tn5 insertion around nucleosomes referred to as a “V-plot”, which resembles the fragmentation pattern around chromatin subjected to chemical fragmentation. We observed across developmental time-points a clear V-plot signal generated by NucleoATAC (example of V-plot from S21 libraries shown in Supp. Fig. 4A).

**Fig. 4.**
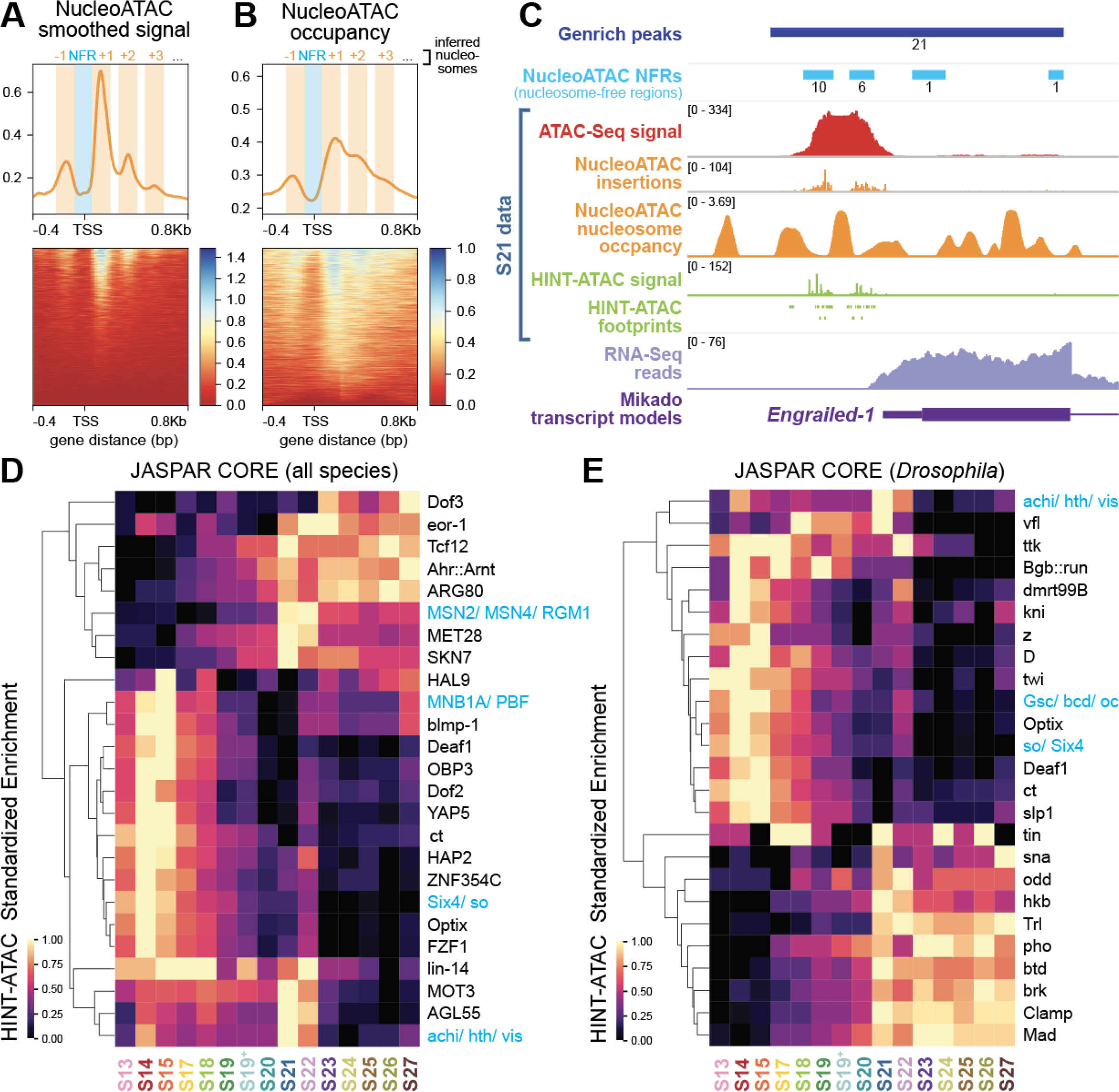
Inference of nucleosome positioning and transcription factor binding from Omni-ATAC-Seq data. A) NucleoATAC smoothed signal at mRNA starts genome-wide. A clear pattern of peaks in the signal can be observed, suggestive of nucleosome positioning. Axis reflects mean signal across all mRNA starts at stage S21. B) Inferred nucleosome occupancy at mRNA starts genome-wide. Axis reflects mean signal across all mRNA starts at stage S21. Inferred nucleosome annotations on A and B added based on observed signal. C) Visualization of NucleoATAC, HINT-ATAC, Omni-ATAC, and RNA-Seq data at the inferred Engrailed-1 promoter at stage S21. Deeper inference on Omni-ATAC-Seq data allows insight into the broader chromatin landscape at developmental time points included in this dataset. D) Summary of standardized enrichment of JASPAR CORE position frequency matrices (PFMs) among HINT-ATAC identified footprints across developmental stages. The top 25 most-enriched PFMs are displayed. Color is based on standardized enrichment, where minimum and maximum enrichment within each row are set to 0 and 1, respectively. PFMs marked in blue had identical enrichment ratios within each group at all timepoints, likely due to having highly similar PFMs. Overall, PFMs appeared enriched at early, late, or middle developmental stages. E) Summary of top 25 most-enriched *Drosophila* PFMs from the JASPAR CORE database. Overall, PFMs appeared enriched at early, late, or middle developmental stages.

A well-established signal of nucleosome positioning found across animals is the presence of strong +1 and -1 nucleosome positioning around gene promoters, and a depletion of nucleosomes at RNA-Pol II binding sites (referred to as nucleosome-free region, or NFR) (Brogaard et al., 2012; Radman-Livaja and Rando, 2010). We visualized NucleoATAC signal and nucleosome occupancy around mRNA starts genome-wide, and observed strong signals associated with +1 and -1 nucleosomes (Fig. 4 A, B; Supp. Fig. 4 B–D). We also observed a depletion of nucleosomes and a decrease in NucleoATAC signal between the +1 and -1 nucleosomes, which may represent nucleosome-free regions. This signal was observed consistently in each of our stage-specific libraries (Supp. Fig. 4 E, F). These data indicate that we were able to identify the stereotypical signal of nucleosomes at promoters from our Omni-ATAC data. Moreover, these results suggest that NucleoATAC is more generally able to infer nucleosome positions and nucleosome-free regions within our dataset genome-wide.

In addition to inferring nucleosome positions, deeply sequenced ATAC-Seq data also enables the inference of transcription factor footprints. We used HINT-ATAC to identify transcription factor footprints in Omni-ATAC-Seq peaks from each developmental stage (Li et al., 2019) and evaluated the enrichment of these TF footprints across developmental time. We observed that HINT-ATAC enriched TF footprints appeared to group into three broad clusters based on the timing of highest enrichment: early, S21/S22, and late (Fig. 4D, E). These trends were true both when examining all JASPAR CORE position frequency matrices (PFMs) (Fornes et al., 2020), and when examining only those JASPAR CORE PFMs from *Drosophila*. These results are consistent with the general trends observed in PCA, which suggest that the early, middle, and late timepoints in development have the greatest variation.

The data generated from these analyses enable the visualization of chromatin accessibility, predicted nucleosome position, inferred transcription factor binding, and RNA-Seq expression at individual loci across around half of *Parhyale* development. The strengths of these data are illustrated in Fig 4C at the predicted promoter of the *Parhyale Engrailed-1* locus for stage S21. Thus, for any genomic region of interest, one can develop a comprehensive prediction of the local chromatin environment at each of the 15 timepoints in our dataset.

### Identifying gene regulatory programs from Omni-ATAC-Seq using fuzzy clustering

To assess the potential for our dataset to provide new biological insights, we attempted to identify distinct gene regulatory programs based on differential accessibility. We used the Mfuzz package to perform fuzzy c-means clustering on the matrix of read counts generated from our Omni-ATAC-Seq peaks (Kumar and E Futschik, 2007). Fuzzy clustering differs from categorical clustering approaches, such as hierarchical clustering and k-means clustering, in that it assigns a probability for each element in the dataset to fall into each of the identified clusters. For example, rather than a given peak falling into either cluster A or B exclusively, a given peak might have a 95% chance of falling into cluster A and a 5% chance of calling into cluster B.

The Mfuzz package allows the specification of two parameters: c, or the number of clusters to create from the data, and m, the stringency of the clustering. To determine the optimal number of clusters in our fuzzy c-means clustering, we varied the c-value from 3 to 13, and evaluated the quality of clustering using the cluster overlap metric provided by Mfuzz, as well as the silhouette score metric (Supp. Fig. 5.2)(Kumar and E Futschik, 2007). We determined that 9 clusters yielded the highest mean silhouette score, while avoiding clusters below the silhouette score average and minimizing size variability between clusters. Figure 5A plots with t-SNE the top 10,000 most significant peaks called by IDE2, where each point is colored by its IDE2 model max fit, while Figure 5B illustrates the same clusters, colored by their Mfuzz cluster.

**Fig. 5:**
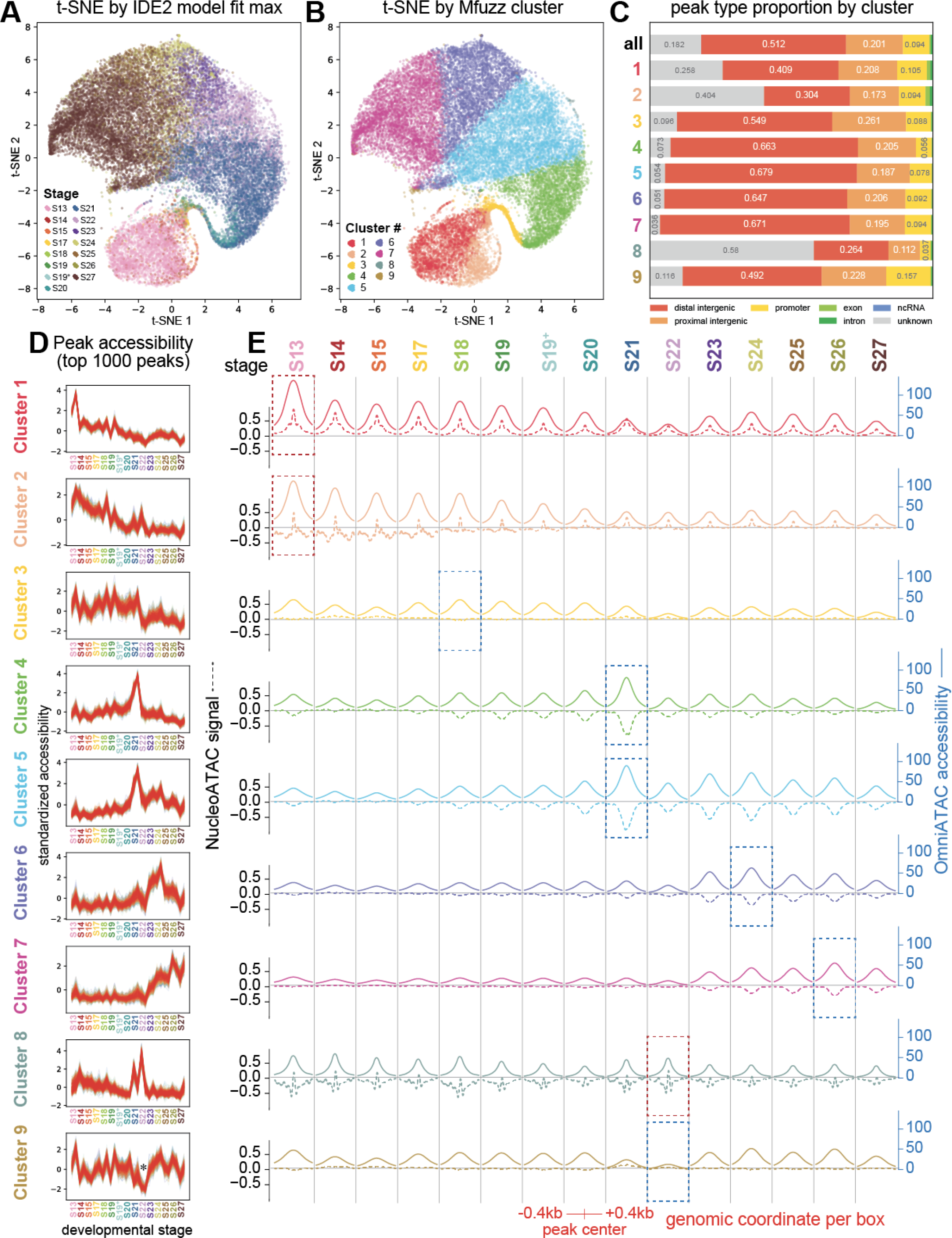
Identification and classification of regulatory element clusters. A) t-SNE plot of the top 10,000 most statistically significant peaks identified by ImpulseDE2 (IDE2), colored by IDE2 max fit. B) t-SNE plot of the same points from A colored by Mfuzz cluster. C) Distribution of peak position categories by Mfuzz cluster. Clusters, 1, 2, and 8 appeared to be enriched for “unknown” peaks, while other clusters appeared to be enriched for distal intergenic peaks. D) Standardized accessibility plot for top 1000 peaks with strongest membership in each of the 9 clusters. Each library is plotted as a separate point along the line plot. Line color indicates the cluster membership value for each peak in each plot, with dark red indicating strong cluster membership, and blue indicating weaker cluster membership. E) Line plots showing Omni-ATAC accessibility (solid line, blue axis) and NucleoATAC signal (dashed line, black axis) for peaks in each cluster across time. Each box contains a line plot summarizing each signal at all peaks within that cluster with respect to a given developmental Omni-ATAC-Seq library. The center of each line plot reflects the average signal at the center of all Omni-ATAC-Seq peaks in that cluster at that developmental stage. Signal is visualized at 0.4kb upstream and downstream of peak centers. Within each signal type, the axes across line plots for each cluster are identical. Dotted lines around subsections of the plot highlight regions of interest. Dotted blue lines indicate the time point during which peaks in a given cluster achieved both maximum accessibility and decreased nucleosome occupancy. Dotted red lines indicate the time point during which peaks in a given cluster achieved both maximum accessibility and increased nucleosome occupancy.

We evaluated the differences between clusters through a variety of approaches. Figure 5D shows the standardized raw accessibility scores of peaks in each of the 9 clusters over time. We first assessed the spatial category distribution of each cluster compared to the global average (Figure 5C). Clusters 1, 2, and 8 appeared to be enriched for “unknown” regions, while clusters 4–7 and 9 appeared to be enriched for distal intergenic regions relative to the global distribution. In addition, we used the IDE2 model fits for each cluster to evaluate the general accessibility dynamics of each cluster (Supp. Fig. 5.1B, C). Clusters 1 and 2 appear to be enriched for peaks that achieve high accessibility early in development and decrease in accessibility over time. Clusters 3–6 appear to be enriched for peaks that achieve maximum accessibility at different timepoints along developmental time. Cluster 7 appears to be enriched for peaks that show low accessibility early in development, and increased accessibility late in development. Finally, clusters 8 and 9 showed a more striking pattern, with Cluster 8 composed of peaks that appear to have a pulsatile increase in accessibility around stages S21–S22, and Cluster 9 composed of peaks with an opposite pattern: those with a sudden pulsatile decrease around S21–S22. Thus, the 9 clusters identified in our analysis appeared to capture distinct patterns of accessibility throughout time.

We further investigated the clusters identified by Mfuzz using additional metrics in order to understand the biological significance of these clusters. First, we investigated the relationship between accessibility dynamics and nucleosome positioning between clusters (Fig. 5E, Supp. Fig. 5.1D). For each developmental time point, we visualized Omni-ATAC accessibility (solid line, black axis) compared to NucleoATAC signal (dashed line, blue axis) for peaks belonging to each cluster. We observed that peaks belonging to Clusters 3–7 and 9 all appeared to have an inverse relationship between accessibility and NucleoATAC signal; this inverse relationship was most obvious at the time points during which these peaks achieved their maximum accessibility (Fig. 5E, blue dashed boxes). Moreover, these peaks all appeared to have a depletion of nucleosome occupancy at peak centers relative to the region around peaks (Supp. Fig. 5.1D, solid line, pink axis). By contrast, peaks in clusters 1, 2, and 8 appeared to show a counterintuitive and opposite result: both high NucleoATAC signal and high nucleosome occupancy at timepoints during which those peaks achieved maximum accessibility (Fig. 5E, Supp. Fig. 5.1D, solid line, pink axis). As with the clusters that showed an inverse relationship between accessibility and occupancy, the clusters that showed a parallel relationship between accessibility and occupancy showed the strongest relationship at those timepoints during which they achieved maximum accessibility (Fig. 5E, red dashed boxes).

To assess the biological function of each of the 9 Mfuzz clusters, we performed GO term enrichment analysis. To assign functions to each peak, we extracted the *Drosophila melanogaster* gene assigned to the nearest gene for each peak, and performed GO enrichment analysis on the list of unique gene names associated with peaks from each cluster relative to the list of all OrthoFinder-assigned gene names in the genome. Fig. 6A illustrates the top 10 most-enriched GO terms for each of the clusters.

**Fig. 6:**
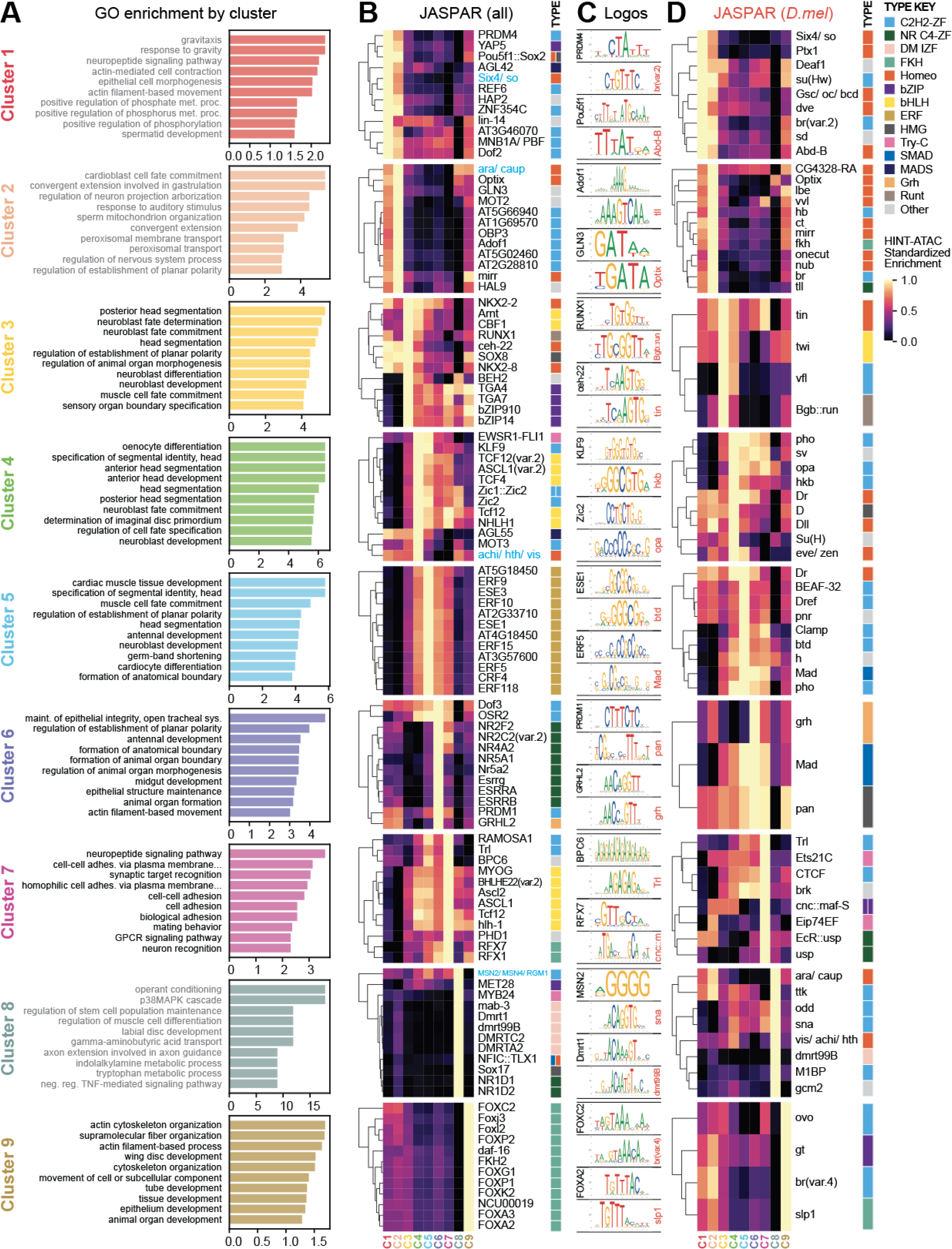
Functional annotation of clusters using GO enrichment and transcription factor binding prediction. A) GO enrichment per cluster based on the nearest gene for each peak. For each peak in each cluster, we identified the nearest gene and extracted the *Drosophila* OrthoFinder gene name for that gene. We performed GO enrichment using *Drosophila* GO terms, using the list of all OrthoFinder gene names found in our genome as the background for the analysis. Bar charts display fold enrichment for each GO term. Gray text represents GO terms that did not have a significant p-value (p < 0.05) after false discovery rate (FDR) correction. B) HINT-ATAC enrichment per cluster for JASPAR CORE PFMs. For each cluster, we identified all unique TF footprints across all 15 timepoints found within peaks of that cluster and performed enrichment relative to a randomly-selected background. We show the top 12 PFMs with the greatest enrichment for each cluster that also had the highest enrichment in that cluster relative to other clusters. Colored boxes represent the transcription factor family of each transcription factor in the clustermap. C) JASPAR motif logos for select pairs of transcription factors between non-*Drosophila* and *Drosophila* TFs with similar motifs as determined by STAMP alignment. For each cluster, two pairs of transcription factors are shown: a non-*Drosophila* transcription factor (labeled in black) and a *Drosophila* transcription factor (labeled in black). In some cases, a strong similarity is observed between the non-*Drosophila* and *Drosophila* PWMs. D) HINT-ATAC enrichment per cluster for *Drosophila* JASPAR CORE PFMs. Some clusters had fewer than 12 PFMs for which that cluster had the highest enrichment.

While we were not able to identify significantly enriched GO-terms (FDR < 0.05) for Clusters 1, 2, and 8, Clusters 3–7 and 9 all showed enrichment of developmental GO-terms. These enriched GO-terms generally appeared to match the expected biological functions of peaks based on their accessibility dynamics. For example, Cluster 3 peaks, which generally achieve maximum accessibility during germ band elongation stages in development, showed enrichment for GO terms including head segmentation and neuroblast fate, two processes which take place during this developmental period. Meanwhile, Cluster 7 peaks appeared to be enriched for neuronal terms (neuropeptide signaling pathway, synaptic target recognition, neuron recognition), potentially reflecting the function of genes near these peaks which achieve their maximum accessibility towards the very end of the developmental time-course, during which much of the morphology of the organism has been specified, and the embryo begins to twitch inside the egg.

To further investigate the clusters in our dataset, we examined the enrichment of TF footprints generated from HINT-ATAC in each of our clusters. For each cluster, we compiled all of the TF footprints predicted across developmental stages for all peaks within that cluster. We compared the enrichment of those TF footprints to our randomly generated background data, and examined all JASPAR CORE transcription factors, as well as the subset of JASPAR CORE transcription factors found in *Drosophila*.

We observed strong differential enrichment of different groups of transcription factors in each of our 9 clusters. Notably, particular families of transcription factors appeared enriched in several of the clusters. For example, Cluster 1 and Cluster 2 appeared to have enrichment for footprints matching C2H2 zinc-finger transcription factors and homeodomain transcription factors. Clusters 4 and 7 showed enrichment for footprints matching beta helix-loop-helix (b-HLH) transcription factors. Meanwhile, clusters 5, 8, and 9 showed strong enrichment for footprints of ethylene response factor (ERF), DM-type intertwined zinc-finger factor (DM IZF), and forkhead domain (FKH) transcription factor families, respectively.

Given that many JASPAR PFMs come from non-arthropod sources, we attempted to identify possible *Drosophila* motif matches for the enriched transcription factor footprints found in the JASPAR CORE dataset by comparing the most-enriched JASPAR CORE and JASPAR *Drosophila* PFMs. We used the STAMP tool through the JASPAR webpage to align motif sequences based on similarity (Mahony and Benos, 2007). We identified groups of aligned PFMs between non-*Drosophila* and *Drosophila* PFMs (see Supp. Fig. 6). Selected pairs of motifs are shown in Fig. 6C for each cluster (non-*Drosophila* labeled in black, *Drosophila* labeled in red). For some pairs, we were able to identify candidate *Drosophila* transcription factors that appeared highly similar to non-*Drosophila* sequences. For example, in Cluster 9, we observed strong enrichment for FKH family transcription factors, whose PFMs matched the br(var.4) and slp1 PFMs. This suggests that peaks found in Cluster 9 may be regulated by br(var.4) and slp1 binding in *Parhyale*.

For other clusters, we had variable success. For example, we were unable to identify a strong *Drosophila* match to explain the strong enrichment of ERF transcription factors (found in *Arabidopsis thaliana*) in Cluster 5. Nonetheless, footprints matching ERF transcription factors appeared to be highly enriched in Cluster 5, suggesting that there may be a transcription factor within the *Parhyale* genome whose PFM matches that of ERF transcription factors, and which may be important for the regulation of Cluster 5 peaks.

Altogether, our results indicate that clustering Omni-ATAC data using accessibility can identify groups of transcription factors with similar accessibility trajectories. In addition, these clusters can be further analyzed using nucleosome occupancy, GO enrichment, and transcription factor footprint enrichment to understand possible biological functions and genetic mechanisms behind differential accessibility.

### Concordant and discordant expression and accessibility dynamics appear across development

To understand the relationship between peak accessibility and gene expression, we examined the correlation between Omni-ATAC accessibility in individual peaks with respect to RNA-Seq expression for the nearest gene at each of the developmental stages for which we generated RNA-Seq data. We first examined the relationship between accessibility at all peaks compared to their nearest genes (Supp. Fig. 7A) and observed a very weak (Pearson R^2^ = 0.11–0.15, Spearman ρ = 0.1–0.14) but statistically significant (p < 0.001) positive correlation between peak accessibility and the expression of nearby genes. When we examined only promoter peaks (Fig. 7B), we observed a slightly greater correlation between accessibility and gene expression (Pearson R^2^ = 0.19–0.22, Spearman ρ = 0.17–0.25) correlation that was also significant (p < 0.001). Notably, at each developmental stage, we observed a higher correlation between accessibility and expression at promoter peaks than at all peaks.

**Fig. 7:**
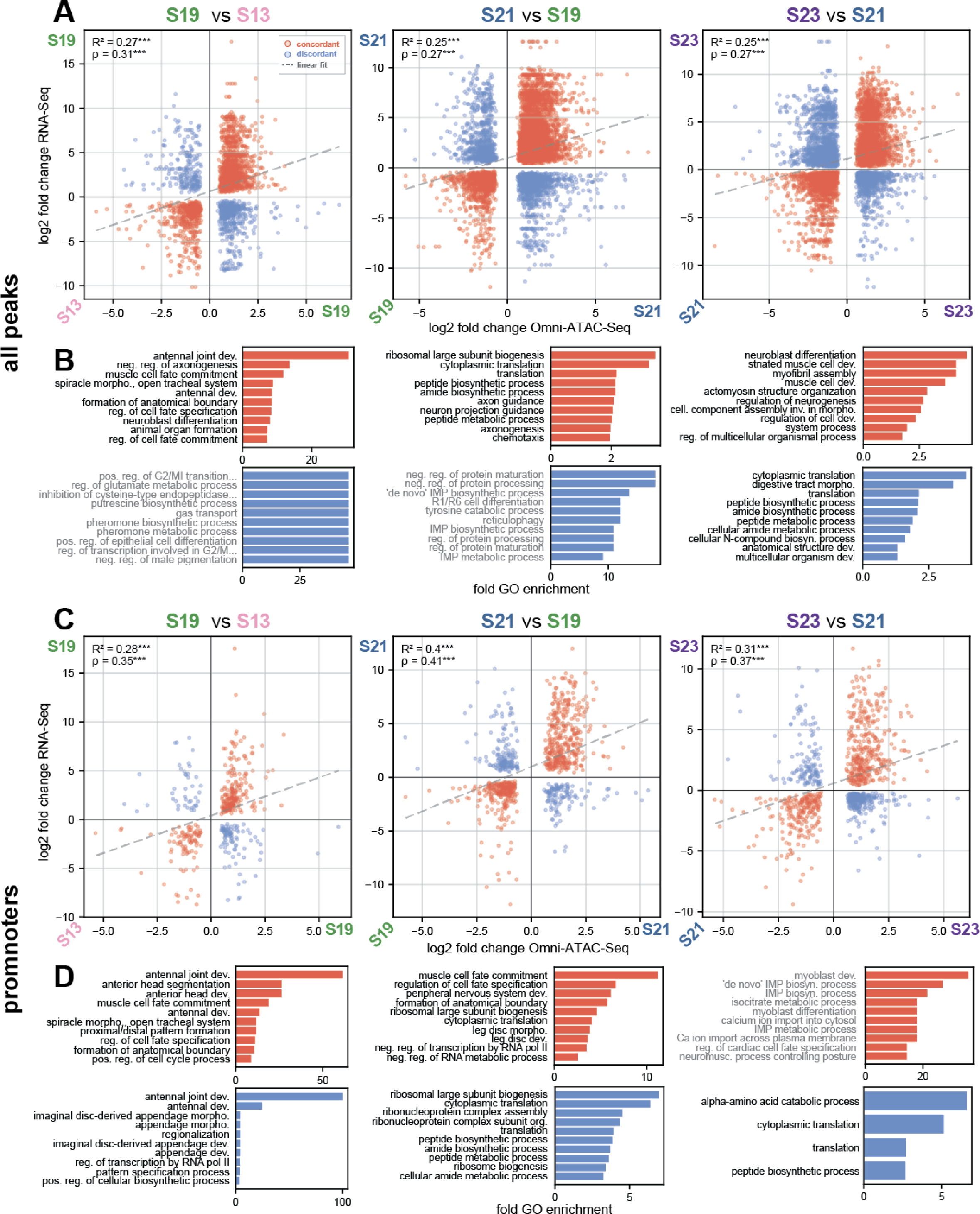
Correlation between RNA-Seq and Omni-ATAC-Seq. A) Correlation between log_2_-fold change RNA-Seq and log_2_-fold change Omni-ATAC-Seq at peak-gene pairs in sequential pairwise comparisons of developmental stages. Only peak-gene pairs with a significant difference in expression and accessibility between the two time points are plotted. Positive axis reflects higher expression or accessibility at the later developmental stage for each plot. Points colored in red reflect concordant relationships between log_2_-fold change in RNA-Seq and Omni-ATAC-Seq; blue peaks reflect discordant relationships. Dotted line represents a linear fit to all data in the plot. Pearson correlation R^2^ and Spearman correlation ρ for all points are displayed in each plot. B) GO-term enrichment for gene-peak pairs with concordant or discordant expression and accessibility log_2_-fold change, for all peaks. Gene lists were extracted based on the nearest Mikado gene to each peak. C) Correlation between log_2_-fold change RNA-Seq and log_2_-fold change Omni-ATAC-Seq at all peaks in each developmental stage. Positive axis reflects higher expression at the later developmental stage for each plot. D) GO-term enrichment for gene-peak pairs with concordant or discordant expression and accessibility log_2-_fold change, for promoter peaks. (*** = p < 0.001)

We expect that both enhancers and repressors might be found among our ATAC-Seq peaks. To further classify peaks, we examined the relationship between the change in accessibility of individual peaks and the change in expression of nearby genes. We used DESeq2 to evaluate the log_2_-fold change of both expression (RNA-Seq) and accessibility (ATAC-Seq) for all genes and peaks. We then correlated the differential accessibility and gene expression for all peak-gene pairs where both the differential accessibility of a peak and expression of the peak’s nearest gene were statistically significant (FDR/ padj < 0.05). When examining all peaks, we observed a weak but significant correlation between log_2_-fold change in accessibility and gene expression (Pearson R^2^ = 0.25–0.27, Spearman ρ = 0.27–0.31)(Fig. 7A). This correlation was slightly stronger at promoter peaks (Pearson R^2^ = 0.28–0.40, Spearman ρ = 0.35–0.41)(Fig. 7C).

Among our differential accessibility and expression analyses, we classified two broad classes of peak-gene pairs representing concordant and discordant accessibility and expression dynamics. Peaks with concordant accessibility and gene expression were those where the sign of the log_2_-fold change in expression and accessibility were in agreement—for example, peaks where an increase in accessibility was observed concurrently with an increase in expression. Meanwhile, peaks with discordant accessibility and gene expression were those where an increase in accessibility was observed concurrently with a decrease in expression, or vice versa.

We observed both classes of peak-gene pairs among all peaks, but also among promoter peaks.

Over time, the number of concordant peak-gene pairs increased from 1,888 at the S19-vs-S13 comparison, to 5,738 at the S21-vs-S19 comparison, and then decreased to 5,240 at the S23-vs-S21 comparison (Supp Fig. 7D). Meanwhile, the number of discordant peaks gradually increased over time (761 to 2,231 to 3,115 peak-gene pairs)(Supp Fig. 7D). These trends were also observed for the promoter-only peaks. The gradual increase in discordant peaks may indicate an increase in repressive gene regulatory mechanisms over time, as developmental gene networks become refined over the course of differentiation.

Overall, the classification of peaks as concordant or discordant in accessibility and gene expression may provide insight both into the function of individual peaks as enhancers or silencers, and the presence of repressive regulation within promoter peaks. These data provide additional insights into the relationship between accessibility and gene expression during development in this organism.

### Omni-ATAC-Seq re-identifies known *Parhyale* regulatory elements

To assess the practical usefulness of our Omni-ATAC-Seq dataset, we compared our Omni-ATAC-Seq peaks to known regulatory elements in *Parhyale*. A very limited set of regulatory elements have been described in *Parhyale*: a muscle reporter, PhMS (Pavlopoulos and Averof, 2005); a heat shock element, HS2a (Pavlopoulos et al., 2009); an embryonic ubiquitous reporter, PEB (Danielle Liubicich, unpublished); and two reporter constructs for *Parhyale Opsin-1* and *Opsin-2* (Ramos et al., 2019). We were unable to locate any Omni-ATAC-Seq peaks for the *Parhyale Opsin* genes, likely owing to their late expression (outside of our developmental time course) and very low cell number (a handful of cells per eye). However, strong Omni-ATAC-Seq enrichment was observed at the PhMS, HS2a, and PEB elements, as illustrated in Supp. Fig. 8.1.

**Fig. 8:**
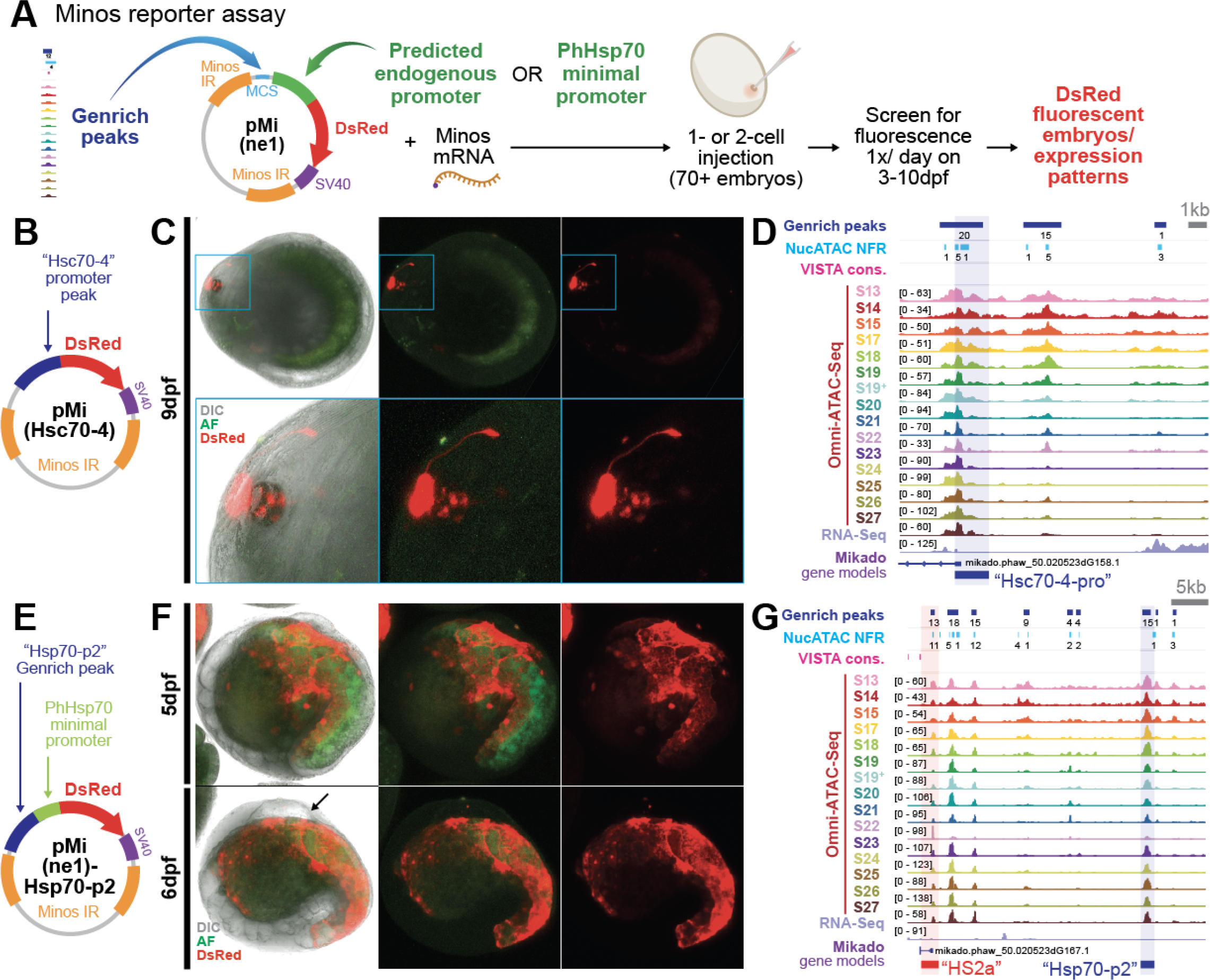
Minos transposase reporter assay identification of novel regulatory elements. A) Minos transposase reporter assay and screening approach. B) Plasmid schematic for the pMi(Hsc70-4) reporter. C) Expression of the pMi(Hsc70-4) reporter in neurons associated with the eye of a 9hpf embryo. Bottom row shows a zoomed-in image of the head region. A clear projection of the neurons can be seen. D) The genomic region of the Hsc70-4 promoter peak. E) Plasmid schematic for the pMi(ne1)-Hsp70-p2 reporter. F) Expression of pMi(ne1)-Hsp70-p2 reporter at 5 and 6dpf showing strong expression in the yolk. Arrow marks aberrant morphology at the dorsal side of the embryo observed consistently in all embryos with fluorescent expression. G) The genomic region of the Hsp70-p2 peak relative to the HS2a element. The PhHsp70 minimal promoter used in this construct is located within the HS2a element.

Each of these reporter constructs consists of the first exon and intron of a gene. In each case, we observed an Omni-ATAC peak overlapping the 5’ end of the gene. For the PhMS reporter (Supp. Fig. 8.1B), previous work had identified a cluster of putative b-HLH transcription factor binding sites. These sites were captured within our Omni-ATAC peak. The HS2a reporter has been reported to contain two binding sites for heat shock factor (HSF) proteins upstream of a minimal promoter (PhHsp70), both of which are captured within our Omni-ATAC peak (Supp. Fig. 8.1D). Finally, the PEB element appears to contain two Omni-ATAC peaks. These results suggest Omni-ATAC is able to identify the position of functional cis-regulatory elements, and has been able to capture those regions important for cis-regulatory element function.

### *Minos* transposase reporter assays can reveal novel promoters and distal enhancers

Given that our Omni-ATAC peaks were able to capture known regulatory elements, we attempted to use these peaks to identify novel regulatory elements. Careful examination of developmentally important genes revealed that many are surrounded by large numbers of peaks (>10) spread over large genomic distances. To further filter our Omni-ATAC peaks, we identified regions of sequence conservation to another amphipod crustacean, *Hyalella azteca* (Poynton et al., 2018), using the VISTA sequence alignment software (Ratnere and Dubchak, 2009)(Supp. Fig. 8.2A). *Hyalella* serves as a useful comparison to *Parhyale*, as its genome size is smaller (1.05 Gb), but as it is also an amphipod crustacean, we expect that key developmental regulatory elements might have some level of sequence conservation.

Among our Omni-ATAC peaks, we took two strategies to identify candidate reporter elements. In the first approach, we identified all peaks within 5kb of mRNA starts. In doing so, we were able to identify a handful of genomic regions in which we were able to locate both a putative promoter and a candidate proximal enhancer. We also identified several putative promoters, which we tested in isolation. In the second approach, we examined the genomic regions around important developmental genes of interest. Many of the genes we examined showed very large numbers of strong peaks, and were therefore intractable to thorough analysis. For the purposes of this study, we focused on three regions: the region around the *Engrailed-1* gene, the region around the *Sp-69* gene, and the region around the *Heat shock protein 70* complex, where two previous cis-regulatory elements (PhMS and HS2a) had been identified.

To assess the function of candidate regulatory elements identified by Omni-ATAC-Seq, we employed a *Minos* transposase reporter assay (Fig. 8A). In this assay, we injected *Minos* transposase mRNA along with a transposon donor plasmid containing a reporter gene construct that expressed DsRed to 1- and 2-cell embryos. Once per day, from 3–10dpf, we screened for DsRed expression on a Zeiss LSM780 confocal microscope.

The reporters we examined are summarized in Table 1. In the course of our experiments, we observed numerous spontaneous expression patterns (Supp. Fig. 9). These expression patterns may represent rare integration events, or may reflect fortuitous insertions into regions of the genome with promiscuous regulatory sequences that facilitate expression.

**Table 1:** Plasmids tested using Minos transgenesis.

| Plasmid | Type | Gene – *Drosophila* expression | # Inj. | # Surv. | # DsRed+ | Phenotype |
| --- | --- | --- | --- | --- | --- | --- |
| pMi(gm) | promoter | N/A – *Parhyale* muscle reporter PhMS | 109 | 39 | 24 | Green muscle expression |
| pMi(retn) | promoter | *retained* – expressed in some neurons | 240 | 104 | 5 | No consistent phenotype |
| pMi(SVP) | promoter | *seven up* – expressed in ventral neuroblasts | 122 | 31 | 0 | None |
| pMi(su-s-2) | pro + peak | *suppressor of sable* – embryonic ubiquitous expression | 127 | 57 | 7 | No consistent phenotype |
| pMi(Rx) | pro + peak | *retinal homeobox* – expressed in various neural tissues | 74 | 28 | 0 | None |
| pMi(Hsc70-4) | promoter | *Heat shock 70 complex protein 4* – embryonic ubiquitous | 212 | 99 | 22 | Neurons associated with the eye |
| pMi(En1) | promoter | *engrailed* – parasegment stripes | 71 | 33 | 0 | None |
| pMi(En1)-p1 | pro + peak | “ | 67 | 46 | 0 | None |
| pMi(En1)-p1.5 | pro + peak | “ | 70 | 51 | 0 | None |
| pMi(En1)-p5 | pro + peak | “ | 68 | 58 | 1 | Faint expression domain |
| pMi(Sp69) | promoter | *Sp69* – leg gap gene | 74 | 56 | 0 | None |
| pMi(Sp69)-p1 | pro + peak | “ | 50 | 30 | 0 | None |
| pMi(Sp69)-p1.5 | pro + peak | “ | 86 | 32 | 3 | No consistent phenotype |
| pMi(Sp69)-p2.5 | pro + peak | “ | 71 | 35 | 0 | None |
| pMi(Sp69)-p3 | pro + peak | “ | 241 | 121 | 8 | Head, leg, gill |
| pMi(ne1)-Hsp70-p1 | pro + peak | *Hsp70 complex* | 45 | 26 | 0 | None |
| pMi(ne1)-Hsp70-p2 | pro + peak | *Hsp70 complex* | 310 | 81 | 61 | Yolk cells |

One expression pattern observed from the Sp69-p3 enhancer appeared to be repeated in 5 embryos along the ∼300 we injected. This expression pattern was characterized by strong expression in the head, antenna 1, and antenna 2 (Supp. Fig. 10). In the same set of experiments, we also observed three spontaneous expression patterns that appeared to differ amongst each other, labeling different regions of the limbs and gills. It is possible that the repeated expression observed from the Sp69-p3 enhancer represents a genuine expression pattern. We tested this hypothesis by raising three Sp69-p3 head-expressing animals to adulthood and mating the two adults that showed the strongest expression during development. However, none of the offspring of this cross showed obvious head expression, or any obvious DsRed expression elsewhere in the embryo.

Among the novel reporter constructs we tested, two showed robust expression: pMi(Hsc70-4) and pMi(ne1)-Hsp70-p2. The pMi(Hsc70-4) construct was constructed using the putative promoter peak of the Hsc70-4 gene (Fig. 8B, D). This reporter showed strong expression in a cluster of neurons associated with the eye beginning at around 9dpf (Fig. 8C). The expression pattern of these neurons appears to be distinct from those observed using a synthetic 3XP3 enhancer, which has previously been used as a positive marker of transformation *Parhyale* (Pavlopoulos and Averof, 2005). Thus, it is possible to build new reporters from our Omni-ATAC peaks using putative promoter peaks.

The second reporter that showed robust expression was the pMi(ne1)-Hsp70-p2 construct, which contains the PhHsp70 minimal promoter along with a strong peak extracted from the Hsp70 cluster, about 30kb away from the location of the minimal promoter (Fig. 8E, G). Hsp70-p2 drives strong expression in cells of the embryonic yolk (Fig. 8F). Some yolk cells labeled by this reporter moved dynamically around the yolk, and while others appeared static and showed spongiform morphology (Supp. Video 1). We observed defects in the development of the dorsal portion of the embryo in mid-stage (∼6–7dpf) embryos injected with this construct, which stereotypically displayed a separation between the embryo proper and the eggshell (Fig. 8F, bottom panel, marked with arrow). While we were unable to establish a genetic line using this construct, we observed strong and reproducible expression in many embryos. This reporter is the first indication that distal enhancers exist and can be identified in *Parhyale*.

While it remains challenging to identify novel cis-regulatory elements in *Parhyale*, our results suggest that functional regulatory elements can be identified from our Omni-ATAC data. Among the numerous peaks identified in our dataset, we expect that many will reveal novel gene regulatory dynamics. Future work to identify such elements will require the optimization of reporter expression and transgenic strategies to increase the throughput and sensitivity of reporter gene assays.

## Discussion

New genomic data enable deeper understanding of underlying biology. This is especially true in emerging research organisms, for which data and resources are limited. This work provides a wealth of genomic resources for the amphipod crustacean *Parhyale hawaiensis*. In addition to providing Omni-ATAC-Seq and RNA-Seq data for a broad developmental time course, our work generates multiple new transcriptomes, an updated genome annotation, a catalog of dynamically accessible chromatin regions, and predictions of nucleosome occupancy and transcription factor binding. Moreover, our thorough analysis of regulatory element dynamics through clustering, GO enrichment, predicted TF binding, and correlations with RNA-Seq present numerous hypotheses for cis-regulatory element function. Such data will support researchers in the growing *Parhyale* community in efforts to identify and characterize developmental regulatory elements, and provide the foundations for more advanced approaches, including single-cell sequencing techniques, which rely on high-quality reference datasets.

### The *Parhyale* genome contains many distant and dynamic regulatory elements

From our data, we were able to glean new information about the global dynamics of gene regulation during *Parhyale* development, as well as the composition of the *Parhyale* genome. We observed that the vast majority of our peaks showed dynamic accessibility over developmental time, and that peaks could have a variety of different temporal dynamics, including transient increases and decreases in accessibility alongside more absolute increases and decreases. These results indicate that much of the *Parhyale* genome undergoes dynamic changes in accessibility over developmental time.

With an improved genome annotation, we were also able to determine that the vast majority (>70%) of regulatory elements in *Parhyale* are located in intergenic regions, and that among all elements, a majority (51%) were located greater than 10kb away from the nearest gene. These results are notable, given that many of the currently studied protostome genomes are small (for example, *Drosophila melanogaster* is 180Mb, *Tribolium castaneum* is 200Mb, and *Bombyx mori* is 530Mb). In such organisms, regulatory elements tend to occur within short distances from promoters, and it is common in such systems to attempt to build reporter genes using a short window upstream of the promoter of a gene of interest.

*Parhyale* is an example of an arthropod with a large genome (3.6 Gb, or ∼10% larger than *Homo sapiens* at 3.2 Gb). Given the large genome size, and the large proportion of elements located distant to genes, we expect that regulatory elements within *Parhyale* may be located more distantly from gene promoters than in other arthropods. Future attempts at building reporter genes in this organism will need to account for the large number of distant regulatory elements, which may be critical to proper reporter expression.

Our data are among the first to identify regulatory elements genome-wide in a non-insect arthropod (Gatzmann et al., 2018; Kissane et al., 2021), and also examine one of the largest sequenced arthropod genomes available to date (Kao et al., 2016). These data will enable other researchers to examine the relationship between genome size and regulatory element composition across more diverse taxa, and will be useful in developing a deeper understanding of how genome organization affects gene expression across the metazoans.

### Combined short- and long-read transcriptomes can improve genome annotations

In addition to providing a resource for the *Parhyale* community, our work can also serve as a guide for other researchers working on developing such datasets in emerging research organisms. First, our work demonstrates the utility of combining short- and long-read sequencing in generating more accurate genome annotations for emerging research organisms. Even with a small number of sequences (∼1.2 million reads) generated using Nanopore sequencing, we were able to assemble a transcriptome with a moderate BUSCO score (71%). Combining short reads and long reads (StringTie2 SL), we were able to construct a transcriptome with both a high BUSCO score (91.8%) and low rate of gene fragmentation.

Moreover, the completeness of the StringTie2 SL transcriptome met or exceeded that of several of our Trinity-derived transcriptomes. Examination of individual genomic regions in the *Parhyale* genome indicates that Trinity generates numerous spurious transcripts, which can be confounding for whole-genome analysis (see Supp. Fig. 3.2 for examples). Thus, for other researchers with access to a genome assembly, we would strongly recommend performing both short- and long-read sequencing, and assembling using multiple different assembly strategies to generate higher quality genome annotations. For those without access to a high-quality genome, we would caution against relying strictly on transcripts generated from software such as Trinity, which in our system generated many transcripts that did not appear to match with previous RACE data. Filtering transcripts by expression values, as well as employing transcriptome-merging pipelines such as Mikado (in the case of a high-quality genome) and EvidentialGene (in the absence of a genome), would likely help remove such spurious transcripts.

### Deep analysis of ATAC-Seq data can reveal more than just dynamic accessibility

While the most direct analysis of ATAC-Seq data can enable the identification of dynamic regulatory elements, our work uses additional tools to infer nucleosome positioning and transcription factor binding from our ATAC-Seq data. Using NucleoATAC, we were able to recover clear signals of nucleosome positioning at promoters, a quality observed in numerous organisms. Using HINT-ATAC, we were able to predict transcription factor binding across developmental stages. Such analyses enable a glimpse of the possible chromatin landscape of the genome, particularly in the absence of more direct and specific, but also more time-consuming and expensive approaches such as ChIP-Seq or CUT&RUN.

In addition to performing inference, we were able to identify clusters of peaks with potentially distinct biological functions using fuzzy clustering. These clusters of peaks appeared to differ in their accessibility dynamics, nucleosome positioning, and enrichment of transcription factor footprints, indicating that meaningful biological differences can be gleaned from deeper analysis of ATAC-Seq data. For example, we observed strong enrichment in Cluster 9 for FKH transcription factors footprints matching the PFMs for *Drosophila* br(var.4) and slp1. The strongest peaks in this cluster appeared to share a common decrease in accessibility during stages S21–S22, and an enrichment for genes with GO terms related to cytoskeletal function. This group of peaks appeared to show a decrease in accessibility at a time point during which the embryo undergoes a dramatic morphogenetic event in which it splits along the midline. Based on our data, it is possible that FKH domain-containing transcription factors may play a role in that important morphogenetic event. Using our analyses as a starting point, other *Parhyale* researchers may be able to make hypotheses about their own developmental processes of interest.

### Omni-ATAC-Seq identifies old and new regulatory elements, including enhancers

Our data were able to capture most of the previously-identified regulatory elements that had been described for *Parhyale*. In addition, we were able to demonstrate that our data contain new regulatory elements. Notably, we demonstrate the first identification of a distal regulatory element, Hsp70-p2, located about 30kb from the PhHsp70 minimal promoter. These results are the first identification of an enhancer separated by a large distance from a minimal promoter element in this organism.

Our results demonstrate that Omni-ATAC-Seq is able to identify novel regulatory elements, and we expect that numerous new reporter genes will be built from these data. Previous work relied on examining candidate regulatory elements within individual genomic regions, or attempting to build new reporters through random integration events. With this dataset, it is now possible to identify candidate regulatory elements for any gene of interest and to screen for expression patterns using the Minos reporter system. As this approach enables the direct investigation of candidate regulatory regions with known sequence, this will also enable the generation of novel synthetic reporters generated by modifying endogenous sequences. Such future directions will enable thorough dissection of regulatory elements to determine how transcription factor binding influences gene expression during *Parhyale* development.

While we were unable to identify regulatory elements that drove clear expression patterns for the handful of genes we tested, numerous additional peaks near these genes remain to be investigated.

Future work to build new reporters in *Parhyale* may require the development of more efficient transgenesis and screening strategies. For example, improving the efficiency of CRISPR-mediated homologous recombination or CRISPR-mediated NHEJ transgenesis could enable researchers to insert a reporter near a candidate regulatory region of interest. Moreover, future approaches will also need to account for the numerous distant regulatory elements in the *Parhyale* genome, which may be important for gene regulation. Examining local DNA interactions using approaches such as Hi-C or other chromatin conformation capture approaches will be instrumental in identifying distant regulatory regions.

Together, these approaches have enabled the exploration of chromatin dynamics and transcription factor binding in an emerging model organism. Our work illustrates how a single, easily adapted protocol can yield data amenable to deep analysis. We would recommend that other researchers using ATAC-Seq or similar assays in emerging model organisms also take advantage of the additional information that could be gleaned from deep analysis of their data, which could provide further insights into how both local and global changes to nucleosome occupancy and transcription factor binding influence their biological processes of interest.

## Conclusion

By combining Omni-ATAC-Seq with RNA-Seq across a broad developmental time course, our work is able to identify and classify numerous candidate cis-regulatory elements in the genome of the amphipod crustacean *Parhyale hawaiensis*. We demonstrate how deep analysis of Omni-ATAC data can facilitate the identification of peaks with distinct accessibility, nucleosome occupancy, and transcription factor footprint enrichment. We further classify peaks as concordant or discordant regulatory elements by integrating differential accessibility and differential expression, revealing candidate activating and repressive elements found in distal and proximal regulatory elements. Moreover, we show the potential to identify novel reporter genes using candidate promoters and enhancers from our data.

This work provides a substantial resource to the *Parhyale* community, and should accelerate the study of gene regulation in this emerging research organism. In addition, our work can serve as a framework for other researchers deploying ATAC-Seq and RNA-Seq approaches in emerging research organisms. Through deep analysis of ATAC-Seq data, combined short- and long-read sequencing, and integration of accessibility and gene expression, such researchers can identify regulatory elements with distinct biological functions and advance the study of gene regulation in their organisms of interest.

## Methods

### Crustacean Cultures

*Parhyale hawaiensis* were raised at 25°C and fed a diet of carrots, shrimp pellets, and Spirulina flakes in Tropic Marin artificial seawater with a salinity between 31–35ppm in plastic tanks.

### Embryo Cultures

Embryos were collected using previously described protocols (Rehm et al., 2009), staged using the Browne et al. 2003 staging guide, and raised at 27°C in a humidity-controlled incubator in filter-sterilized ASW (FASW). For ATAC-Seq and RNA-Seq experiments, clutches of 15 or more embryos were collected and staged between S1–S6, as these timepoints are among the shortest and most morphologically identifiable during early development.

### Embryo Staging

Two groups of five embryos each from a single clutch were used in the ATAC-Seq experiments, and three groups of five embryos each from a single clutch were used in the RNA-Seq experiments. Embryos of the correct stage were selected based on morphological characteristics as described in the Browne et al. staging guide, and any embryos with abnormal or asynchronous morphology were discarded.

Morphologically representative embryos from the same clutch as used for the ATAC-Seq were photographed on ventral and lateral positions within 10 minutes of beginning the tagmentation procedure. Any remaining embryos were boiled briefly and fixed using the protocol described in (Rehm et al., 2009) and DAPI stained to further confirm staging. The process of staging embryos, imaging embryos, boiling, and mashing embryos for Omni-ATAC tagmentation were all performed within a 30 minute time interval for each developmental stage.

### RNA Sequencing

Embryos for RNA isolation were homogenized in TriZol using a DWK Life Sciences (Kimble) Biomasher II Closed System Disposable Tissue Homogenizer, and RNA was isolated using the Zymo Direct-Zol Miniprep Plus kit. RNA quality was assessed based on fragment analysis using a Bioanalyzer. cDNA was generated from RNA using the TaKaRa SMART-Seq v4 Ultra Low-Input kit. cDNA was then sequenced using the Illumina NovaSeq to generate short reads. Using the Nanopore Direct cDNA barcoding kit (SQK-LSK109), cDNA was also sequenced on a Nanopore MinION flow cell to generate long reads.

### Limb development RNA Sequencing Methods

A pool of embryos consisting of embryonic stages 19, 20, 22, 23, 25, and 28 were homogenized with DWK Life Sciences Kimble Kontes Pellet Pestle Cordless Motor in Trizol and extracted using Trizol; PolyA+ libraries were prepared with the Truseq V1 kit (Illumina), starting with 0.6–3.5 mg of total mRNA, and sequenced on the Illumina HiSeq 2000 as paired-end 100 base reads, at the QB3 Vincent J. Coates Genomics Sequencing Laboratory.

### Limb development RNA-Seq *de novo* Transcriptome assembly

For the Trinity-limb (old) assembly, transcripts were assembled *de novo* using Trinity r2013_08_14 (Grabherr et al., 2011)with parameters --JM 170G --CPU 10 --inchworm_cpu 6 --min_kmer_cov 2 --min_contig_length 49 --group_pairs_distance 700. Input RNA-Seq reads were treated as paired-ends. Output transcript assemblies shorter than 200 bp were discarded. The remaining assemblies were screened for contaminants with BLASTX (BLAST+ v2.2.26; parameters: -num_descriptions 50 -num_alignments 50 -evalue 1e-5 -lcase_masking -soft_masking true -seg yes)(Camacho et al., 2009) against a database of all bacterial proteins downloaded from NCBI (retrieved 2013-08-31), then against Swiss-Prot UniProt human sequences (retrieved 2013-08-31)(The UniProt Consortium, 2019). All hits were further required to cover a minimum of 40% of each assembled transcript sequence. When filtering for human contaminants, a 98% identity threshold was also required. For the Trinity-limb (new) assembly, transcripts were assembled using Trinity v2.5.1 using standard settings.

### Transcriptome Assemblies

Short-read RNA-Seq data was used to generate both *de novo* and genome-guided transcriptomes using Trinity. For genome-guided assembly, reads were mapped using HISAT2. Long-read RNA-Seq data was mapped to the most recent *Parhyale* genome (phaw_5.0) using minimap2 and assembled using StringTie2. A combined transcriptome using both HISAT2-mapped short reads and minimap2-mapped long reads was also generated using StringTie2. See Supplementary Methods for additional information about assembly parameters.

### Mikado Transcriptome

Short-read Trinity transcriptomes and the transcriptome from the Kao et al. manuscript were mapped to the phaw_5.0 genome using GMAP. Short-read transcriptomes and long-read StringTie2 transcriptomes were merged using the Mikado software along with a previous genome annotation generated by Leo Blondel (using MAKER). See supplementary methods for additional information about Mikado parameters.

### Functional Annotation of Transcripts

The merged Mikado combined transcriptome was annotated using eggNOG and an Orthofinder comparison to *Drosophila*. The MAKER genome annotation was also annotated using eggNOG and OrthoFinder. See Supplementary Methods for additional information about functional annotation.

### ATAC-Seq Data

We performed conventional ATAC-Seq as per Buenrostro et al. 2013 for stages S13–S22. Data were used for benchmarking in comparison to Omni-ATAC data, but were not used for downstream analyses.

### Omni-ATAC-Seq Tagmentation

Tagmentation was performed using the reagents described in the Corces et al. 2017 Omni-ATAC-Seq paper, with the following modifications. Instead of the Illumina Nextera TD buffer, 2x Tagmentation Buffer from Wang et al. 2013 was used (Corces et al., 2017; Wang et al., 2013). Homemade Tn5 enzymes were purified and received as a gift from Jase Gehring and assembled into Tn5 transposomes as per Picelli et al 2014, using adapters purchased from IDT with 5’ phosphorylation (Picelli et al., 2014). Prior to adding RSB+D, embryos were washed 3x using 1X PBS. Embryos were mashed into a near-uniform solution using the tip of a low-retention p10 pipette, and the pipette was visually inspected for any remaining debris. We used the QIAGEN MinElute kit for purification and concentration of DNA.

### Omni-ATAC-Seq Library Preparation

Saturation PCR conditions for Omni-ATAC-Seq libraries were performed using a Roche Lightcycler 480 as per the Corces et al. 2017 paper. Optimal conditions for additional amplification after pre-amplification were determined based on the number of additional PCR cycles corresponding to one-third of the maximum value, rounded up. Barcodes were added to samples using primers supplied by QB3-Berkeley using PCR as described in the Corces et al. 2017 Omni-ATAC-Seq paper. Following amplification, adapters were removed from libraries using Ampure bead purification and analyzed using a Bioanalyzer or Fragment Analyzer machine to assess library quality. Final libraries were pooled together to have relatively equal proportions based on additional qPCR quantification and size-selected using a Pippin Prep to remove fragments greater than 1.5kb in size. All libraries were sequenced on one lane of each of Illumina NovaSeq SP 150PE and NovaSeq S1 150PE. Adapters used for libraries are listed in Supplementary Table 1.

### Omni-ATAC Sequencing Quality Control

Adapter trimming was performed using Trim Galore, which leverages Cutadapt and FASTQC (Andrews, 2010; Kreuger, 2017; Martin, 2011). FASTQC was also performed before and after adapter trimming to confirm removal of sequencing and Tn5 adapters. Percentage of reads remaining after deduplication was estimated based on FASTQC metrics. Library size was estimated by multiplying raw read count by percentage of reads remaining after deduplication for each lane, and then summing the two lanes. Reads were aligned to the phaw_5.0 genome as well as the *Parhyale* chrM using Bowtie2, and percentage read mapping was determined using Bowtie2 output (Langmead and Salzberg, 2012). After peak calling, merged peaks were used to evaluate correlation between replicates. Replicates were highly correlated, with a mean Pearson correlation of 0.988 and a mean Spearman correlation of 0.936 (Supp. Fig. 2 H, I).

### Omni-ATAC Peak Calling

Aligned reads were quality-filtered with a Q value of 10 in samtools and used for Genrich analysis. Output .bedgraph-like files from Genrich were reformatted using a custom Python script to be standard .bedgraph files, which were converted to bigWig files using the bedGraphToBigWig executable from the UCSC Genome Tools software bundle. Output .narrowPeak files from Genrich were converted into .bed files for ease of visualization in IGV using a custom Python script.

### Assigning spatial categories to Omni-ATAC peaks

We first assigned the nearest peak within 5kb of the first 200bp of each gene (mRNA and ncRNA) as a promoter peak using bedtools closest. We then assigned the remaining peaks into categories based on their position relative to mRNA and ncRNA annotations. Peaks that overlapped with mRNA and ncRNA annotations were assigned as exonic or intronic regulatory elements. The remaining peaks—those which had not been classified as promoters, and which did not overlap with genes—were classified as intergenic peaks. The intergenic peaks were divided into two categories: proximal and distal intergenic peaks.

Proximal peaks were those less than 10kb away from the nearest gene, while distal intergenic peaks were those greater than 10kb away from the nearest gene.

### Differential Expression and Accessibility Analyses

We performed differential expression analysis using ImpulseDE2 and DESeq2 using standard settings. To generate the Omni-ATAC-Seq read count matrix, we used bedtools multicov, using the merged Genrich peaks as our regions of interest. For our ImpulseDE2 analyses using the Omni-ATAC-Seq data, ImpulseDE2 model fits were extracted from the ImpulseDE2 output and used to visualize model expression using a custom Python script (see Supplementary Methods). To generate the RNA-Seq read count matrix for DESeq2, we generated a gene_trans_map file for the Mikado transcriptome, as would be available for the Trinity RNA-Seq analysis pipeline, and used the built-in Trinity differential expression pipeline (align_and_estimate_abundance.pl, abundance_estimates_to_matrix.pl) with Kallisto to generate a matrix of read counts. To comply with the requirement for integer counts in DESeq2 analysis, we rounded each value to the nearest whole number.

### NucleoATAC Nucleosome Predictions

Quality-filtered reads from each biological duplicate were merged and analyzed using NucleoATAC, with genomic regions set as +/- 500bp windows around Genrich peaks.

### HINT-ATAC Transcription Factor Footprinting

Quality-filtered reads from each biological duplicate were merged and analyzed using rgt-hint footprinting.

We used the JASPAR2020 database and converted the position weight matrices from JASPAR format into a simple matrix format expected by RGT-HINT using the R package “universalmotif,” and generated a “.mtf” file to store database information (see Supplementary Methods). For enrichment analyses, we used bedtools random to generate 13 million random 20bp sequences, as this was the average footprint size of genuine footprints detected by RGT-HINT in our data. This set of random sequences was used as background for our enrichment analyses. For cluster-specific enrichment analyses, we collated all unique transcription factor footprints from all developmental stages for each cluster (e.g. all footprints across S13, S14, etc. for all peaks in a given cluster) and compared enrichment levels to our randomly generated background.

### Minos Transposon Cloning

Minos transposon reporter plasmids were cloned using Gibson homology-mediated cloning approaches and the NEB Gibson Assembly or NEBuilder kits. As a base plasmid, we used the pMi(ne1) plasmid, which contains the Hsp70 minimal promoter, a DsRed protein sequence, and an SV40 3’UTR sequence, as well as two Minos inverted repeats. For plasmids containing the Hsp70 minimal promoter, the insert was integrated between the EcoRV and BglII restriction sites. For plasmids containing an endogenous promoter, the insert replaced the sequence between the EcoRV and NcoI restriction sites, thereby removing the Hsp70 minimal promoter. Cloned plasmids were Sanger sequenced to confirm a full DsRed ORF and inclusion of desired genomic sequences.

### Minos Transposase Assay

Minos transposase mRNA was generated using the ThermoFisher mMESSAGE mMACHINE T7 or T7 ULTRA kit using NotI-digested pBlueSK-MimRNA (Addgene #102535). mRNA and concentrated DNA were mixed into a final concentration of 1 µg/µL in a solution of 0.1% phenol red in nuclease-free water. 1- and 2-cell *Parhyale* embryos were injected with approximately 3-5 picoliters of injection mix using a borosilicate glass capillary needle pulled using a Sutter P-80 or P-85 instrument. Embryos were raised until hatching and examined once per day from 3dpf–10dpf using a Zeiss LSM780 confocal microscope to screen for DsRed fluorescence.

## Data Availability

Omni-ATAC-Seq raw reads will be available at SRA, Bioproject: PRJNA765106. RNA-Seq raw reads will be available at SRA. Omni-ATAC and RNA-Seq downstream analyses will be available at GEO by time of future publication. Downstream analysis files for Omni-ATAC-Seq will include BigWig files for visualizing Omni-ATAC read pileups; stage-specific and combined Genrich peaks; NucleoATAC signal, occupancy, insert, v-plots, and other miscellaneous data; HINT-ATAC transcription factor footprinting predictions and enrichment statistics; JASPAR2020 converted database and .mtf file for HINT-ATAC analyses; and differential accessibility analysis tables. Downstream analysis files for RNA-Seq will include BigWig files for visualizing RNA-Seq read pileups; Nanopore alignment sequences; individual transcriptome GFF and FASTA files; eggNOG annotations and OrthoFinder annotations of transcripts; and RNA-Seq differential expression analysis tables. In addition, the GEO database will contain the updated Mikado genome annotation and assignment of peaks based on their position relative to genes, along with classification data for peaks as concordant or discordant in expression and accessibility. Code for analysis and visualizations will be curated and deposited in a GitHub repository prior to publication.

## Contributions

D.A.S. conceived of, designed, and performed experiments and analyses. D.A.S. wrote and revised the manuscript. H.S.B. collected sequencing data for the limb development transcriptome and J.V.B. assembled it. N.H.P. provided guidance and supervised the project. J.V.B., N.H.P., and H.S.B. and provided feedback on the manuscript.

## Acknowledgments

We thank Jase Gehring and Jenna Haines for providing tips and reagents for performing and troubleshooting ATAC-Seq, and Kasia Oktaba for suggesting the technique. Jenna Haines and Shaked Afik also provided helpful programming references for performing ATAC-Seq analyses. We are grateful to Aaron Pomerantz for helping to troubleshoot and perform Nanopore sequencing, and for tips on RNA-Seq analysis. This work would have been much more challenging without fantastic sequencing support from the QB3 Biosciences Functional Genomics Lab, particularly Shana McDevitt, Karen Lundy, and Justin Choi. We are also grateful to Dan Rokhsar for helpful comments on the data analysis and manuscript. Thank you to the members of the Patel Lab, the Miller Lab, and the Garcia Lab for providing feedback and suggestions to the manuscript, particularly Craig Miller, Tyler Square, and Brandon Schlomann.

## Conflicts of Interest

The authors declare no conflicts of interest.

## Notes

### Competing Interest Statement

The authors have declared no competing interest.

### Summary of Updates

Figs. 1, 2, 4, 6, and 7 have minor changes to legends and axes. Fig. 3 has been simplified to focus more specifically on comparisons between the MAKER annotation and Mikado transcriptome, and previous data has been added to Supp. Figs. 1 and 3.1. Fig. 5 has been simplified and previous data added to Supp. Fig. 5.1. Fig. 8 has been simplified and VISTA method added to Supp. Fig. 8.1. Additional supplementary figures and tables have been generated, and supplementary figure number has been adjusted to more directly pair supplementary to primary figures. Additional minor revisions are added to prose and citations throughout the text. Throughout, we replace references to the "Blondel" annotation with references to the "MAKER" annotation. Additionally, we clarify the use of a developmental transcriptome generated by Jessen Bredeson and Heather Bruce covering limb developmental stages ("Trinity-limb") as part of the dataset, and add these two researchers as authors to the manuscript for their data contributions.

## References

1. Andrews, S. (2010). FastQC: A Quality Control Tool for High Throughput Sequence Data.

2. Brogaard, K., Xi, L., Wang, J.-P., and Widom, J. (2012). A map of nucleosome positions in yeast at base-pair resolution. Nature 486, 496–501.

3. Browne, W.E., Price, A.L., Gerberding, M., and Patel, N.H. (2005). Stages of embryonic development in the amphipod crustacean, Parhyale hawaiensis. Genesis 42, 124–149.

4. Bruce, H.S., and Patel, N.H. (2020). Knockout of crustacean leg patterning genes suggests that insect wings and body walls evolved from ancient leg segments. Nat. Ecol. Evol. 4, 1703–1712.

5. Buenrostro, J.D., Wu, B., Chang, H.Y., and Greenleaf, W.J. (2015). ATAC-seq: A Method for Assaying Chromatin Accessibility Genome-Wide. Curr Protoc Mol Biol 109, 21.29.1-9.

6. Camacho, C., Coulouris, G., Avagyan, V., Ma, N., Papadopoulos, J., Bealer, K., and Madden, T.L. (2009). BLAST+: architecture and applications. BMC Bioinformatics 10, 421.

7. Cazet, J.F., Cho, A., and Juliano, C.E. (2021). Generic injuries are sufficient to induce ectopic Wnt organizers in Hydra. ELife 10, e60562.

8. Corces, M.R., Trevino, A.E., Hamilton, E.G., Greenside, P.G., Sinnott-Armstrong, N.A., Vesuna, S., Satpathy, A.T., Rubin, A.J., Montine, K.S., Wu, B., et al. (2017). An improved ATAC-seq protocol reduces background and enables interrogation of frozen tissues. Nat. Methods 14, 959–962.

9. Costa-Silva, J., Domingues, D., and Lopes, F.M. (2017). RNA-Seq differential expression analysis: An extended review and a software tool. PLOS ONE 12, e0190152.

10. Crawford, K., Diaz Quiroz, J.F., Koenig, K.M., Ahuja, N., Albertin, C.B., and Rosenthal, J.J.C. (2020). Highly Efficient Knockout of a Squid Pigmentation Gene. Curr. Biol. 30, 3484–3490.e4.

11. Duan, Y., Zhang, W., Cheng, Y., Shi, M., and Xia, X.-Q. (2020). A systematic evaluation of bioinformatics tools for identification of long noncoding RNAs. RNA.

12. Emms, D.M., and Kelly, S. (2019). OrthoFinder: phylogenetic orthology inference for comparative genomics. Genome Biol. 20, 238.

13. Fischer, D.S., Theis, F.J., and Yosef, N. (2018). Impulse model-based differential expression analysis of time course sequencing data. Nucleic Acids Res. 46, e119–e119.

14. Fornes, O., Castro-Mondragon, J.A., Khan, A., van der Lee, R., Zhang, X., Richmond, P.A., Modi, B.P., Correard, S., Gheorghe, M., Baranašić, D., et al. (2020). JASPAR 2020: update of the open-access database of transcription factor binding profiles. Nucleic Acids Res. 48, D87–D92.

15. Gaspar, J.M. Genrich. Gatzmann, F., Falckenhayn, C., Gutekunst, J., Hanna, K., Raddatz, G., Carneiro, V.C., and Lyko, F. (2018). The methylome of the marbled crayfish links gene body methylation to stable expression of poorly accessible genes. Epigenetics Chromatin 11, 57.

16. Gehrke, A.R., Neverett, E., Luo, Y.-J., Brandt, A., Ricci, L., Hulett, R.E., Gompers, A., Ruby, J.G., Rokhsar, D.S., Reddien, P.W., et al. (2019). Acoel genome reveals the regulatory landscape of whole-body regeneration. Science 363, eaau6173.

17. Gilbert, D.G. (2019). Longest protein, longest transcript or most expression, for accurate gene reconstruction of transcriptomes? BioRxiv 829184.

18. Giresi, P.G., Kim, J., McDaniell, R.M., Iyer, V.R., and Lieb, J.D. (2007). FAIRE (Formaldehyde-Assisted Isolation of Regulatory Elements) isolates active regulatory elements from human chromatin. Genome Res. 17, 877–885.

19. Grabherr, M.G., Haas, B.J., Yassour, M., Levin, J.Z., Thompson, D.A., Amit, I., Adiconis, X., Fan, L., Raychowdhury, R., Zeng, Q., et al. (2011). Full-length transcriptome assembly from RNA-Seq data without a reference genome. Nat. Biotechnol. 29, 644–652.

20. Haas, B.J., Papanicolaou, A., Yassour, M., Grabherr, M., Blood, P.D., Bowden, J., Couger, M.B., Eccles, D., Li, B., Lieber, M., et al. (2013). De novo transcript sequence reconstruction from RNA-seq using the Trinity platform for reference generation and analysis. Nat. Protoc. 8, 1494–1512.

21. Huerta-Cepas, J., Szklarczyk, D., Heller, D., Hernández-Plaza, A., Forslund, S.K., Cook, H., Mende, D.R., Letunic, I., Rattei, T., Jensen, L.J., et al. (2019). eggNOG 5.0: a hierarchical, functionally and phylogenetically annotated orthology resource based on 5090 organisms and 2502 viruses. Nucleic Acids Res. 47, D309–D314.

22. JL, and G.-S., and Shubin, N. (2015). Deep conservation of wrist and digit enhancers in fish. Proc. ….

23. Kao, D., Lai, A.G., Stamataki, E., Rosic, S., Konstantinides, N., Jarvis, E., Donfrancesco, A., Natalia, P.-S., Semon, M., Grillo, M., et al. (2016). The genome of the crustacean Parhyale hawaiensis, a model for animal development, regeneration, immunity and lignocellulose digestion. Elife 5, 065789.

24. Kissane, S., Dhandapani, V., and Orsini, L. (2021). Protocol for assay of transposase accessible chromatin sequencing in non-model species. STAR Protoc. 2, 100341.

25. Konstantinides, N., and Averof, M. (2014). A common cellular basis for muscle regeneration in arthropods and vertebrates. Science 343, 788–791.

26. Kovaka, S., Zimin, A.V., Pertea, G.M., Razaghi, R., Salzberg, S.L., and Pertea, M. (2019). Transcriptome assembly from long-read RNA-seq alignments with StringTie2. Genome Biol. 20, 278.

27. Kreuger, F. (2017). TrimGalore.

28. Kumar, L., and E Futschik, M. (2007). Mfuzz: a software package for soft clustering of microarray data. Bioinformation 2, 5–7.

29. Lai, Y.-T., Deem, K.D., Borràs-Castells, F., Sambrani, N., Rudolf, H., Suryamohan, K., El-Sherif, E., Halfon, M.S., McKay, D.J., and Tomoyasu, Y. (2018). Enhancer identification and activity evaluation in the red flour beetle, Tribolium castaneum. Development 145.

30. Langmead, B., and Salzberg, S.L. (2012). Fast gapped-read alignment with Bowtie 2. Nat. Methods 9, 357–359.

31. Li, H. (2018). Minimap2: pairwise alignment for nucleotide sequences. Bioinformatics 34, 3094–3100.

32. Li, Y., Chen, C., Kaye, A.M., and Wasserman, W.W. (2015). The identification of cis-regulatory elements: A review from a machine learning perspective. Biosystems 138, 6–17.

33. Li, Z., Schulz, M.H., Look, T., Begemann, M., Zenke, M., and Costa, I.G. (2019). Identification of transcription factor binding sites using ATAC-seq. Genome Biol. 20, 45.

34. Liu, B., Li, J., and Cairns, M.J. (2014). Identifying miRNAs, targets and functions. Brief. Bioinform. 15, 1–19.

35. Love, M.I., Huber, W., and Anders, S. (2014). Moderated estimation of fold change and dispersion for RNA-seq data with DESeq2. Genome Biol. 15, 550.

36. Ludwig, M.Z., Patel, N.H., and Kreitman, M. (1998). Functional analysis of eve stripe 2 enhancer evolution in Drosophila: rules governing conservation and change. Development 125, 949–958.

37. Mahony, S., and Benos, P.V. (2007). STAMP: a web tool for exploring DNA-binding motif similarities. Nucleic Acids Res. 35, W253–W258.

38. Martin, M. (2011). Cutadapt removes adapter sequences from high-throughput sequencing reads. EMBnetjournal Vol 17 No 1 Gener. Seq. Data Anal. - 1014806ej171200.

39. Martin, A., Serano, J.M., Jarvis, E., Bruce, H.S., Wang, J., Ray, S., Barker, C.A., C. O., Liam, and Patel, N.H. (2015). CRISPR/Cas9 Mutagenesis Reveals Versatile Roles of Hox Genes in Crustacean Limb Specification and Evolution. Curr Biol 26, 14–26.

40. Mattioli, K., Oliveros, W., Gerhardinger, C., Andergassen, D., Maass, P.G., Rinn, J.L., and Melé, M. (2020). Cis and trans effects differentially contribute to the evolution of promoters and enhancers. Genome Biol. 21, 210.

41. Mito, T., Nakamura, T., Bando, T., Ohuchi, H., and Noji, S. (2011). The advent of RNA interference in Entomology. Entomol. Sci. 14, 1–8.

42. Ofran, Y., Mysore, V., and Rost, B. (2007). Prediction of DNA-binding residues from sequence. Bioinformatics 23, i347–i353.

43. Paris, M., Wolff, C., Patel, N.H., and Averof, M. (2021). The Crustacean Model Parhyale hawaiensis. Preprints.

44. Pavlopoulos, A., and Averof, M. (2005). Establishing genetic transformation for comparative developmental studies in the crustacean Parhyale hawaiensis. Proc Natl Acad Sci USA 102, 7888–7893.

45. Pavlopoulos, A., Kontarakis, Z., Liubicich, D.M., Serano, J.M., Akam, M., Patel, N.H., and Averof, M. (2009). Probing the evolution of appendage specialization by Hox gene misexpression in an emerging model crustacean. Proc Natl Acad Sci USA 106, 13897–13902.

46. Pérez-Zamorano, B., Rosas-Madrigal, S., Lozano, O.A.M., Castillo Méndez, M., and Valverde-Garduño, V. (2017). Identification of cis-regulatory sequences reveals potential participation of lola and Deaf1 transcription factors in Anopheles gambiae innate immune response. PloS One 12, e0186435–e0186435.

47. Picelli, S., Björklund, Å.K., Reinius, B., Sagasser, S., Winberg, G., and Sandberg, R. (2014). Tn5 transposase and tagmentation procedures for massively-scaled sequencing projects. Genome Res 24, gr.177881.114.

48. Poynton, H.C., Hasenbein, S., Benoit, J.B., Sepulveda, M.S., Poelchau, M.F., Hughes, D.S.T., Murali, S.C., Chen, S., Glastad, K.M., Goodisman, M.A.D., et al. (2018). The Toxicogenome of Hyalella azteca: A Model for Sediment Ecotoxicology and Evolutionary Toxicology. Environ. Sci. Technol. 52, 6009–6022.

49. Quinlan, A.R., and Hall, I.M. (2010). BEDTools: a flexible suite of utilities for comparing genomic features. Bioinformatics 26, 841–842.

50. Radman-Livaja, M., and Rando, O.J. (2010). Nucleosome positioning: how is it established, and why does it matter? Dev. Biol. 339, 258–266.

51. Ramos, A.P., Gustafsson, O., Labert, N., Salecker, I., Nilsson, D.-E., and Averof, M. (2019). Analysis of the genetically tractable crustacean Parhyale hawaiensis reveals the organisation of a sensory system for low-resolution vision. BMC Biol. 17, 67.

52. Rasys, A.M., Park, S., Ball, R.E., Alcala, A.J., Lauderdale, J.D., and Menke, D.B. (2019). CRISPR-Cas9 Gene Editing in Lizards through Microinjection of Unfertilized Oocytes. Cell Rep. 28, 2288–2292.e3.

53. Ratnere, I., and Dubchak, I. (2009). Obtaining Comparative Genomic Data with the VISTA Family of Computational Tools. Curr. Protoc. Bioinforma. 26, 10.6.1-10.6.17.

54. Rehm, E.J., Hannibal, R.L., Chaw, R.C., Vargas-Vila, M.A., and Patel, N.H. (2009). Fixation and Dissection of Parhyale hawaiensis Embryos. Cold Spring Harb. Protoc. 2009, pdb.prot5127.

55. Robinson, M.D., McCarthy, D.J., and Smyth, G.K. (2010). edgeR: a Bioconductor package for differential expression analysis of digital gene expression data. Bioinforma. Oxf. Engl. 26, 139–140.

56. Rouhana, L., Weiss, J.A., Forsthoefel, D.J., Lee, H., King, R.S., Inoue, T., Shibata, N., Agata, K., and Newmark, P.A. (2013). RNA interference by feeding in vitro-synthesized double-stranded RNA to planarians: methodology and dynamics. Dev. Dyn. Off. Publ. Am. Assoc. Anat. 242, 718–730.

57. Schep, A.N., Buenrostro, J.D., Denny, S.K., Schwartz, K., Sherlock, G., and Greenleaf, W.J. (2015). Structured nucleosome fingerprints enable high-resolution mapping of chromatin architecture within regulatory regions. Genome Res.

58. Schmitt-Engel, C., Schultheis, D., Schwirz, J., Ströhlein, N., Troelenberg, N., Majumdar, U., Dao, V.A., Grossmann, D., Richter, T., Tech, M., et al. (2015). The iBeetle large-scale RNAi screen reveals gene functions for insect development and physiology. Nat. Commun. 6, 7822.

59. Serano, J.M., Martin, A., Liubicich, D.M., Jarvis, E., Bruce, H.S., La, K., Browne, W.E., Grimwood, J., and Patel, N.H. (2016). Comprehensive analysis of Hox gene expression in the amphipod crustacean Parhyale hawaiensis. Dev Biol 409, 297–309.

60. Sharma, P.P., Schwager, E.E., Giribet, G., Jockusch, E.L., and Extavour, C.G. (2013). Distal-less and dachshund pattern both plesiomorphic and apomorphic structures in chelicerates: RNA interference in the harvestman Phalangium opilio (Opiliones). Evol. Dev. 15, 228–242.

61. Signor, S.A., and Nuzhdin, S.V. (2018). The Evolution of Gene Expression in cis and trans. Trends Genet. 34, 532–544.

62. Srivastava, M., Mazza-Curll, K.L., van Wolfswinkel, J.C., and Reddien, P.W. (2014). Whole-Body Acoel Regeneration Is Controlled by Wnt and Bmp-Admp Signaling. Curr. Biol. 24, 1107–1113.

63. Stamataki, E., and Pavlopoulos, A. (2016). Non-insect crustacean models in developmental genetics including an encomium to Parhyale hawaiensis. Curr Opin Genet Dev 39, 149–156.

64. Sun, D.A., and Patel, N.H. (2019). The amphipod crustacean Parhyale hawaiensis: An emerging comparative model of arthropod development, evolution, and regeneration. WIREs Dev. Biol. 8, e355.

65. The UniProt Consortium (2019). UniProt: a worldwide hub of protein knowledge. Nucleic Acids Res. 47, D506–D515.

66. Venturini, L., Caim, S., Kaithakottil, G.G., Mapleson, D.L., and Swarbreck, D. (2018). Leveraging multiple transcriptome assembly methods for improved gene structure annotation. GigaScience 7.

67. Wang, L., and Brown, S.J. (2006). BindN: a web-based tool for efficient prediction of DNA and RNA binding sites in amino acid sequences. Nucleic Acids Res. 34, W243–W248.

68. Wang, Q., Gu, L., Adey, A., Radlwimmer, B., Wang, W., Hovestadt, V., Bähr, M., Wolf, S., Shendure, J., Eils, R., et al. (2013). Tagmentation-based whole-genome bisulfite sequencing. Nat Protoc 8, 2022–2032.

69. Wasserman, W.W., and Sandelin, A. (2004). Applied bioinformatics for the identification of regulatory elements. Nat Rev Genet 5, 276–287.

70. Wetterstramd, K. DNA Sequencing Costs: Data from the NHGRI Genome Sequencing Program (GSP).

71. Wittkopp, P.J., Haerum, B.K., and Clark, A.G. (2004). Evolutionary changes in cis and trans gene regulation. Nature 430, 85–88.

72. Yan, F., Powell, D.R., Curtis, D.J., and Wong, N.C. (2020). From reads to insight: a hitchhiker’s guide to ATAC-seq data analysis. Genome Biol. 21, 22.

