## Supplementary figures and images for "Identification and classification of cis-regulatory elements in the amphipod crustacean *Parhyale hawaiensis*"

### Supplementary Video 1

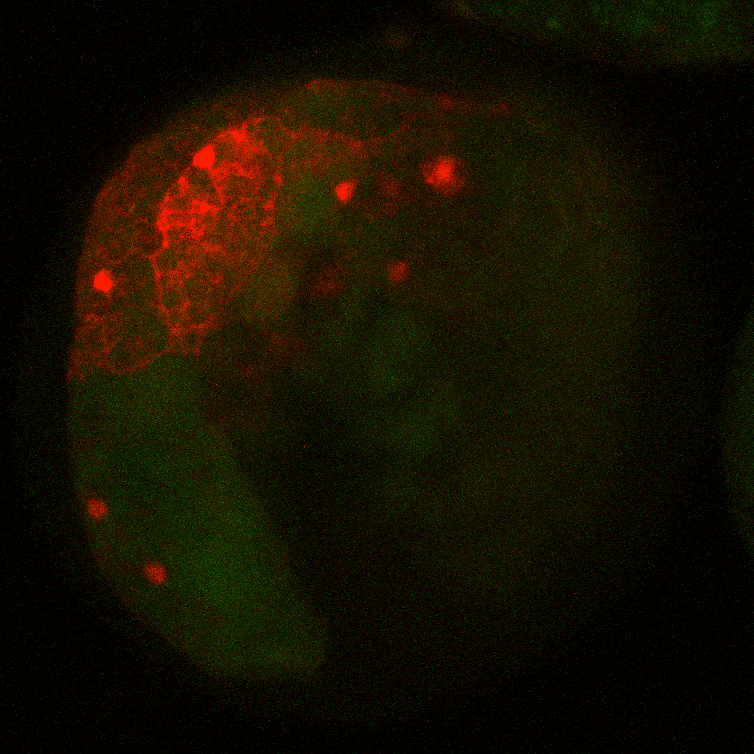
