## Supplementary Figures and Methods for "Identification and classification of cis-regulatory elements in the amphipod crustacean *Parhyale hawaiensis*"

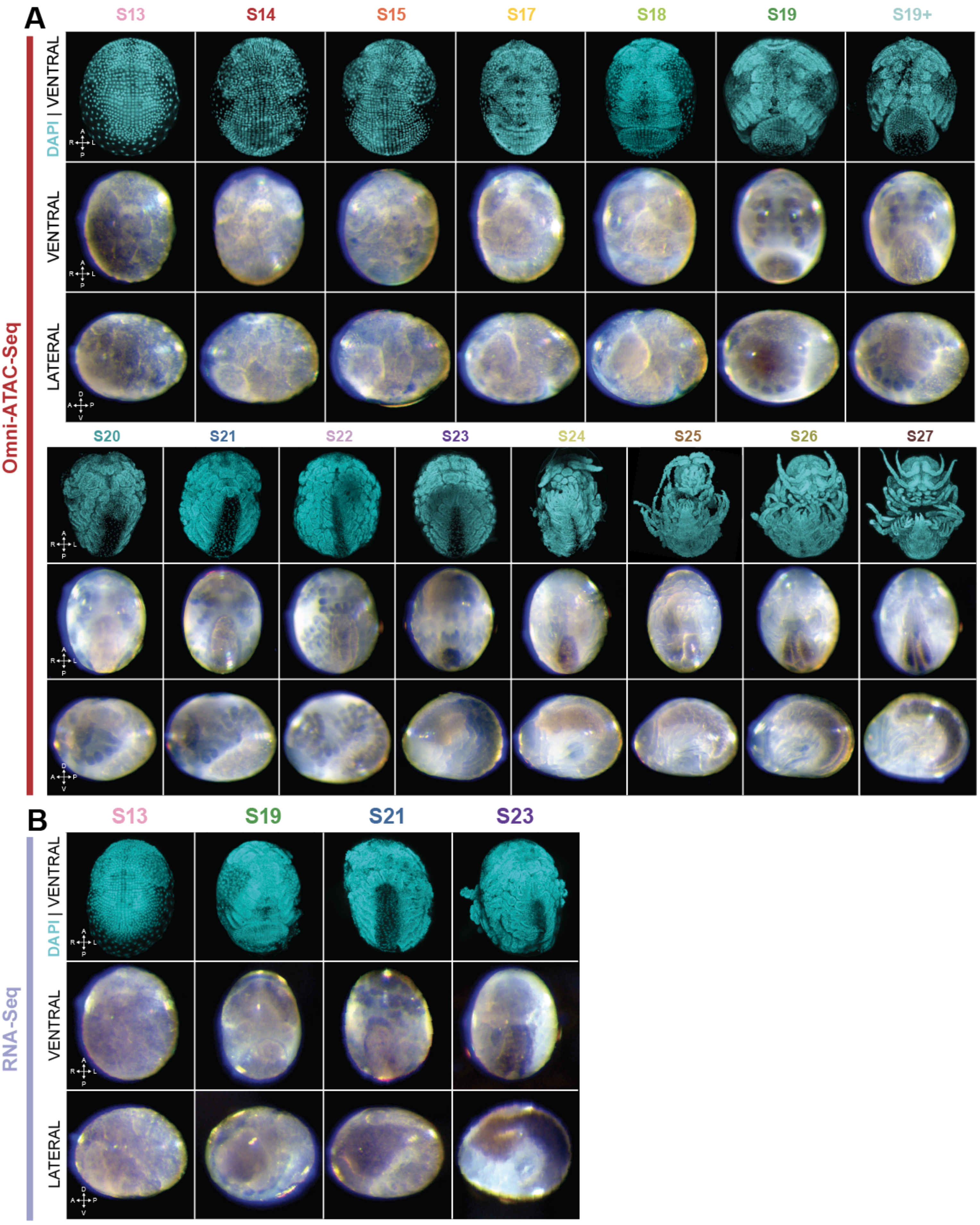

A
S13
S14
S15
S17
S18
S19
S19+
DAPI | VENTRAL
VENTRAL
LATERAL
Omni-ATAC-Seq
S20
S21
S22
S23
S24
S25
S26
S27
B
S13
S19
S21
S23
DAPI | VENTRAL
VENTRAL
LATERAL
RNA-Seq

#### Supp. Fig. 1: Embryo images for time course Omni-ATAC-Seq and RNA-Seq

A) Representative embryo images from Omni-ATAC-Seq libraries. Top row shows ventral view of DAPI stained embryo. Bottom rows show ventral and lateral brightfield images of embryos shortly before tagmentation.

B) Representative embryo images from RNA-Seq libraries. Top row shows ventral view of DAPI stained embryo. Bottom rows show ventral and lateral brightfield images of embryos shortly before RNA extraction.

(A: anterior, P: posterior, L: left, R: right; D: dorsal, V: ventral).

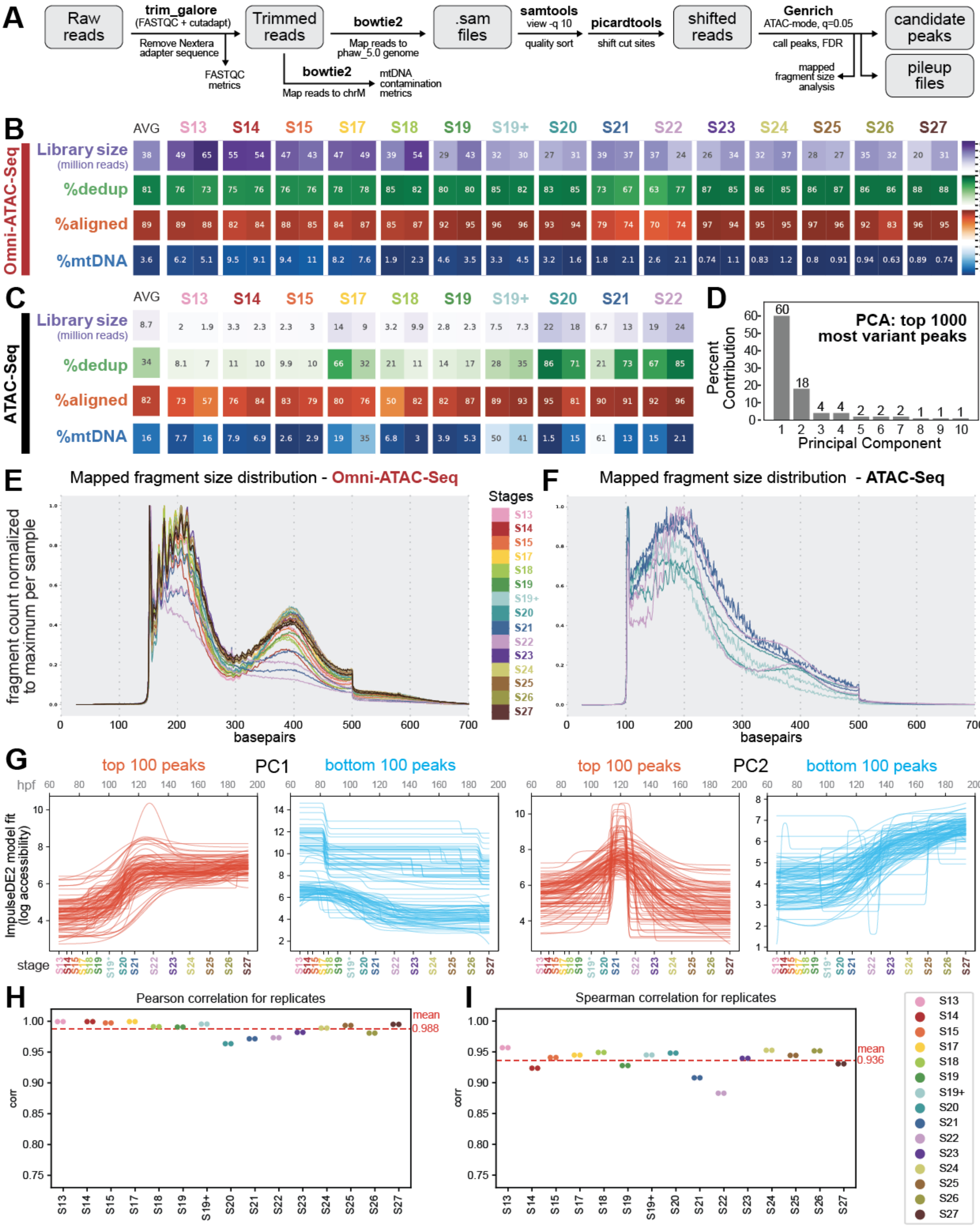
A
Raw reads
trim_galore (FASTQC + cutadapt)
Remove Nextera adapter sequence
FASTQC metrics
Trimmed reads
bowtie2
Map reads to phaw_5.0 genome
bowtie2
Map reads to chrM
mtDNA contamination metrics
.sam files
samtools view -q 10
quality sort
picardtools
shift cut sites
shifted reads
Genrich ATAC-mode, q=0.05
call peaks, FDR
mapped fragment size analysis
candidate peaks
pileup files
B
Omni-ATAC-Seq
AVG S13 S14 S15 S17 S18 S19 S19+ S20 S21 S22 S23 S24 S25 S26 S27
Library size (million reads)
38 49 65 55 54 47 43 47 49 39 54 29 43 32 30 27 31 39 37 37 24 26 34 32 37 28 27 35 32 20 31
%dedup
81 76 73 75 76 76 76 78 78 85 82 80 80 85 82 83 85 73 67 63 77 87 85 86 85 86 87 86 86 88 88
%aligned
89 89 88 82 84 88 85 84 87 85 87 92 95 96 96 93 94 79 74 70 74 97 94 95 95 97 96 92 83 96 95
%mtDNA
3.6 6.2 5.1 9.5 9.1 9.4 11 8.2 7.6 1.9 2.3 4.6 3.5 3.3 4.5 3.2 1.6 1.8 2.1 2.6 2.1 0.74 1.1 0.83 1.2 0.8 0.91 0.94 0.63 0.89 0.74
C
ATAC-Seq
AVG S13 S14 S15 S17 S18 S19 S19+ S20 S21 S22
Library size (million reads)
8.7 2 1.9 3.3 2.3 2.3 3 14 9 3.2 9.9 2.8 2.3 7.5 7.3 22 18 6.7 13 19 24
%dedup
34 8.1 7 11 10 9.9 10 66 32 21 11 14 17 28 35 86 71 21 73 67 85
%aligned
82 73 57 76 84 83 79 80 76 50 82 82 87 89 93 95 81 90 91 92 96
%mtDNA
16 7.7 16 7.9 6.9 2.6 2.9 19 35 6.8 3 3.9 5.3 50 41 1.5 15 61 13 15 2.1
D
PCA: top 1000 most variant peaks
Percent Contribution
60 18 4 4 2 2 2 1 1 1
Principal Component
E
Mapped fragment size distribution - Omni-ATAC-Seq
fragment count normalized to maximum per sample
basepairs
Stages
S13 S14 S15 S17 S18 S19 S19+ S20 S21 S22 S23 S24 S25 S26 S27
F
Mapped fragment size distribution - ATAC-Seq
basepairs
G
top 100 peaks PC1 bottom 100 peaks
top 100 peaks PC2 bottom 100 peaks
hpf
ImpulseDE2 model fit (log accessibility)
stage
H
Pearson correlation for replicates
mean 0.988
corr
I
Spearman correlation for replicates
mean 0.936
corr
S13 S14 S15 S17 S18 S19 S19+ S20 S21 S22 S23 S24 S25 S26 S27

#### Supp. Fig. 2: Omni-ATAC-Seq analysis pipeline and quality control metrics

A) Analysis pipeline for generating Omni-ATAC peaks and QC metrics from raw reads. Reads were quality and adapter-trimmed using trim_galore, and trimmed reads were mapped to either the entire genome or to only the mitochondrial chromosome (ChrM) to generate a conservative estimate of mtDNA contamination. Mapped reads were further filtered using samtools (Q = 10) and read ends were shifted using Picardtools to reflect the mechanism of Tn5 insertion. Shifted reads were passed to Genrich in ATAC mode using a q-value cutoff of 0.05. Files produced by Genrich included stage-specific peaks and bedgraph-like read pileup formats.

B) Quality control metrics for Omni-ATAC-Seq libraries. Library size, percentage of reads remaining after deduplication (%dedup), percentage of aligned reads (%aligned), and percentage of reads mapping to the mitochondrial genome (%mtDNA) are displayed for individual Omni-ATAC libraries.

C) Metrics from B visualized for conventional ATAC-Seq libraries. Overall, Omni-ATAC seq shows similar or improved performance on all metrics.

D) Percentage contribution to variance of each of the top 10 principal components from PCA in Fig. 2C.

E) Mapped fragment size distribution of Omni-ATAC-Seq reads. Two clear peaks are seen in the fragment size distribution, reflecting proper sub-nucleosomal insertions.

F) Mapped fragment size distribution of conventional ATAC-Seq reads for 4 developmental stages: S19, S19+, S20, S21. Fragment size peaks are less distinct as compared to Omni-ATAC-Seq libraries.

G) Plots of ImpulseDE2 model fits to the top 100 and bottom 100 peaks loading for PC1 and PC2 in figure 1C. PC1 appears to be associated with developmental time, while PC2 appears to be associated with peaks that show an increase in accessibility in the middle of development.

H) Pearson correlation of mapped read counts in merged peaks between replicates for each developmental stage. Replicate libraries show a high average correlation value (0.988).

I) Spearman correlation of mapped read counts in merged peaks between replicates for each developmental stage. Replicate libraries show a high average correlation value (0.936). A lower correlation coefficient compared to Pearson may suggest that strong outliers contribute to the high Pearson correlation measure.

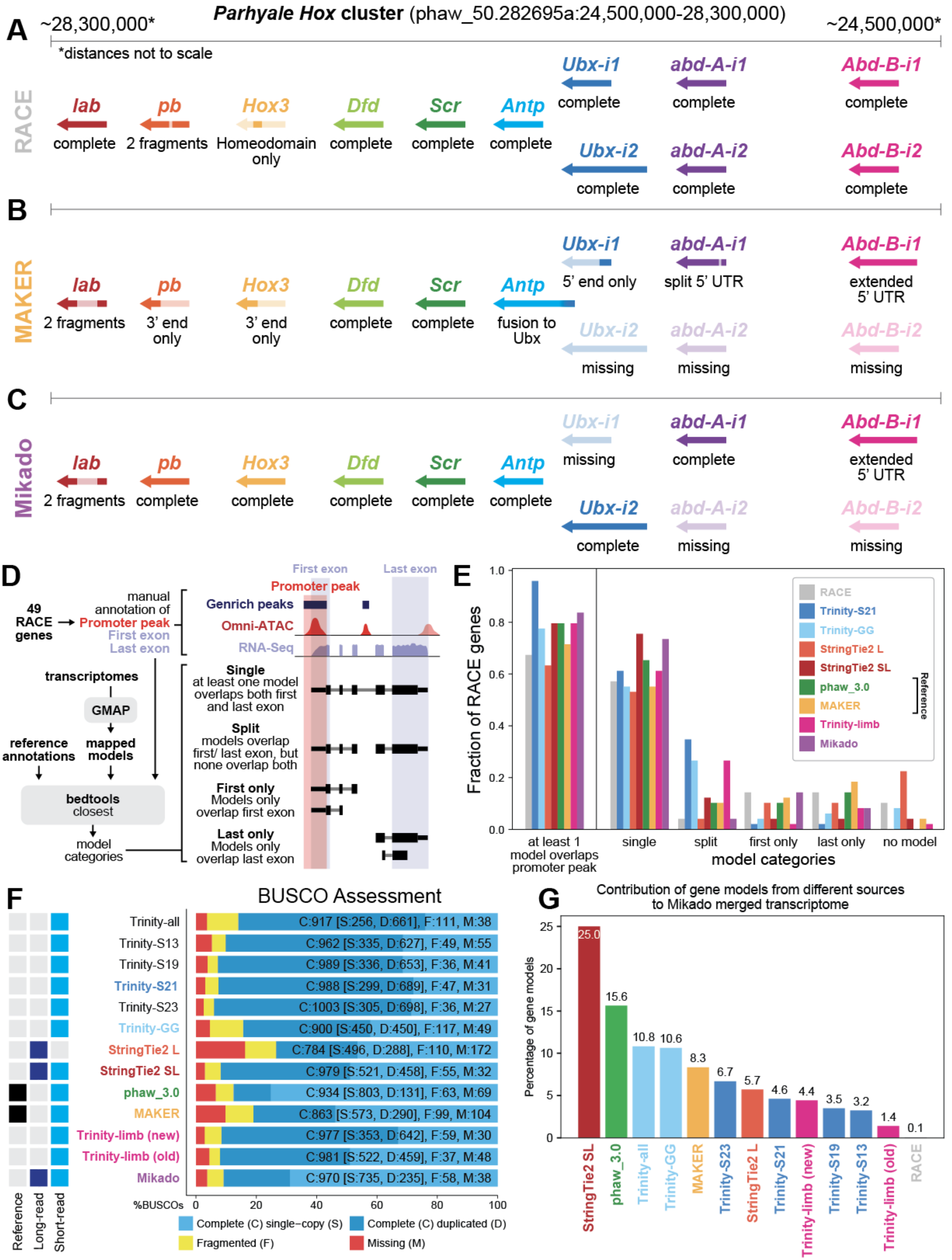
A
Parhyale Hox cluster (phaw_50.282695a:24,500,000-28,300,000)
~28,300,000*
~24,500,000*
*distances not to scale
RACE
lab complete
pb 2 fragments
Hox3 Homeodomain only
Dfd complete
Scr complete
Antp complete
Ubx-i1 complete
abd-A-i1 complete
Abd-B-i1 complete
Ubx-i2 complete
abd-A-i2 complete
Abd-B-i2 complete
B
MAKER
lab 2 fragments
pb 3' end only
Hox3 3' end only
Dfd complete
Scr complete
Antp fusion to Ubx
Ubx-i1 5' end only
abd-A-i1 split 5' UTR
Abd-B-i1 extended 5' UTR
Ubx-i2 missing
abd-A-i2 missing
Abd-B-i2 missing
C
Mikado
lab 2 fragments
pb complete
Hox3 complete
Dfd complete
Scr complete
Antp complete
Ubx-i1 missing
abd-A-i1 complete
Abd-B-i1 extended 5' UTR
Ubx-i2 complete
abd-A-i2 missing
Abd-B-i2 missing
D
49 RACE genes
manual annotation of
Promoter peak
First exon
Last exon
Genrich peaks
Omni-ATAC
RNA-Seq
transcriptomes
GMAP
reference annotations
mapped models
bedtools closest
model categories
Single
at least one model overlaps both first and last exon
Split
models overlap first/ last exon, but none overlap both
First only
Models only overlap first exon
Last only
Models only overlap last exon
E
Fraction of RACE genes
model categories
at least 1 model overlaps promoter peak
single
split
first only
last only
no model
RACE
Trinity-S21
Trinity-GG
StringTie2 L
StringTie2 SL
phaw_3.0
MAKER
Trinity-limb
Mikado
Reference
F
BUSCO Assessment
Trinity-all C:917 [S:256, D:661], F:111, M:38
Trinity-S13 C:962 [S:335, D:627], F:49, M:55
Trinity-S19 C:989 [S:336, D:653], F:36, M:41
Trinity-S21 C:988 [S:299, D:689], F:47, M:31
Trinity-S23 C:1003 [S:305, D:698], F:36, M:27
Trinity-GG C:900 [S:450, D:450], F:117, M:49
StringTie2 L C:784 [S:496, D:288], F:110, M:172
StringTie2 SL C:979 [S:521, D:458], F:55, M:32
phaw_3.0 C:934 [S:803, D:131], F:63, M:69
MAKER C:863 [S:573, D:290], F:99, M:104
Trinity-limb (new) C:977 [S:353, D:642], F:59, M:30
Trinity-limb (old) C:981 [S:522, D:459], F:37, M:48
Mikado C:970 [S:735, D:235], F:58, M:38
%BUSCOs
Reference
Long-read
Short-read
Complete (C) single−copy (S)
Complete (C) duplicated (D)
Fragmented (F)
Missing (M)
G
Contribution of gene models from different sources to Mikado merged transcriptome
Percentage of gene models
StringTie2 SL 25.0
phaw_3.0 15.6
Trinity-all 10.8
Trinity-GG 10.6
MAKER 8.3
Trinity-S23 6.7
StringTie2 L 5.7
Trinity-S21 4.6
Trinity-limb (new) 4.4
Trinity-S19 3.5
Trinity-S13 3.2
Trinity-limb (old) 1.4
RACE 0.1

#### Supp. Fig. 3.1: Additional transcriptome and gene model evaluation metrics

A) Summarized completeness of RACE *Hox* gene models. Hox gene models for most genes are complete, including two isoforms each for *Ubx*, *abd-A*, and *Abd-B*. For *Hox3*, only a homeodomain sequence was isolated.

B) Summarized completeness of MAKER (phaw_5.0) *Hox* gene models. Many gene models show deviations from RACE sequences, including fragmentation, artefactual fusion of gene models, and extension of 5' UTRs.

C) Summarized completeness of *Hox* gene models from Mikado transcriptome. The Mikado transcriptome appears to have more complete *Hox* gene models than the MAKER genome annotation. In addition, a complete *Hox3* sequence was generated.

D) Model evaluation methods for gene model fragmentation. 49 RACE genes were selected based on the ability to unambiguously annotate a first exon, last exon, and promoter peak based on examination of all gene models, Omni-ATAC-Seq peaks, and Omni-ATAC-Seq and RNA-Seq read pileups. For each transcriptome or genome annotation, for each RACE gene, all models were compared to manual annotation windows using bedtools. If a single model from all models in a given transcriptome overlapped with the first and last exon of the manual annotation, that RACE gene was classified as a "single" model for that transcriptome. If no single model overlapped both first and last exons, but models existed that overlapped either, the RACE gene was classified as "split" for that transcriptome. If gene models only overlapped with manually annotated first or last exons, then the RACE gene was classified as "first" or "last". Finally, if no gene models overlapped, the RACE gene was classified as "no model".

E) Summary of gene model evaluations from D for select transcriptomes. Overall, the Trinity-S21 transcriptome had the highest number of models that overlapped the Omni-ATAC promoter peak, and the Mikado transcriptome had the second highest. The StringTie2 SL transcriptome had the highest number of single gene models, and the Mikado transcriptome had the second highest. For both metrics, the Mikado transcriptome outperformed the MAKER genome annotation.

F) BUSCO evaluation for additional transcriptomes included in the dataset.

G) Fraction of gene models attributed to each transcriptome in the final Mikado transcriptome. Overall, the StringTie2 SL transcriptome produced the greatest number of transcripts evaluated as "best" by the Mikado pipeline.

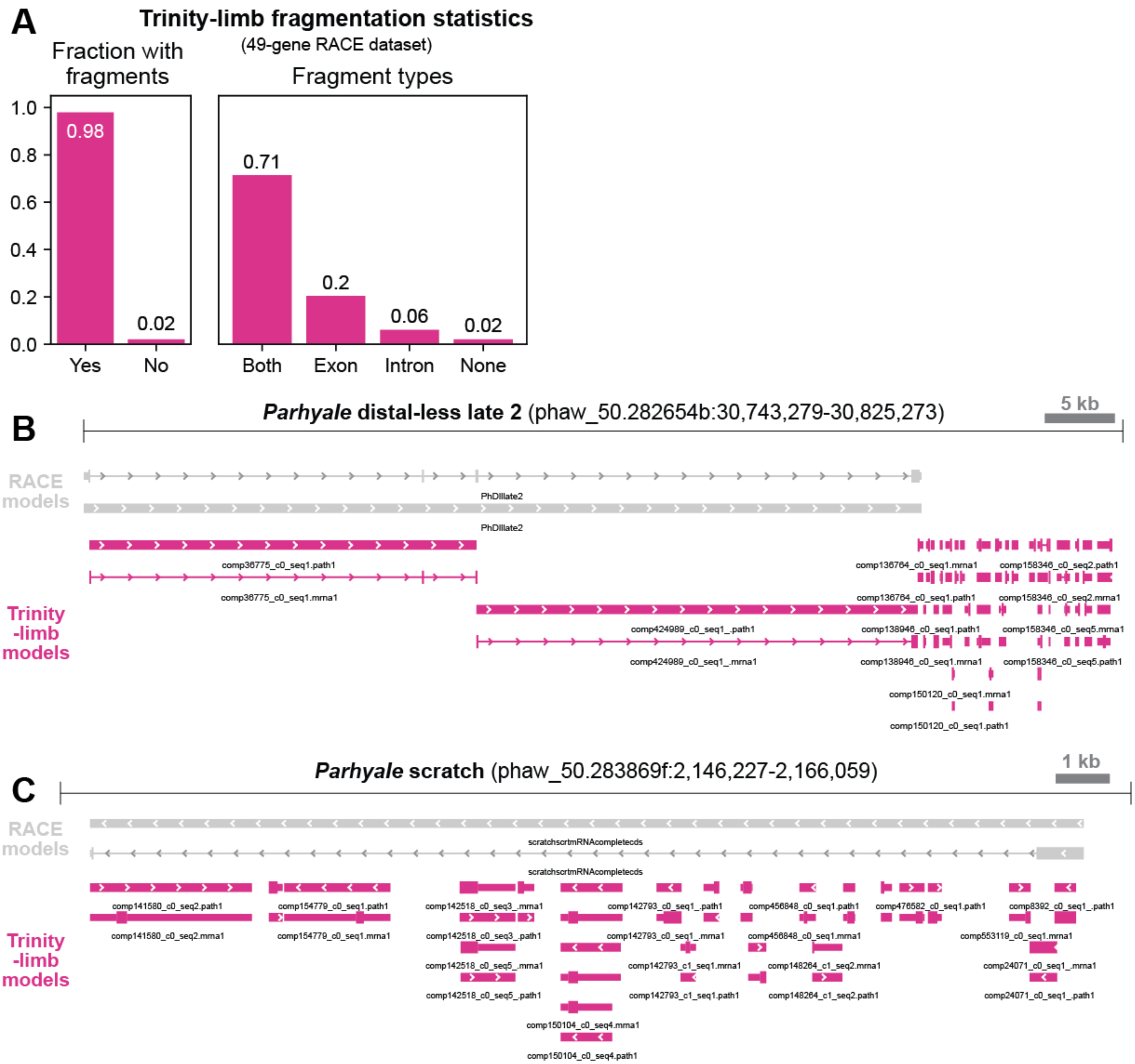
A
Trinity-limb fragmentation statistics
(49-gene RACE dataset)
Fraction with fragments
1.0
0.8
0.6
0.4
0.2
0.0
0.98
0.02
Yes
No
Fragment types
0.71
0.2
0.06
0.02
Both
Exon
Intron
None
B
Parhyale distal-less late 2 (phaw_50.282654b:30,743,279-30,825,273)
5 kb
RACE models
PhDllate2
PhDllate2
Trinity -limb models
comp36775_c0_seq1.path1
comp36775_c0_seq1.mrna1
comp424989_c0_seq1_.path1
comp424989_c0_seq1_.mrna1
comp136764_c0_seq1.mrna1
comp136764_c0_seq1.path1
comp138946_c0_seq1.path1
comp138946_c0_seq1.mrna1
comp158346_c0_seq2.path1
comp158346_c0_seq2.mrna1
comp158346_c0_seq5.mrna1
comp158346_c0_seq5.path1
comp150120_c0_seq1.mrna1
comp150120_c0_seq1.path1
C
Parhyale scratch (phaw_50.283869f:2,146,227-2,166,059)
1 kb
RACE models
scratchscrtmRNAcompletecds
scratchscrtmRNAcompletecds
Trinity -limb models
comp141580_c0_seq2.path1
comp141580_c0_seq2.mrna1
comp154779_c0_seq1.path1
comp154779_c0_seq1.mrna1
comp142518_c0_seq3_.mrna1
comp142518_c0_seq3_.path1
comp142518_c0_seq5_.mrna1
comp142518_c0_seq5_.path1
comp150104_c0_seq4.mrna1
comp150104_c0_seq4.path1
comp142793_c0_seq1_.path1
comp142793_c0_seq1_.mrna1
comp142793_c1_seq1.mrna1
comp142793_c1_seq1.path1
comp456848_c0_seq1.path1
comp456848_c0_seq1.mrna1
comp148264_c1_seq2.mrna1
comp148264_c1_seq2.path1
comp476582_c0_seq1.path1
comp8392_c0_seq1_.path1
comp553119_c0_seq1.mrna1
comp24071_c0_seq1_.mrna1
comp24071_c0_seq1_.path1

#### Supp. Fig. 3.2: Additional transcriptome and gene model evaluation metrics

A) Fraction of Trinity-limb (old) gene models with spurious transcript fragments when compared with RACE data. Fragments were observed overlapping with introns, exons, or both; some transcript fragments appeared to contain erroneous splice junctions joining exons and introns, while others were strictly intronic.

B) Example of transcript fragments compared to RACE data at the *Parhyale distal-less late 2* locus. This gene model had two transcript fragments overlapping with known exons, but no single gene model spanning the entire locus. Moreover, a large number of small 3'UTR-mapping fragments were observed, a signal that was frequently observed across genes in the RACE dataset.

C) Example of transcript fragments compared to RACE data at the *Parhyale scratch* locus. Transcript fragments that overlap with both intronic and exonic sequences are observed, as well as strictly intronic fragments. This gene model was a particularly extreme example of transcript fragmentation observed among the genes in the RACE dataset.

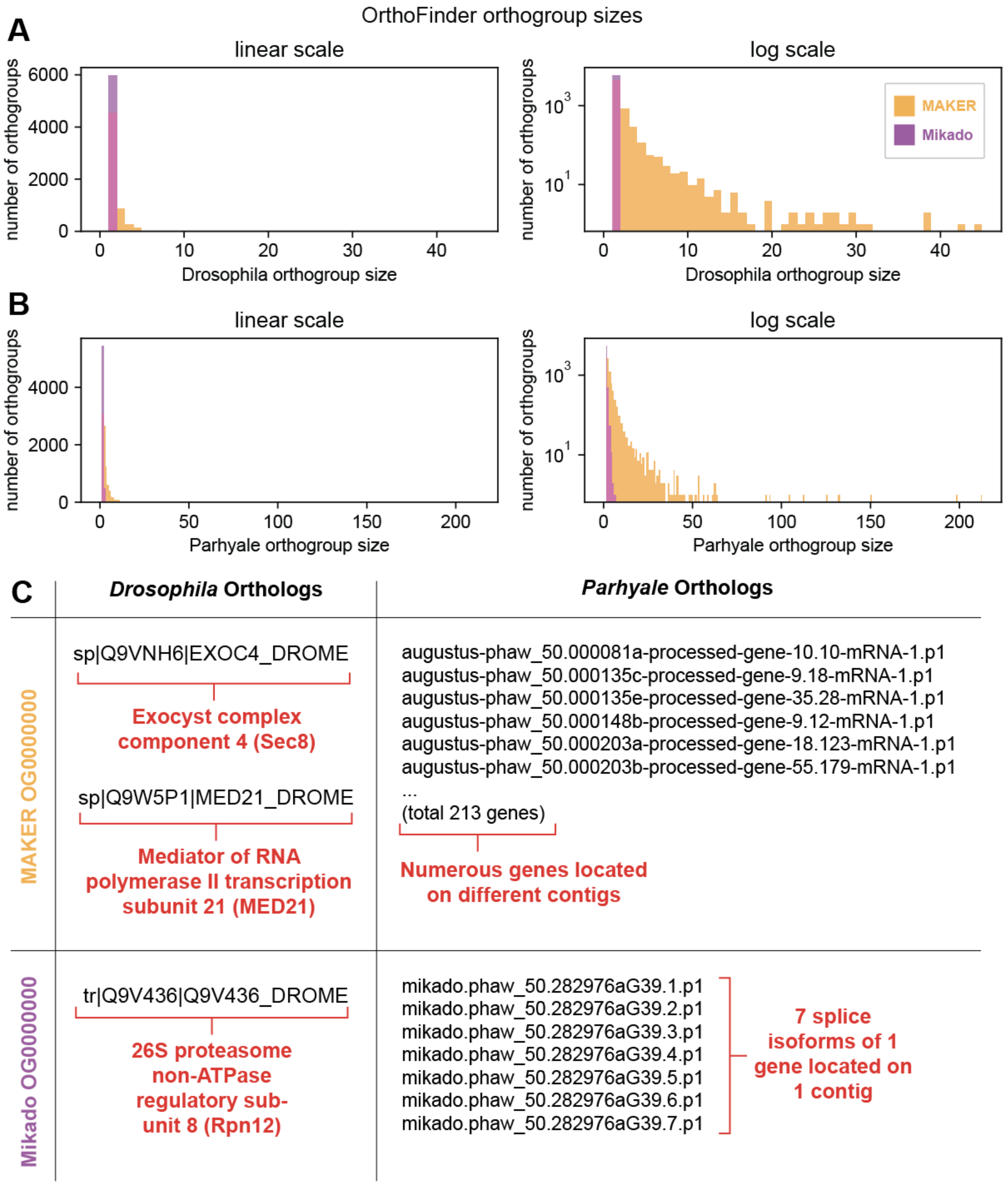

OrthoFinder orthogroup sizes
A
linear scale
log scale
number of orthogroups
Drosophila orthogroup size
MAKER
Mikado
B
linear scale
log scale
number of orthogroups
Parhyale orthogroup size
C
*Drosophila* Orthologs
*Parhyale* Orthologs
MAKER OG0000000
sp|Q9VNH6|EXOC4_DROME
Exocyst complex component 4 (Sec8)
sp|Q9W5P1|MED21_DROME
Mediator of RNA polymerase II transcription subunit 21 (MED21)
augustus-phaw_50.000081a-processed-gene-10.10-mRNA-1.p1
augustus-phaw_50.000135c-processed-gene-9.18-mRNA-1.p1
augustus-phaw_50.000135e-processed-gene-35.28-mRNA-1.p1
augustus-phaw_50.000148b-processed-gene-9.12-mRNA-1.p1
augustus-phaw_50.000203a-processed-gene-18.123-mRNA-1.p1
augustus-phaw_50.000203b-processed-gene-55.179-mRNA-1.p1
...
(total 213 genes)
Numerous genes located on different contigs
Mikado OG0000000
tr|Q9V436|Q9V436_DROME
26S proteasome non-ATPase regulatory sub-unit 8 (Rpn12)
mikado.phaw_50.282976aG39.1.p1
mikado.phaw_50.282976aG39.2.p1
mikado.phaw_50.282976aG39.3.p1
mikado.phaw_50.282976aG39.4.p1
mikado.phaw_50.282976aG39.5.p1
mikado.phaw_50.282976aG39.6.p1
mikado.phaw_50.282976aG39.7.p1
7 splice isoforms of 1 gene located on 1 contig

#### Supp. Fig. 3.3: Additional transcriptome and gene model evaluation metrics

A) Orthogroup sizes grouped by number of *Drosophila* orthologs in each orthogroup for the MAKER genome annotation and the Mikado transcriptome. The MAKER genome annotation, when annotated using OrthoFinder, produces many gene models that are grouped into orthogroups with large numbers of putative orthologs when compared to the results of using OrthoFinder on the Mikado transcriptome.

B) Orthogroup sizes grouped by number of *Parhyale* orthologs in each orthogroup for the MAKER genome annotation and the Mikado transcriptome. As with the *Drosophila* measure, the number of orthogroups with large numbers of orthologs is greater for the MAKER genome annotation.

C) Example of peptides spuriously classified as orthologs when comparing the MAKER genome annotation to the Mikado transcriptome. OG0000000 is the single largest orthogroup identified by OrthoFinder when comparing the *Drosophila* UNIPROT database to either the MAKER genome annotation or the Mikado transcriptome. In the MAKER annotation, the largest orthogroup contains two unrelated *Drosophila* proteins (Sec8 and MED21), which are grouped together along with 213 different *Parhyale* gene models scattered across the genome. Examination of other orthogroups revealed a similar result, in which unrelated *Drosophila* proteins were grouped along with numerous *Parhyale* gene models, impeding precise identification of orthologs between the species, and suggesting problems in the quality of the MAKER gene models. In the Mikado annotation, the largest orthogroup contains a single *Drosophila* protein (Rpn12) grouped along with 7 splice isoforms of a single *Parhyale* gene model located on a single contig. Examination of other orthogroups derived from the Mikado transcriptome revealed fewer obviously incorrect groupings than observed for the MAKER annotation.

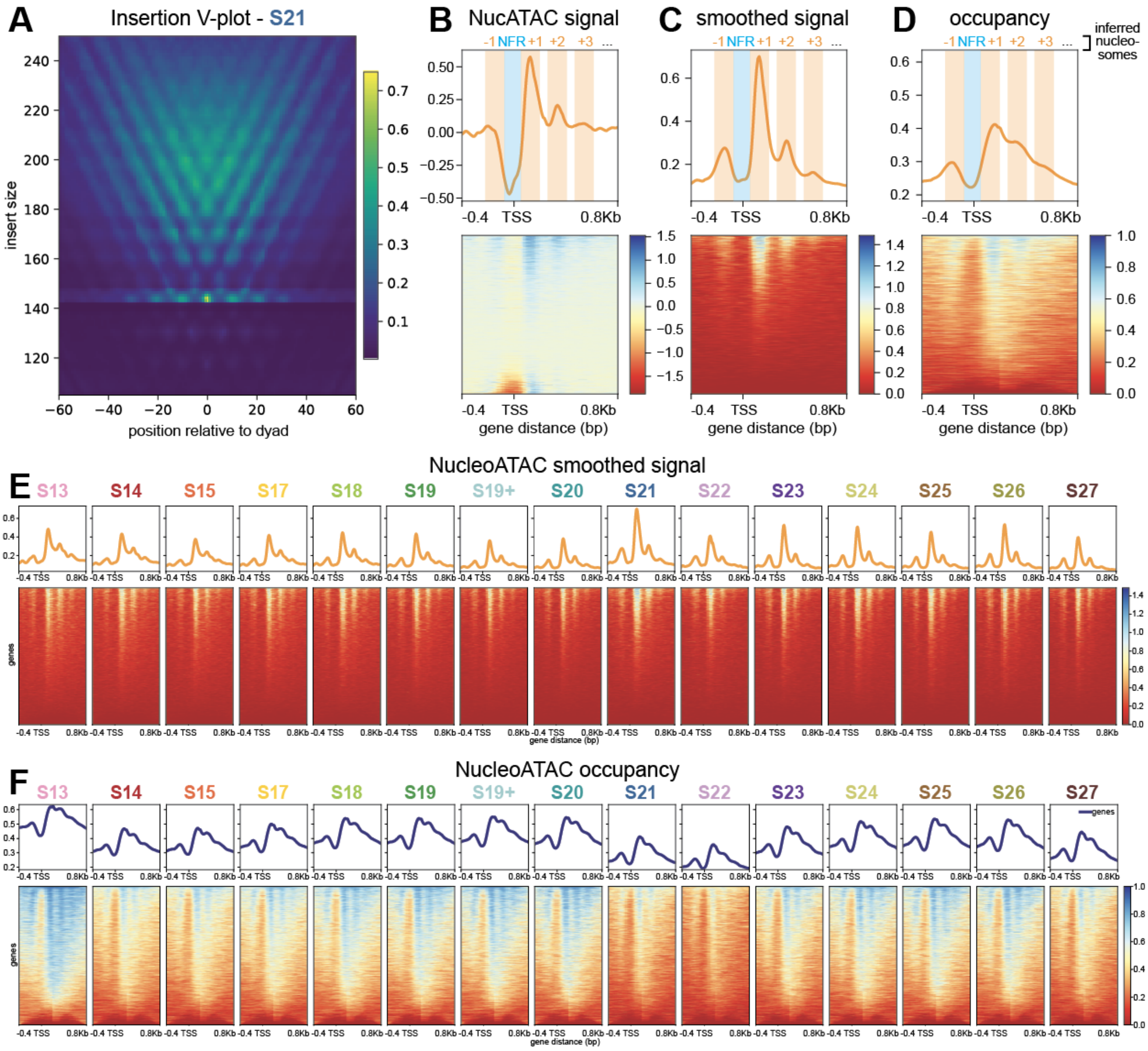
A
Insertion V-plot - S21
insert size
position relative to dyad
B
NucATAC signal
-1 NFR +1 +2 +3 ...
gene distance (bp)
C
smoothed signal
D
occupancy
inferred nucleo-somes
-0.4 TSS 0.8Kb
E
NucleoATAC smoothed signal
S13 S14 S15 S17 S18 S19 S19+ S20 S21 S22 S23 S24 S25 S26 S27
genes
F
NucleoATAC occupancy
S13 S14 S15 S17 S18 S19 S19+ S20 S21 S22 S23 S24 S25 S26 S27
genes

#### Supp. Fig. 4: NucleoATAC quality control and signal over time

A) Insertion V-plot generated from NucleoATAC using reads from S21 libraries. A clear V-shaped pattern is observed, as expected from read data fragmented along a nucleosomal distribution.

B) NucleoATAC raw signal at mRNA starts genome-wide. A strong +1 and -1 nucleosome position signal is observed. A strong negative signal is observed at the expected nucleosome-free region (NFR). Overlaid windows in B-D were drawn manually based on observed signal.

C) NucleoATAC smoothed signal at mRNA starts genome-wide. A strong +1 and -1 nucleosome position signal is observed.

D) NucleoATAC occupancy at mRNA starts genome-wide. A strong +1 and -1 nucleosome position signal is observed.

E) NucleoATAC smoothed signal at mRNA starts genome-wide for all developmental stages.

F) NucleoATAC occupancy signal at mRNA starts genome-wide for all developmental stages. The occupancy value calculated by NucleoATAC varies based on library size; in this plot, values are not normalized based on library size.

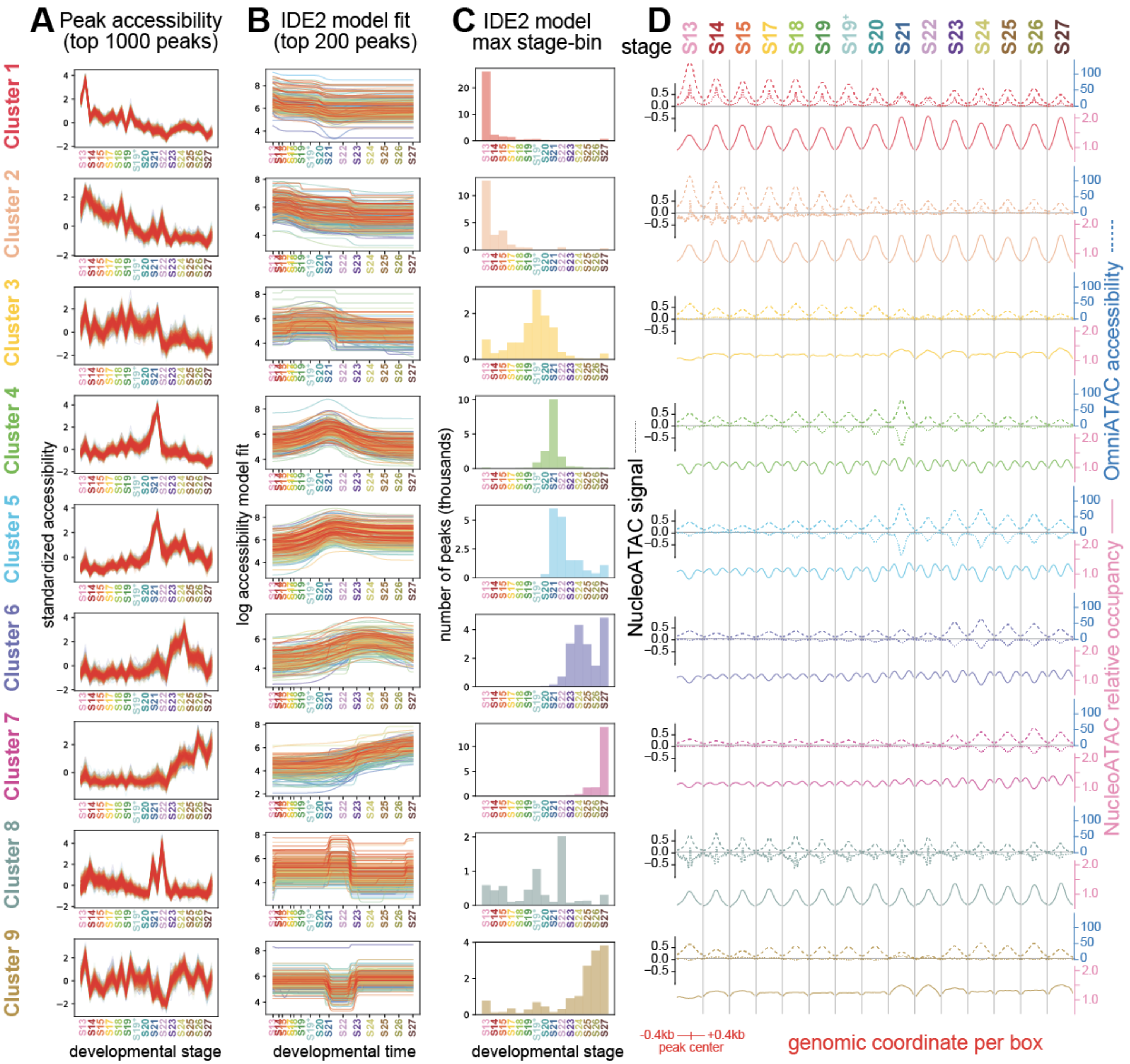

A Peak accessibility (top 1000 peaks)
B IDE2 model fit (top 200 peaks)
C IDE2 model max stage-bin
D
stage
S13 S14 S15 S17 S18 S19 S19⁺ S20 S21 S22 S23 S24 S25 S26 S27
Cluster 1
Cluster 2
Cluster 3
Cluster 4
Cluster 5
Cluster 6
Cluster 7
Cluster 8
Cluster 9
standardized accessibility
log accessibility model fit
number of peaks (thousands)
NucleoATAC signal
OmniATAC accessibility
NucleoATAC relative occupancy
developmental stage
developmental time
developmental stage
-0.4kb +0.4kb
peak center
genomic coordinate per box

#### Supp. Fig. 5.1: Identification and classification of regulatory element clusters

A) Standardized accessibility plot for top 1000 peaks with strongest membership in each of the 9 clusters. Each library is plotted as a separate point along the line plot.

B) ImpulseDE2 model fits for the top 200 peaks with strongest membership in each of the 9 clusters.

C) Histogram of when all peaks in each cluster achieve their maximum accessibility as calculated by IDE2 max fit.

D) Line plots showing Omni-ATAC accessibility (dashed line, blue axis), NucleoATAC signal (dotted line, black axis), and relative NucleoATAC histone occupancy (solid line, pink axis) for peaks in each cluster across time. Each box contains a line plot summarizing each of the three signals at all peaks within that cluster with respect to a given developmental Omni-ATAC-Seq library. The center of each line plot reflects the average signal at the center of all Omni-ATAC-Seq peaks in that cluster at that developmental stage. Signal is visualized at 0.4kb upstream and downstream of peak centers. Within each signal type, the axes across line plots for each cluster are identical.

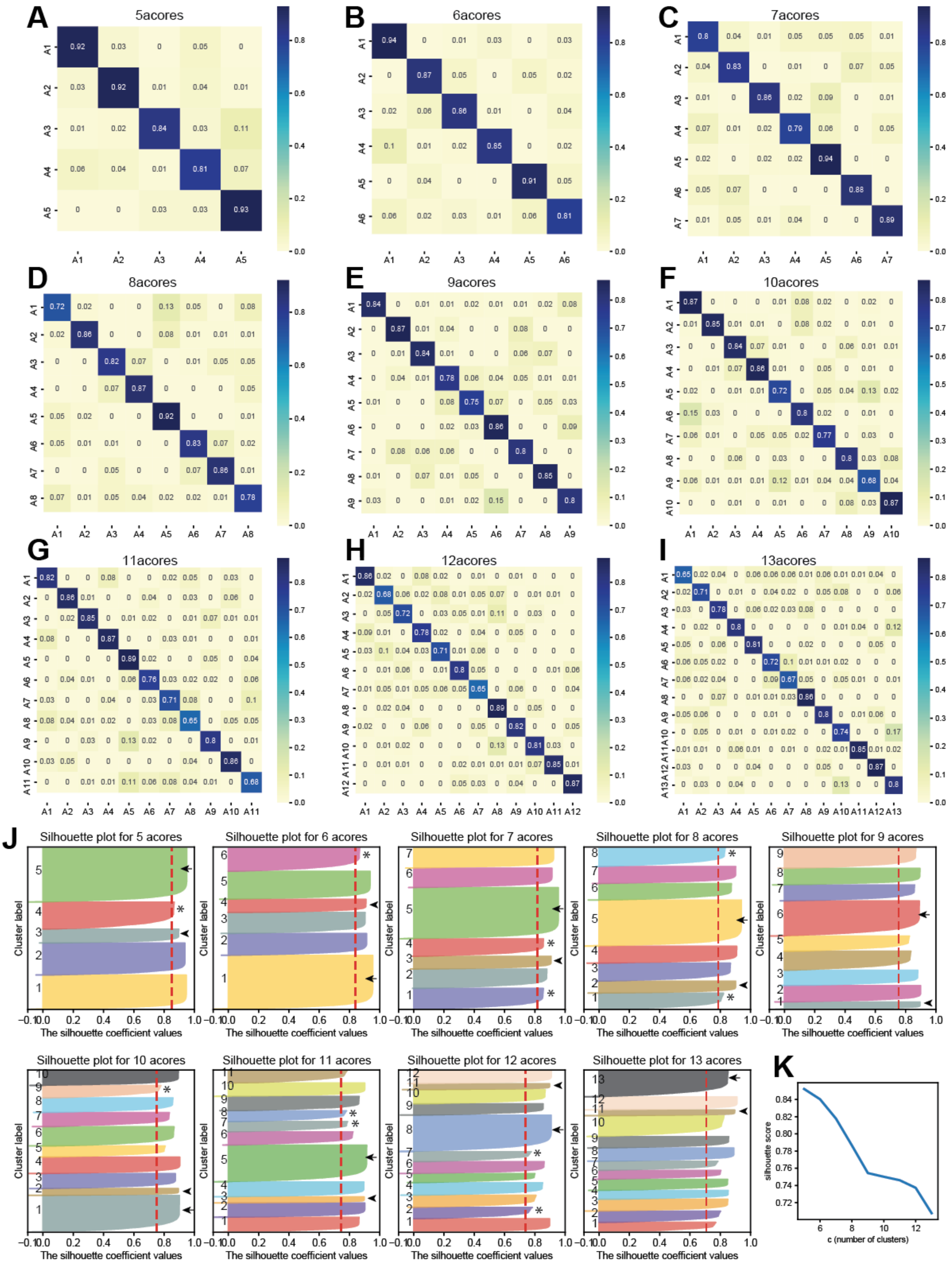
A
5acores
B
6acores
C
7acores
D
8acores
E
9acores
F
10acores
G
11acores
H
12acores
I
13acores
J
Silhouette plot for 5 acores
Silhouette plot for 6 acores
Silhouette plot for 7 acores
Silhouette plot for 8 acores
Silhouette plot for 9 acores
Silhouette plot for 10 acores
Silhouette plot for 11 acores
Silhouette plot for 12 acores
Silhouette plot for 13 acores
Cluster label
The silhouette coefficient values
K
silhouette score
c (number of clusters)

#### Supp. Fig. 5.2: Mfuzz clustering assessment

Note: cluster labels in this plot are for assessment purposes, and do not correspond to cluster labels used in other figures.
A–I) Cluster overlap matrices generated my Mfuzz indicating percentage overlap between clusters.
J) Silhouette plots for different numbers of clusters. Dashed red line marks the average silhouette score. Asterisks mark clusters with marginal silhouettes (silhouettes at or below the average). An arrowhead marks the thinnest silhouette, while an arrow marks the thickest silhouette within each plot. All cluster numbers other than 9 and 13 showed marginal clusters and large contrast between the thinnest and thickest silhouette widths.
K) Mean silhouette score by number of clusters.

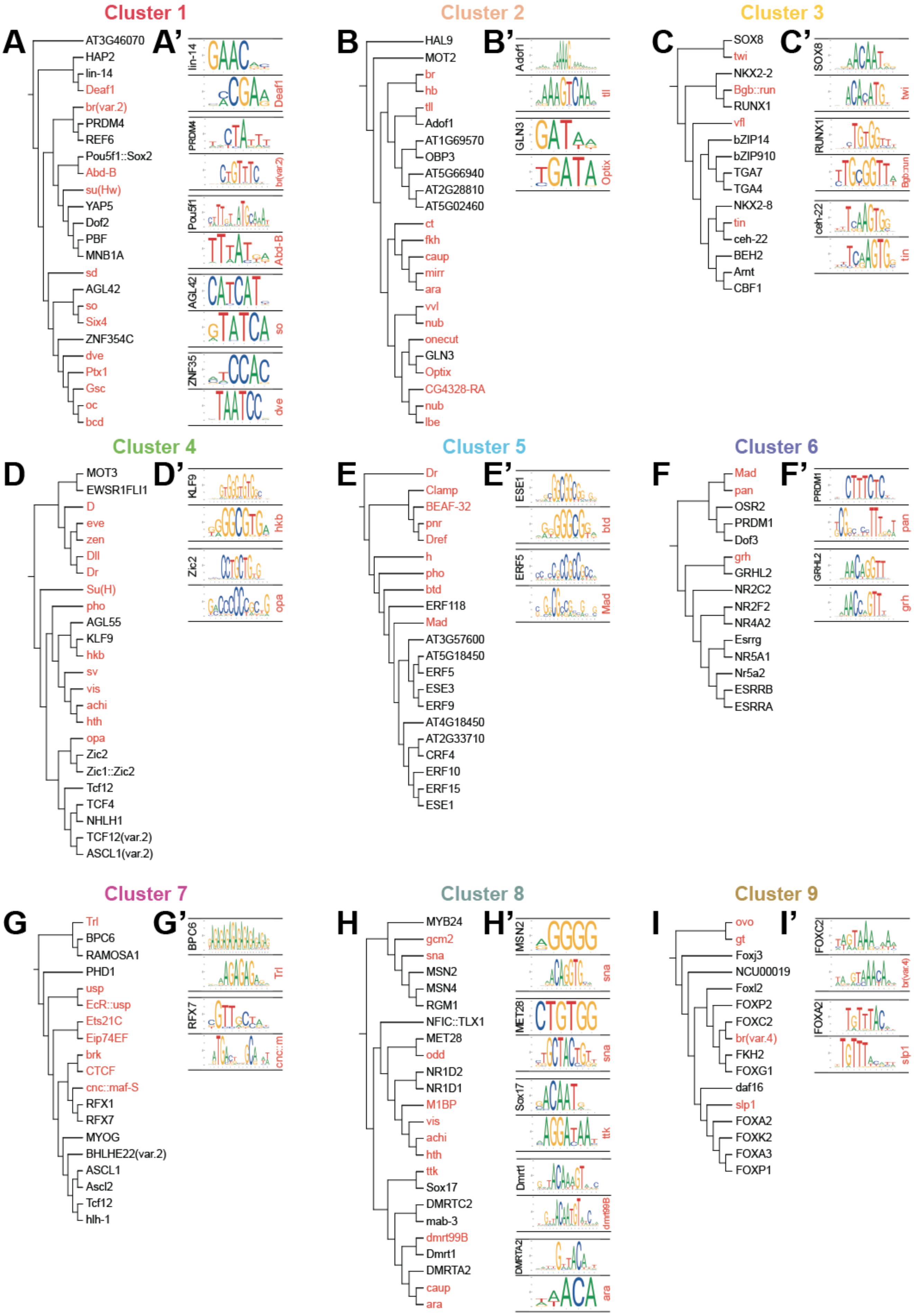

Cluster 1
A
AT3G46070
HAP2
lin-14
Deaf1
br(var.2)
PRDM4
REF6
Pou5f1::Sox2
Abd-B
su(Hw)
YAP5
Dof2
PBF
MNB1A
sd
AGL42
so
Six4
ZNF354C
dve
Ptx1
Gsc
oc
bcd
A'
lin-14
Deaf1
PRDM4
br(var.2)
Pou5f1
Abd-B
AGL42
so
ZNF354C
dve
Cluster 2
B
HAL9
MOT2
br
hb
tll
Adof1
AT1G69570
OBP3
AT5G66940
AT2G28810
AT5G02460
ct
fkh
caup
mirr
ara
vvl
nub
onecut
GLN3
Optix
CG4328-RA
nub
lbe
B'
Adof1
tll
GLN3
Optix
Cluster 3
C
SOX8
twi
NKX2-2
Bgb::run
RUNX1
vfl
bZIP14
bZIP910
TGA7
TGA4
NKX2-8
tin
ceh-22
BEH2
Arnt
CBF1
C'
SOX8
twi
RUNX1
Bgb::run
ceh-22
tin
Cluster 4
D
MOT3
EWSR1FLI1
D
eve
zen
Dll
Dr
Su(H)
pho
AGL55
KLF9
hkb
sv
vis
achi
hth
opa
Zic2
Zic1::Zic2
Tcf12
TCF4
NHLH1
TCF12(var.2)
ASCL1(var.2)
D'
KLF9
hkb
Zic2
opa
Cluster 5
E
Dr
Clamp
BEAF-32
pnr
Dref
h
pho
btd
ERF118
Mad
AT3G57600
AT5G18450
ERF5
ESE3
ERF9
AT4G18450
AT2G33710
CRF4
ERF10
ERF15
ESE1
E'
ESE1
btd
ERF5
Mad
Cluster 6
F
Mad
pan
OSR2
PRDM1
Dof3
grh
GRHL2
NR2C2
NR2F2
NR4A2
Esrrg
NR5A1
Nr5a2
ESRRB
ESRRA
F'
PRDM1
pan
GRHL2
grh
Cluster 7
G
Trl
BPC6
RAMOSA1
PHD1
usp
EcR::usp
Ets21C
Eip74EF
brk
CTCF
cnc::maf-S
RFX1
RFX7
MYOG
BHLHE22(var.2)
ASCL1
Ascl2
Tcf12
hlh-1
G'
BPC6
Trl
RFX7
cnc::m
Cluster 8
H
MYB24
gcm2
sna
MSN2
MSN4
RGM1
NFIC::TLX1
MET28
odd
NR1D2
NR1D1
M1BP
vis
achi
hth
ttk
Sox17
DMRTC2
mab-3
dmrt99B
Dmrt1
DMRTA2
caup
ara
H'
MSN2
sna
MET28
sna
Sox17
ttk
Dmrt1
dmrt99B
DMRTA2
ara
Cluster 9
I
ovo
gt
Foxj3
NCU00019
Foxl2
FOXP2
FOXC2
br(var.4)
FKH2
FOXG1
daf16
slp1
FOXA2
FOXK2
FOXA3
FOXP1
I'
FOXC2
br(var.4)
FOXA2
slp1

#### Supp. Fig. 6: Alignment of motifs in fuzzy clusters

A–I) Dendrograms generated from sequence alignment of JASPAR motifs using the online STAMP tool visualized using Interactive Tree of Life (iTOL). Top 12 most-enriched transcription factor binding sites from the entire JASPAR CORE database are in black for each dendrogram; up to top 12 most-enriched transcription factor binding sites from the *Drosophila* transcription factors in JASPAR CORE are in red for each dendrogram.
A'–I') Select pairs of transcription factor binding sites with apparently similar sequences based on STAMP alignment. All JASPAR CORE TFs are in black; *Drosophila* TFs are in red.

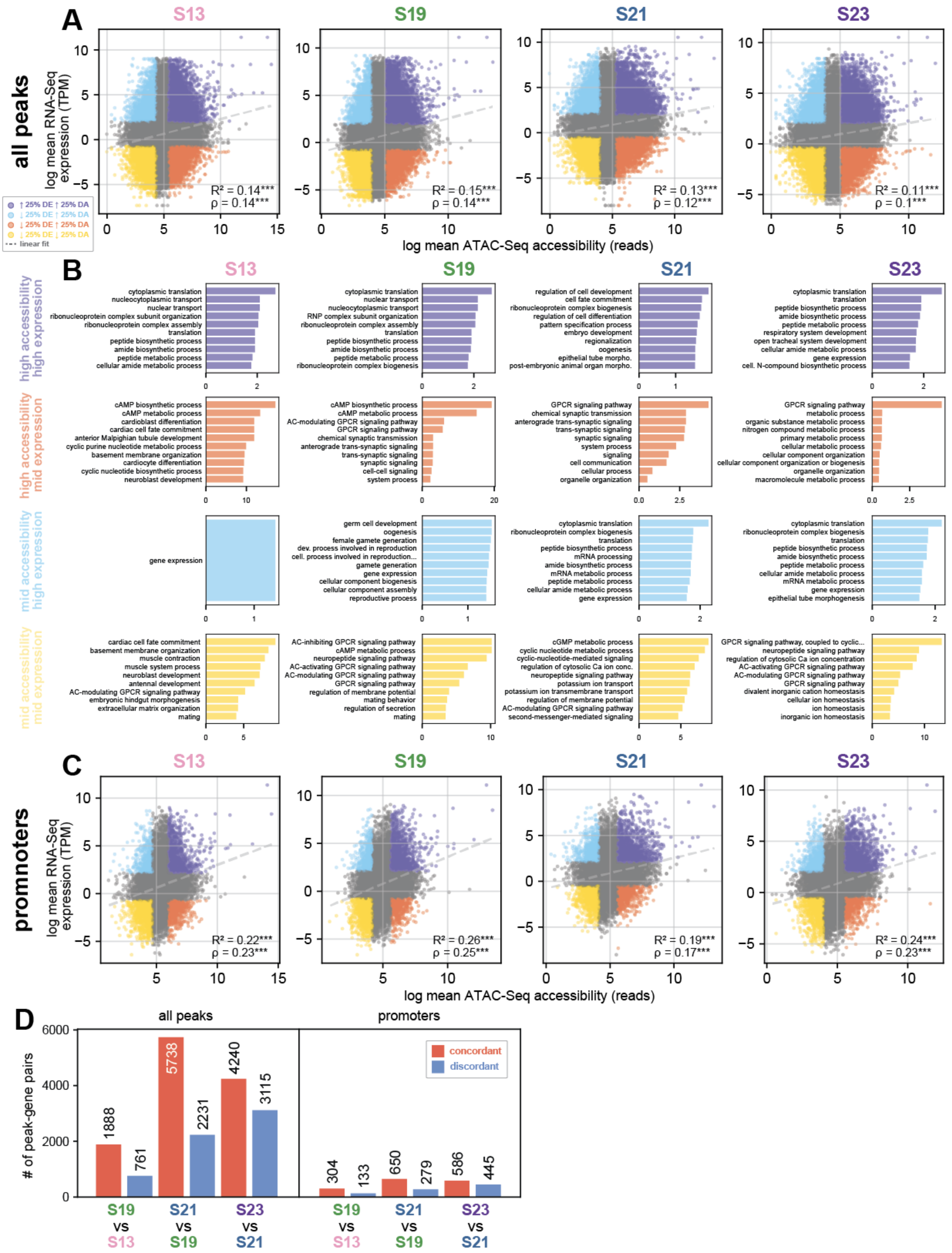
A
B
C
D
S13
S19
S21
S23
all peaks
promoters
log mean RNA-Seq expression (TPM)
log mean ATAC-Seq accessibility (reads)
↑25% DE ↑25% DA
↓25% DE ↑25% DA
↓25% DE ↑25% DA
↓25% DE ↓25% DA
linear fit
R² = 0.14***
ρ = 0.14***
R² = 0.15***
ρ = 0.14***
R² = 0.13***
ρ = 0.12***
R² = 0.11***
ρ = 0.1***
high accessibility high expression
high accessibility mid expression
mid accessibility high expression
mid accessibility mid expression
cytoplasmic translation
nucleocytoplasmic transport
nuclear transport
ribonucleoprotein complex subunit organization
ribonucleoprotein complex assembly
translation
peptide biosynthetic process
amide biosynthetic process
peptide metabolic process
cellular amide metabolic process
RNP complex subunit organization
ribonucleoprotein complex biogenesis
regulation of cell development
cell fate commitment
regulation of cell differentiation
pattern specification process
embryo development
regionalization
oogenesis
epithelial tube morpho.
post-embryonic animal organ morpho.
respiratory system development
open tracheal system development
gene expression
cell. N-compound biosynthetic process
cAMP biosynthetic process
cAMP metabolic process
cardioblast differentiation
cardiac cell fate commitment
anterior Malpighian tubule development
cyclic purine nucleotide metabolic process
basement membrane organization
cardiocyte differentiation
cyclic nucleotide biosynthetic process
neuroblast development
AC-modulating GPCR signaling pathway
GPCR signaling pathway
chemical synaptic transmission
anterograde trans-synaptic signaling
trans-synaptic signaling
synaptic signaling
cell-cell signaling
system process
signaling
cell communication
cellular process
organelle organization
metabolic process
organic substance metabolic process
nitrogen compound metabolic process
primary metabolic process
cellular metabolic process
cellular component organization
cellular component organization or biogenesis
macromolecule metabolic process
germ cell development
female gamete generation
dev. process involved in reproduction
cell. process involved in reproduction...
gamete generation
cellular component biogenesis
cellular component assembly
reproductive process
mRNA processing
mRNA metabolic process
epithelial tube morphogenesis
cardiac cell fate commitment
muscle contraction
muscle system process
antennal development
embryonic hindgut morphogenesis
extracellular matrix organization
mating
AC-inhibiting GPCR signaling pathway
neuropeptide signaling pathway
AC-activating GPCR signaling pathway
regulation of membrane potential
mating behavior
regulation of secretion
cGMP metabolic process
cyclic nucleotide metabolic process
cyclic-nucleotide-mediated signaling
regulation of cytosolic Ca ion conc.
potassium ion transport
potassium ion transmembrane transport
second-messenger-mediated signaling
GPCR signaling pathway, coupled to cyclic...
regulation of cytosolic Ca ion concentration
divalent inorganic cation homeostasis
cellular ion homeostasis
ion homeostasis
inorganic ion homeostasis
R² = 0.22***
ρ = 0.23***
R² = 0.26***
ρ = 0.25***
R² = 0.19***
ρ = 0.17***
R² = 0.24***
ρ = 0.23***
### of peak-gene pairs
concordant
discordant
1888
761
5738
2231
4240
3115
304
133
650
279
586
445
S19 vs S13
S21 vs S19
S23 vs S21

#### Supp. Fig. 7: Correlation between accessibility and gene expression at individual timepoints

A) Correlation between log mean RNA-Seq TPM and log mean Omni-ATAC-Seq accessibility at all peaks in each developmental stage. Peaks colored based on highest and lowest quartiles of expression and accessibility. Dotted line represents a linear fit to all data in the plot. Pearson correlation $R^2$ and Spearman correlation ρ for all points are displayed in each plot.

B) GO-term enrichment for gene-peak pairs with high accessibility and high expression (purple), high accessibility and medium expression (coral), medium accessibility and high expression (blue), and medium accessibility and medium expression (yellow), for all peaks. The threshold for “high” expression and accessibility were all values above the 75th percentile, while “mid” expression and accessibility were all values below the 25th percentile. Gene lists were extracted based on the nearest Mikado gene to each peak, and orthologous *Drosophila* names were assigned to genes using OrthoFinder.

C) Correlation between log mean RNA-Seq TPM and log mean Omni-ATAC-Seq accessibility at promoter peaks.

D) Quantification of number of significant peak-gene pairs at each time point comparison in Fig. 7. Left plot quantifies number of concordant and discordant peaks across all peak-gene pairs, while right plot quantifies peak-gene pairs that were assigned as promoters.

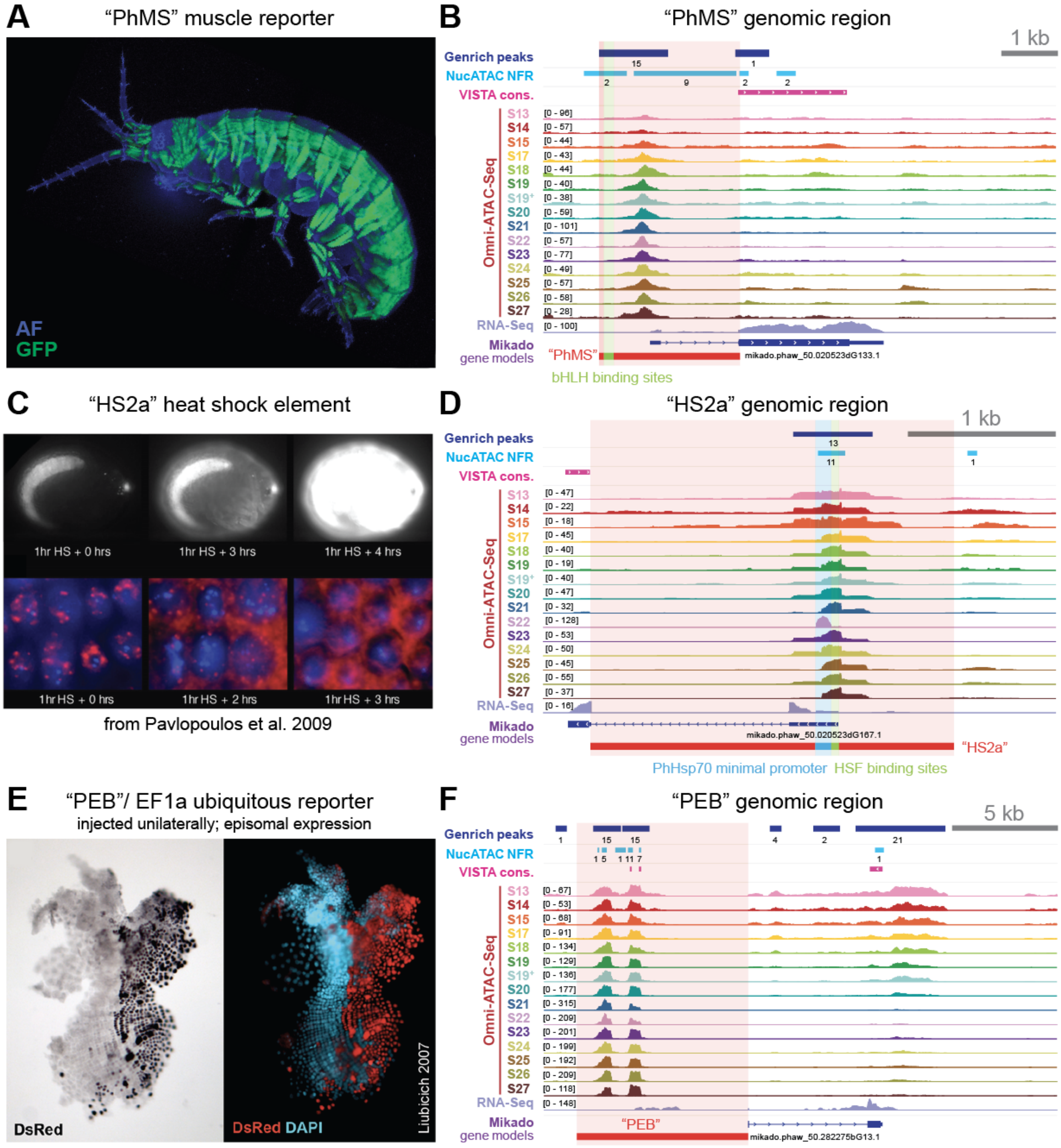
A "PhMS" muscle reporter
AF
GFP
B "PhMS" genomic region
1 kb
Genrich peaks
NucATAC NFR
VISTA cons.
Omni-ATAC-Seq
S13 S14 S15 S17 S18 S19 S19* S20 S21 S22 S23 S24 S25 S26 S27
RNA-Seq
Mikado gene models
"PhMS"
mikado.phaw_50.020523dG133.1
bHLH binding sites
C "HS2a" heat shock element
1hr HS + 0 hrs
1hr HS + 3 hrs
1hr HS + 4 hrs
1hr HS + 0 hrs
1hr HS + 2 hrs
1hr HS + 3 hrs
from Pavlopoulos et al. 2009
D "HS2a" genomic region
1 kb
Genrich peaks
NucATAC NFR
VISTA cons.
Omni-ATAC-Seq
RNA-Seq
Mikado gene models
mikado.phaw_50.020523dG167.1
"HS2a"
PhHsp70 minimal promoter
HSF binding sites
E "PEB"/ EF1a ubiquitous reporter
injected unilaterally; episomal expression
DsRed
DsRed DAPI
Liubicich 2007
F "PEB" genomic region
5 kb
Genrich peaks
NucATAC NFR
VISTA cons.
Omni-ATAC-Seq
RNA-Seq
Mikado gene models
"PEB"
mikado.phaw_50.282275bG13.1

##### Supp. Fig. 8.1: Omni-ATAC-Seq recovers previously identified regulatory elements

A) Expression of the PhMS reporter visualized using GFP in a juvenile *Parhyale*.
B) The PhMS genomic region. A strong Genrich peak overlaps with a predicted cluster of b-HLH binding sites.
C) Heat shock expression images from a transgenic *Parhyale* embryo carrying the HS2a element.
D) The HS2a genomic region. A peak covers the PhHsp70 minimal promoter and heat shock factor binding sites.
E) Expression of the PEB reporter, visualized in a dissected embryo using a DsRed antibody. Injected embryos showed nearly full expression of this reporter in each germ layer (Liubicich 2007 PhD thesis).
F) The PEB genomic region. Two strong peaks overlap with the 5’ end of this reporter.

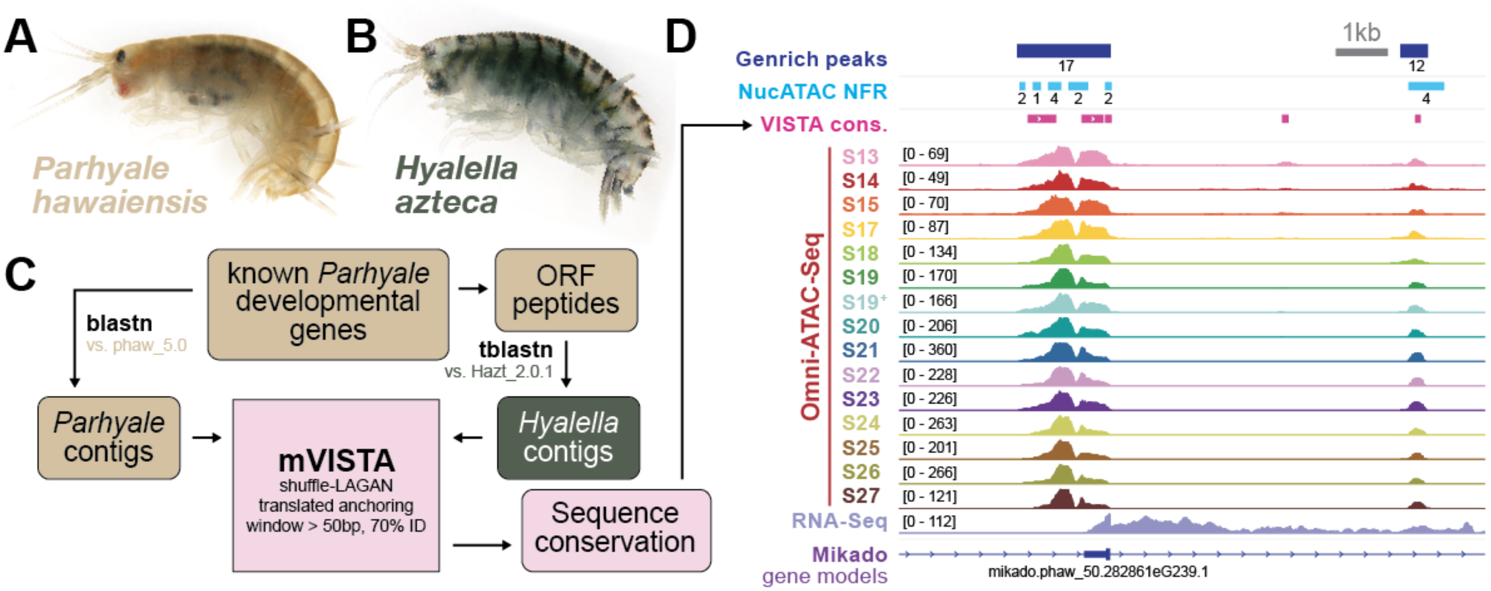

A
Parhyale hawaiensis
B
Hyalella azteca
C
known Parhyale developmental genes
ORF peptides
blastn
vs. phaw_5.0
tblastn
vs. Hazt_2.0.1
Parhyale contigs
mVISTA
shuffle-LAGAN
translated anchoring
window > 50bp, 70% ID
Hyalella contigs
Sequence conservation
D
1kb
Genrich peaks
NucATAC NFR
VISTA cons.
Omni-ATAC-Seq
S13
S14
S15
S17
S18
S19
S19+
S20
S21
S22
S23
S24
S25
S26
S27
RNA-Seq
Mikado gene models
mikado.phaw_50.282861eG239.1

#### Supp. Fig. 8.2: VISTA homology identification between *Parhyale* and *Hyalella*

A) Adult female *Parhyale hawaiensis*.
B) Adult female *Hyalella azteca*.
C) Pipeline for identification of orthologous genomic regions between *Hyalella* and *Parhyale* for VISTA sequence conservation analysis.
D) Visualization of Genrich peaks, NucleoATAC nucleosome-free regions (NFRs), VISTA conservation, Omni-ATAC-Seq signal, RNA-Seq signal, and Mikado gene models at the putative Sp69 promoter peak.

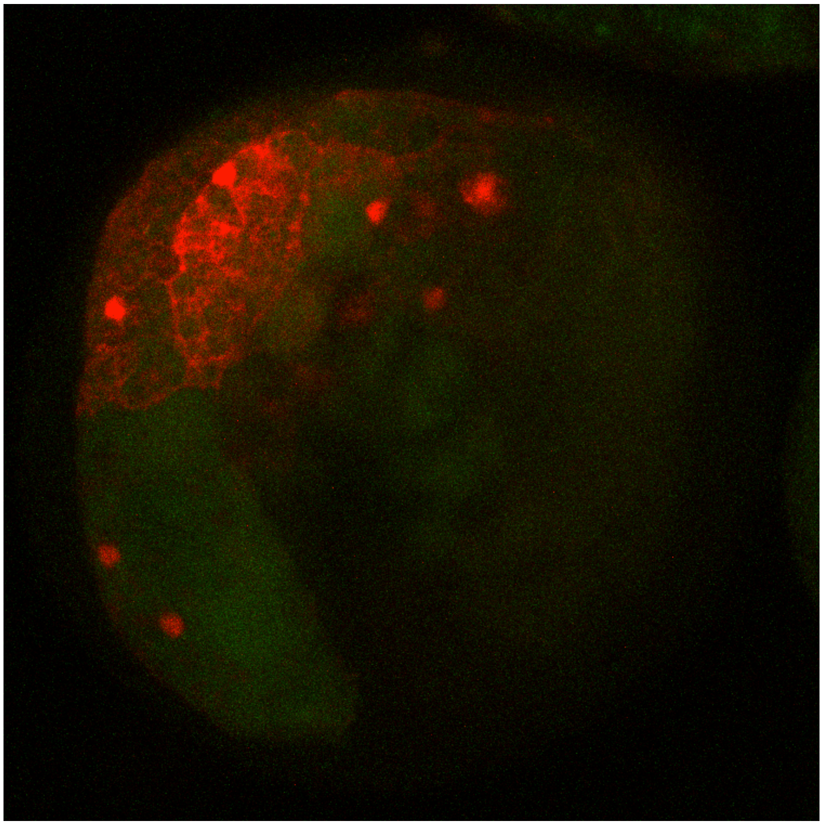

**Supplementary Video 1: Yolk cells labeled by the Hsp70-p2 reporter**

Max intensity projection; frames taken 5 minutes apart.

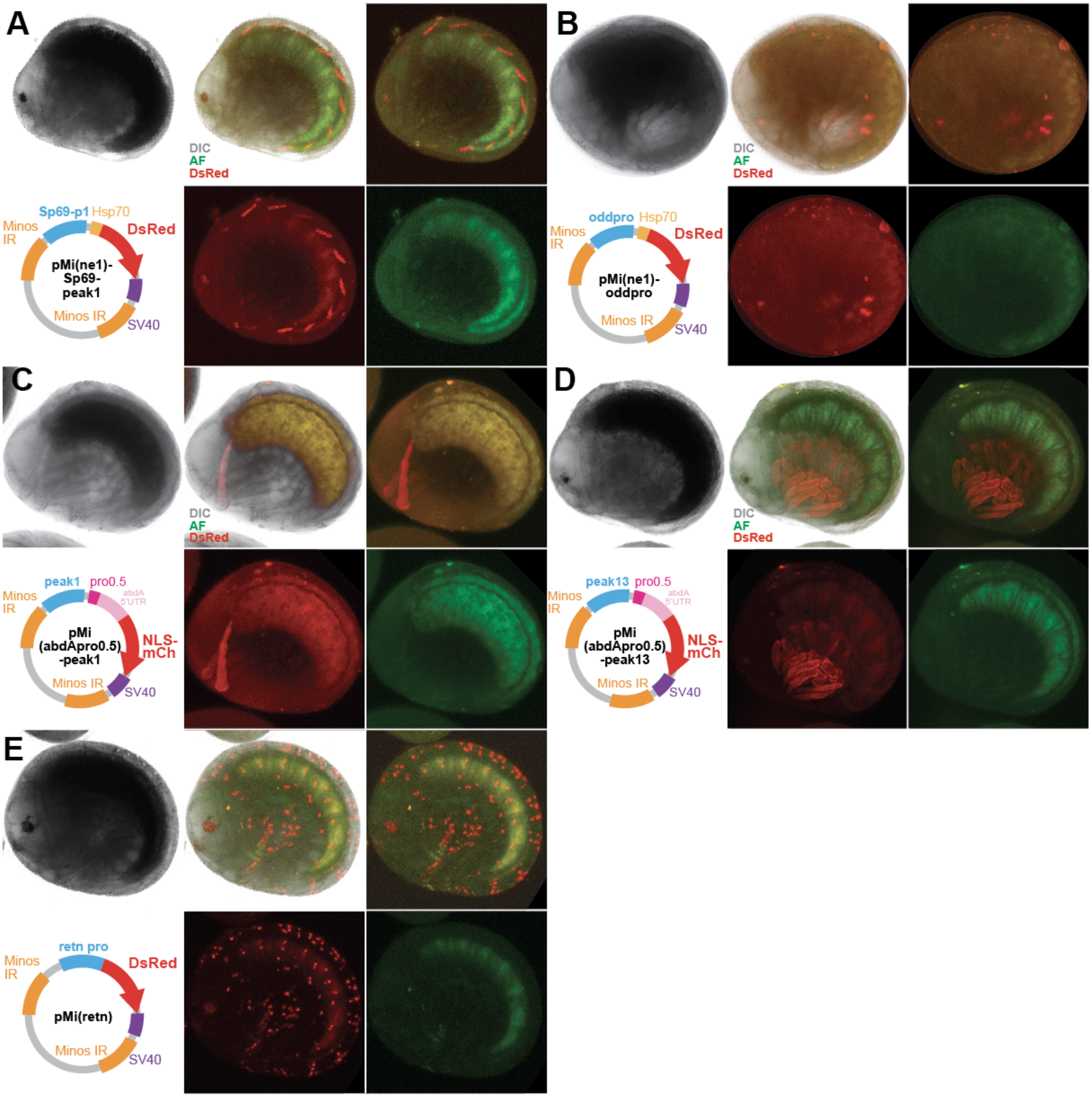

A
DIC
AF
DsRed
Sp69-p1
Hsp70
Minos IR
DsRed
pMi(ne1)-Sp69-peak1
Minos IR
SV40
B
DIC
AF
DsRed
oddpro
Hsp70
Minos IR
DsRed
pMi(ne1)-oddpro
Minos IR
SV40
C
DIC
AF
DsRed
peak1
pro0.5
abdA 5'UTR
Minos IR
pMi (abdApro0.5) -peak1
NLS-mCh
Minos IR
SV40
D
DIC
AF
DsRed
peak13
pro0.5
abdA 5'UTR
Minos IR
pMi (abdApro0.5) -peak13
NLS-mCh
Minos IR
SV40
E
retn pro
Minos IR
DsRed
pMi(retn)
Minos IR
SV40

#### Supp. Fig. 9: Spontaneous, non-reproducible expression patterns observed in Minos assays

A) Expression observed in one embryo injected with a plasmid containing the PhHsp70 minimal promoter and the Sp69-p1 peak.

B) Expression observed in one embryo injected with a plasmid containing proximal promoter peak for the odd-skipped gene along with an Hsp80 minimal promoter.

C) Expression observed in one embryo injected with a plasmid containing a putative minimal promoter for the abd-A gene and an upstream peak (abd-A-peak1).

D) Expression observed in one embryo injected with a plasmid containing a putative minimal promoter for the abd-A gene and an upstream peak (abd-A-peak13).

E) Expression observed in one embryo injected with a plasmid containing the putative *Parhyale-retn* promoter.

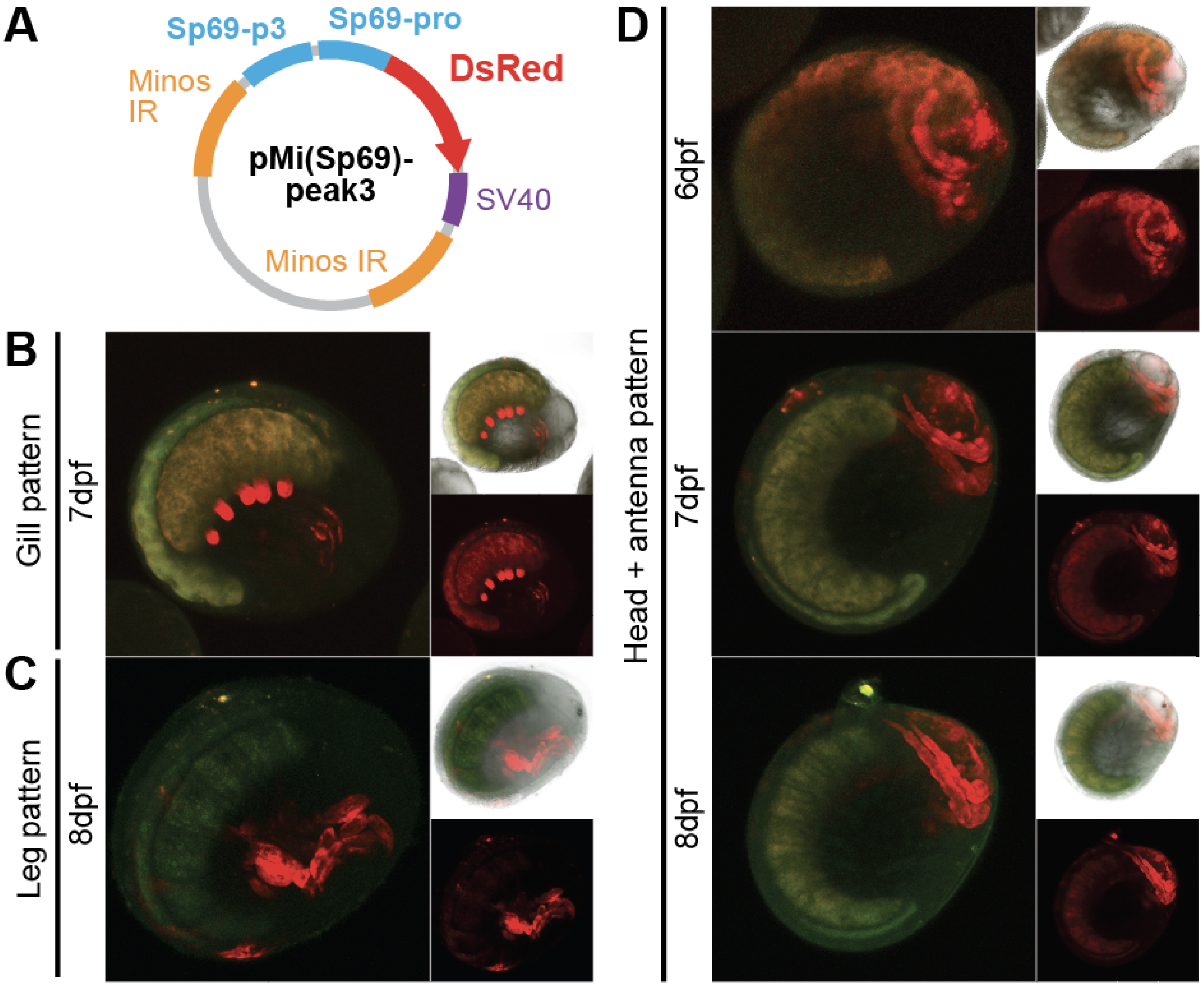
A
Sp69-p3
Sp69-pro
DsRed
Minos IR
pMi(Sp69)-peak3
SV40
Minos IR
B
Gill pattern
7dpf
C
Leg pattern
8dpf
D
Head + antenna pattern
6dpf
7dpf
8dpf

#### Supp. Fig. 10: Expression observed from Sp69-p3 reporter

A) Plasmid map of the pMi(Sp69)-peak3 construct. The construct contains both the putative promoter peak of Sp69, along with a distal peak (peak3).
B) Gill and claw expression observed in one embryo injected with this plasmid.
C) Leg expression observed in one embryo injected with this plasmid.
D) Head and antennal expression patterns observed in 5 embryos injected with this plasmid; embryo with strongest expression pattern shown.

**Supplementary Table 1: Illumina sequencing indices for Omni-ATAC libraries**

| Sample ID | i7 (4000) | i5 (4000) PE | i5 (4000) SR Flowcell/MiSeq |
|---|---|---|---|
| S13A | TACAGAGC | CTTGGATG | CATCCAAG |
| S13B | AAGCGTTC | GGAAGGAT | ATCCTTCC |
| S14A | AGTGACCT | TTCTCTCG | CGAGAGAA |
| S14B | ACCATAGG | AAGTCCGT | ACGGACTT |
| S15A | CTCGAACA | ATGGTCCA | TGGACCAT |
| S15B | TGCGTAAC | TGTCCAGA | TCTGGACA |
| S17A | AACAGTCC | GATTGCTC | GAGCAATC |
| S17B | CTAAGACC | GTTGTAGC | GCTACAAC |
| S18A | TGTTCCGT | GTGAATCC | GGATTCAC |
| S18B | ACTCAACG | ACTCCATC | GATGGAGT |
| S19A | GCCTTCTT | TGAGGTGT | ACACCTCA |
| S19B | TGCTCTAC | TCTTGACG | CGTCAAGA |
| S19plusA | GTACCACA | GATACTGG | CCAGTATC |
| S19plusB | GCATTGGT | ATCTTCGG | CCGAAGAT |
| S20A | CTGTGGTA | ACATTGCG | CGCAATGT |
| S20B | TTACCGAC | GTTGTTCG | CGAACAAC |
| S21A | TGACCGTT | CATGGAAC | GTTCCATG |
| S21B | GCTAAGGA | AACGTGGA | TCCACGTT |
| S22A | TAGCCATG | CTGCACTT | AAGTGCAG |
| S22B | ACTTGGCT | GTAGCATC | GATGCTAC |
| S23A | TGTCACAC | GTGCTTAC | GTAAGCAC |
| S23B | GACACAGT | CAACCTAG | CTAGGTTG |
| S24A | GTGGTATG | ATCTCGCT | AGCGAGAT |
| S24B | ATGCGCTT | GGACCTAT | ATAGGTCC |
| S25A | AGGAACAC | ACGTTACC | GGTAACGT |
| S25B | TGGAAGCA | TACTGCGT | ACGCAGTA |
| S26A | GTCGTTAC | GGTACTAC | GTAGTACC |
| S26B | GAACCTTC | AGCTTGAG | CTCAAGCT |
| S27A | TATGACCG | TGTGAAGC | GCTTCACA |
| S27B | AACGCACA | CGTTATGC | GCATAACG |

### Supplementary Table 2: Manual annotation coordinates of 49 selected RACE genes

| gene_name | IGV_address | Strand | pro_peak_address | exon1_address | exon-1_address | Trinity fragments? | Fragment type |
|---|---|---|---|---|---|---|---|
| **AbdominalBisoform1cloneJ7b11** | phaw_50.282695a:24623698-24688129 | + | phaw_50.282695a:24622599-24624450 | phaw_50.282695a:24,623,624-24,624,479 | phaw_50.282695a:24,687,725-24,692,898 | Yes | Both |
| **abdomminalAisoform2cloneK313** | phaw_50.282695a:25629814-25736948 | + | phaw_50.282695a:25628608-25630296 | phaw_50.282695a:25,627,088-25,630,363 | phaw_50.282695a:25,736,469-25,741,275 | Yes | Both |
| **Antennapedia5racecloneL18** | phaw_50.282695a:26532519-26613138 | + | phaw_50.282695a:26532243-26532820 | phaw_50.282695a:26,532,505-26,533,152 | phaw_50.282695a:26,611,100-26,613,267 | Yes | Both |
| **ash26F** | phaw_50.283875a:11478705-11507717 | + | phaw_50.283875a:11477924-11479069 | phaw_50.283875a:11,478,677-11,480,112 | phaw_50.283875a:11,507,570-11,508,560 | Yes | Both |
| **betacateninmRNAcompletecds** | phaw_50.282639b:6484601-6538431 | + | phaw_50.282639b:6451035-6452617 | phaw_50.282639b:6,451,925-6,452,777 | phaw_50.282639b:6,537,263-6,541,062 | Yes | Both |
| **col** | phaw_50.283865b:130292-293921 | + | phaw_50.283865b:127661-128803 | phaw_50.283865b:130,178-130,591 | phaw_50.283865b:296,755-299,392 | Yes | Both |
| **DeformedcDNAcloneFLb2** | phaw_50.282695a:27062438-27152191 | + | phaw_50.282695a:27061893-27063246 | phaw_50.282695a:27,062,404-27,063,447 | phaw_50.282695a:27,151,053-27,152,222 | Yes | Exon |
| **deltaproteinmRNAcompletecds** | phaw_50.015400b:11871425-11889172 | + | phaw_50.015400b:11870888-11871925 | phaw_50.015400b:11,871,436-11,871,564 | phaw_50.015400b:11,890,317-11,891,420 | Yes | Intron |
| **distallessEarlymRNAcompletecds** | phaw_50.282654b:30641259-30681459 | + | phaw_50.282654b:30640553-30642307 | phaw_50.282654b:30,641,130-30,641,834 | phaw_50.282654b:30,685,403-30,687,854 | Yes | Both |
| **dpp** | phaw_50.015400a:5773544-5790898 | + | phaw_50.015400a:5772703-5774732 | phaw_50.015400a:5,773,547-5,774,013 | phaw_50.015400a:5,789,594-5,799,808 | Yes | Both |
| **engrailed1** | phaw_50.283864a:3322422-3404220 | + | phaw_50.283864a:3320702-3323202 | phaw_50.283864a:3,321,545-3,323,280 | phaw_50.283864a:3,403,484-3,411,112 | Yes | Both |
| **engrailed2** | phaw_50.283864a:3774988-3800052 | - | phaw_50.283864a:3799353-3800567 | phaw_50.283864a:3,798,531-3,800,059 | phaw_50.283864a:3,757,132-3,758,228 | Yes | Intron |
| **eve2** | phaw_50.283028b:9254180-9262623 | - | phaw_50.283028b:9262111-9262983 | phaw_50.283028b:9,262,071-9,262,616 | phaw_50.283028b:9,250,289-9,256,607 | Yes | Exon |
| **extradenticleproteinexdgene** | phaw_50.283468a:8063095-8076889 | + | phaw_50.283468a:7799208-7800212 | phaw_50.283468a:7,799,472-7,800,056 | phaw_50.283468a:8,097,516-8,101,622 | Yes | Both |
| **forkheadORF** | phaw_50.000135a:2643865-2644971 | - | phaw_50.000135a:2649799-2650196 | phaw_50.000135a:2,649,559-2,649,662 | phaw_50.000135a:2,642,441-2,645,164 | Yes | Exon |
| **hes4** | phaw_50.282654b:31719607-31734841 | + | phaw_50.282654b:31719410-31720007 | phaw_50.282654b:31,719,607-31,719,748 | phaw_50.282654b:31,732,919-31,736,371 | Yes | Both |
| **homothoraxproteinhthgene** | phaw_50.283815b:16274653-16574076 | - | phaw_50.283815b:16604672-16607172 | phaw_50.283815b:16,605,602-16,606,372 | phaw_50.283815b:16,168,073-16,170,044 | Yes | Both |
| **KNIRPS1kni1mRNAcompletecds** | phaw_50.283866:9026088-9035313 | + | phaw_50.283866:9025431-9026438 | phaw_50.283866:9,026,082-9,026,813 | phaw_50.283866:9,034,022-9,037,291 | Yes | Both |
| **KNIRPS2kni2mRNAcompletecds** | phaw_50.283866:9982901-10018092 | - | phaw_50.283866:10018846-10019492 | phaw_50.283866:10,017,879-10,018,098 | phaw_50.283866:9,982,950-9,984,746 | Yes | Exon |
| **Kruppel5raceclone5** | phaw_50.004430:12895048-13017154 | + | phaw_50.004430:12893620-12894503 | phaw_50.004430:12,893,830-12,895,369 | phaw_50.004430:13,014,676-13,017,950 | Yes | Both |
| **notchproteinmRNAcompletecds** | phaw_50.283815c:31626720-31812600 | - | phaw_50.283815c:31812314-31813325 | phaw_50.283815c:31,812,341-31,812,602 | phaw_50.283815c:31,625,065-31,629,214 | No | None |
| **odd1** | phaw_50.000289b:10469078-10470818 | - | phaw_50.000289b:10470131-10471455 | phaw_50.000289b:10,470,089-10,470,823 | phaw_50.000289b:10,469,064-10,470,057 | Yes | Exon |
| **odd3** | phaw_50.000289b:9790462-9792439 | + | phaw_50.000289b:9789912-9790964 | phaw_50.000289b:9,790,460-9,790,970 | phaw_50.000289b:9,791,530-9,795,846 | Yes | Exon |
| **odd5** | phaw_50.000289b:10003623-10007578 | - | phaw_50.000289b:10007157-10007887 | phaw_50.000289b:10,007,151-10,007,894 | phaw_50.000289b:10,001,868-10,005,067 | Yes | Both |
| **opa1consensus** | phaw_50.000135f:28969563-29076765 | + | phaw_50.000135f:28968467-28970375 | phaw_50.000135f:28,969,544-28,970,470 | phaw_50.000135f:29,072,172-29,076,827 | Yes | Both |
| **optixmRNAcompletecds** | phaw_50.283817f:8745320-8792652 | + | phaw_50.283817f:8744687-8746467 | phaw_50.283817f:8,745,412-8,746,075 | phaw_50.283817f:8,788,522-8,794,388 | Yes | Both |
| **Par6** | phaw_50.283823c:3497885-3548486 | - | phaw_50.283823c:3548118-3548740 | phaw_50.283823c:3,548,186-3,548,556 | phaw_50.283823c:3,492,657-3,499,147 | Yes | Both |
| **Pax371proteinmRNAcompletecds** | phaw_50.000214e:5044320-5078499 | - | phaw_50.000214e:5078053-5078983 | phaw_50.000214e:5,078,120-5,078,455 | phaw_50.000214e:5,042,664-5,047,700 | Yes | Both |
| **pdm** | phaw_50.283869d:4505837-4506191 | + | phaw_50.283869d:4420193-4421142 | phaw_50.283869d:4,420,800-4,421,678 | phaw_50.283869d:4,506,146-4,509,682 | Yes | Both |
| **PhDIIlate1** | phaw_50.282654b:30431942-30563945 | + | phaw_50.282654b:30429539-30431011 | phaw_50.282654b:30,430,577-30,432,821 | phaw_50.282654b:30,563,597-30,569,176 | Yes | Both |
| **PhDIIlate2** | phaw_50.282654b:30746554-30806154 | + | phaw_50.282654b:30745371-30747232 | phaw_50.282654b:30,746,377-30,747,054 | phaw_50.282654b:30,805,338-30,813,233 | Yes | Both |
| **pnt3RACE** | phaw_50.282861b:41123311-41125508 | - | phaw_50.282861b:41129249-41130109 | phaw_50.282861b:41,128,992-41,129,723 | phaw_50.282861b:41,112,188-41,123,884 | Yes | Exon |
| **proboscipedia5racecloneI192** | phaw_50.282695a:27659540-27817538 | + | phaw_50.282695a:27657466-27658309 | phaw_50.282695a:27,657,840-27,659,994 | phaw_50.282695a:27,818,947-27,821,498 | Yes | Both |
| **prosperoproteinmRNApartialcds** | phaw_50.283875a:6013042-6035299 | + | phaw_50.283875a:5998087-5999786 | phaw_50.283875a:5,998,945-6,001,387 | phaw_50.283875a:6,034,691-6,038,907 | Yes | Intron |
| **rho5RACEvariant1** | phaw_50.000203b:25395477-25496923 | + | phaw_50.000203b:25395456-25395780 | phaw_50.000203b:25,395,472-25,395,662 | phaw_50.000203b:25,496,406-25,503,322 | Yes | Both |
| **rho5RACEvariant2** | phaw_50.000203b:25265811-25496923 | + | phaw_50.000203b:25265169-25265874 | phaw_50.000203b:25,264,762-25,265,904 | phaw_50.000203b:25,496,405-25,503,316 | Yes | Both |
| **runt2** | phaw_50.283028b:1783456-1810535 | - | phaw_50.283028b:1809649-1811397 | phaw_50.283028b:1,809,475-1,810,690 | phaw_50.283028b:1,776,166-1,784,663 | Yes | Exon |
| **scallopedproteinsdgene** | phaw_50.007301a:763757-786495 | - | phaw_50.007301a:920698-921986 | phaw_50.007301a:921,178-921,537 | phaw_50.007301a:762,031-762,973 | Yes | Both |
| **scratchscrtmRNAcompletecds** | phaw_50.283869f:2146785-2165201 | - | phaw_50.283869f:2270072-2271735 | phaw_50.283869f:2,270,060-2,271,025 | phaw_50.283869f:2,146,761-2,150,484 | Yes | Both |
| **Sexcombsreduced5racecloneE17c1** | phaw_50.282695a:26769068-26862063 | + | phaw_50.282695a:26768486-26770008 | phaw_50.282695a:26,861,434-26,862,081 | phaw_50.282695a:26,768,947-26,770,207 | Yes | Both |
| **shortgastrulationproteinmRNAcompletecds** | phaw_50.015400b:7523582-7867524 | + | phaw_50.015400b:7522832-7524249 | phaw_50.015400b:7,523,573-7,525,453 | phaw_50.015400b:7,866,925-7,867,874 | Yes | Both |
| **slp1** | phaw_50.283811:7718478-7720485 | + | phaw_50.283811:7718255-7718898 | phaw_50.283811:7,718,433-7,720,788 | phaw_50.283811:7,725,774-7,731,670 | Yes | Both |
| **snail2proteinmRNApartialcds** | phaw_50.283864b:718097-721779 | + | phaw_50.283864b:713312-713863 | phaw_50.283864b:713,632-713,751 | phaw_50.283864b:717,219-721,810 | Yes | Both |
| **snail3proteinmRNApartialcds** | phaw_50.283864b:30526-33704 | + | phaw_50.283864b:25948-26939 | phaw_50.283864b:26,453-26,990 | phaw_50.283864b:29,154-33,680 | Yes | Both |
| **Sp69protein** | phaw_50.282861e:14546239-14663576 | + | phaw_50.282861e:14544573-14546375 | phaw_50.282861e:14,545,862-14,546,358 | phaw_50.282861e:14,667,910-14,675,477 | Yes | Both |
| **spi5RACElargefragment3RACE** | phaw_50.000081a:2326730-2344027 | + | phaw_50.000081a:2243466-2244527 | phaw_50.000081a:2,243,861-2,244,179 | phaw_50.000081a:2,347,070-2,349,001 | Yes | Both |
| **SuH** | phaw_50.283826:2280509-2340463 | - | phaw_50.283826:2416373-2417483 | phaw_50.283826:2,416,066-2,416,895 | phaw_50.283826:2,270,773-2,276,731 | Yes | Exon |
| **unc4NPhsequence** | phaw_50.000135e:7542170-7621463 | + | phaw_50.000135e:7541550-7542771 | phaw_50.000135e:7,542,144-7,542,554 | phaw_50.000135e:7,644,646-7,649,912 | Yes | Exon |
| **wnty** | phaw_50.283866:2773623-3090108 | - | phaw_50.283866:3089459-3090541 | phaw_50.283866:3,089,558-3,090,096 | phaw_50.283866:2773623-3090108 | Yes | Both |

#### Supplementary Table 3: Software version numbers

| Software name | Version | Usage |
|---|---|---|
| bamtools | 2.5.1 | Bam file manipulations |
| bedtools | 2.28.0 or 2.30.0 | Bed file manipulations |
| Bowtie2 | 2.3.0 or 2.3.4.1 | Read alignment |
| BUSCO | 3.0.2 | Transcriptome and genome completion evaluation |
| cutadapt | 2.4 | Removing sequencing adapters from reads |
| deeptools | 3.3.1 | Visualization of data genome-wide |
| DESeq2 (R) | 1.34.0 | Differential accessibility/ expression analyses in pairwise comparisons |
| eggNOG-mapper | 2.0.5 | Automated gene function assignment |
| FASTQC | 0.11.7 | Library quality assessment |
| Genrich | 0.6 | Omni-ATAC peak calling |
| GMAP | 2020-11-20 | Aligning transcriptomes to genome for Mikado analyses |
| HISAT2 | 2.1.0 | Aligning RNA_Seq reads to genome |
| igvtools | 2.3.98 | File format conversions for viewing in IGV |
| ImpulseDE2 (R) | 0.99.10 | Performing IDE2 analyses |
| JASPAR2020 (R) | 0.99.10 | Extracting PWMs from JASPAR |
| kallisto | 0.43.1 | Read abundance estimation for RNA-Seq |
| Mikado | 2.3.0 | Mikado transcriptome merging |
| minimap2 | 2.18-r1052-dirty | Aligning long-read Nanopore sequences to *Parhyale* genome |
| NucleoATAC | 0.2.1 | Prediction of nucleosome positions using Omni-ATAC data |
| OrthoFinder | 2.5.4 | Orthology assignment between *Parhyale* and *Drosophila* |
| Portcullis | 1.1.2 | Identifying valid splice junctions from RNA-Seq read data |
| Python | 3.7 or 3.8.8 | Interfacing with Jupyter notebooks and running Python scripts on a SLURM scheduler |
| R | 4.1.1 or 4.0.3 or 3.8 | Performing IDE2, DESeq2 analyses; converting JASPAR files to RGT-HINT format |
| RGT-HINT | 0.13.1 | Inference of transcription factor footprints |
| StringTie | 2.1.7 | Assembly of StringTie2 L and StringTie2 SL transcriptomes |
| Transdecoder | 5.0.2 | Generating peptide sequences for OrthoFinder, Mikado, and other analyses |
| trim_galore | 0.4.4 | Trimming reads used in all analyses |
| Trinity | r2013_08_14 | Assembling Trinity-limb (old) transcriptome |
| Trinity | 2.5.1 | Assembling Trinity-all, Trinity-gg, Trinity-S13, Trinity-S19, Trinity-S21, Trinity-S23, Trinity-limb (new) transcriptomes |
| universalmotif (R) | 1.12.1 | Converting JASPAR PWMs to RGT-HINT compatible format |
| VISTA | web VISTA | Identifying sequence homology between *Parhyale* and *Hyalella* |

#### Supplementary Methods

##### Trinity transcriptome assembly parameters

Trinity *de novo* transcriptome (all stages) was assembled with --max_memory 1500G and --CPU 32. Illumina NovaSeq short reads were supplied as a sample file, where each sample had two read files for read1 and read2. Read files were generated by first trimming transcripts using trim_galore, and then combining the val_1 reads with unpaired_1, and val_2 reads with unpaired_2 for each stage using cat. The same merged read files were used for stage-specific assemblies. For the genome-guided assembly, *in silico* normalized reads generated by Trinity for read1 and read2 across all stages were aligned to the genome using hisat2 --dta and then sorted using samtools. Trinity was run as Trinity --genome_guided_bam <bam file> --genome_guided_max_intron 300000.

##### StringTie2 transcriptome assembly parameters

Nanopore reads, when generated, are partitioned into chunks in different folders in the output. Reads were compiled by barcode. All reads were combined together into a single file and mapped to the genome using minimap2 -ax splice using the phaw_5.0.fa genome file as a target. StringTie2 L assembly was run using stringtie -L <nanopore_reads.bam>, while StringTie2 SL assembly was run using stringtie -mix <illumina_reads.bam> <nanopore_reads.bam>. Both StringTie2 assemblies used default settings.

##### Mikado setup parameters

Trinity transcriptomes were aligned to the phaw_5.0.fa genome using GMAP with --max-intronlength-ends=300000 --format=gff3_gene. All HISAT2 aligned read BAM files from Illumina RNA-Seq were fed into Portcullis using portcullis full --force --copy --verbose -t 32. Due to yet-to-be-resolved errors in the Portcullis package, we manually converted some assignments of the portcullis output using the following commands:

```
$ cat portcullis_out/2-junc/portcullis_all.junctions.tab |awk '!(($11=="?" && $14=="NA") ||
    ($11=="?" && $13=="NA"))' > portcullis_out/2-junc/portcullis_all.junctions.cleaned.tab
$ portcullis filter --verbose portcullis_out/1-prep
    portcullis_out/2-junc/portcullis_all.junctions.cleaned.tab
```

The scoring parameters and other variables for the Mikado configure file are found in the “20210630_list.txt” file. We applied penalties to the Trinity-based transcriptomes and bonuses to the StringTie2 transcriptomes based on our observations that Trinity transcripts, when aligned to the genome, showed many apparently spurious short transcripts at the RACE genes we examined.

For mikado configure, we used --mode permissive --scoring mammalian.yaml --copy-scoring mammalian.yaml -bt uniprot_sprot.fasta. We reasoned that the large genome size of *Parhyale* would be appropriate to analyze using the mammalian settings as opposed to the other available settings.

For the Transdecoder step, we ran the analysis with default settings. We omitted BLAST from our Mikado pipeline, as the extremely large file size of our mikado_prepared.fasta file proved intractable to straightforward BLASTX analysis. We estimated that it would take around 5 months to run using our current computing approaches. An updated Mikado transcriptome could be generated by breaking up the mikado_prepared.fasta file into many smaller BLASTX analyses, which could run much faster. The required files for further Mikado analyses are available in the GEO accession data.

##### eggNOG annotation

We generated a .pep file of the final Mikado transcriptome or the MAKER genome annotation using Transdecoder, and used emapper.py with default settings and -m diamond.

**OrthoFinder annotation**
Using a .pep file generated from the final Mikado transcriptome or the MAKER genome annotation using Transdecoder, we ran OrthoFinder to compare each transcript source to the database of all *Drosophila melanogaster* proteins found in the UNIPROT database.

**DESeq2 parameters**
For differential expression and accessibility analyses, we performed pairwise DESeq2 runs for adjacent pairs of developmental stages (e.g. S19 vs. S13, S21 vs. S19). Analysis was performed using standard settings. Peaks or genes were considered differentially expressed when padj < 0.05.

**ImpulseDE2 analysis**
We performed ImpulseDE2 analysis using standard settings, using the matrix of reads generated from bedtools multicov and used for DESeq2 analysis. For the Time parameter, we used the number of hours for each developmental stage, as indicated in the Browne et al. *Parhyale* staging guide (Browne et al., 2005). The time points were: 72, 77, 80, 87, 90, 96, 104, 112, 120, 132, 144, 155, 168, 180, and 192 hours, representing stages 'S13', 'S14', 'S15', 'S17', 'S18', 'S19', 'S19+', 'S20', 'S21', 'S22', 'S23', 'S24', 'S25', 'S26', 'S27', respectively. We extracted the IDE2 model fits for each peak (available at the GEO accession) as a .tsv file, used a custom Python function to plot accessibility trajectories (will be available at the GitHub repository). To generate such plots, we recreated the accessibility plotting function from the ImpulseDE2 package in R as two Python functions, shown below:

```
def ImpulseFunction(t, vecImpulseParam):
    return (1/vecImpulseParam[3]) * (vecImpulseParam[2] + (vecImpulseParam[3] -
      vecImpulseParam[2]) * (1/(1 + np.exp(-vecImpulseParam[1] * (t - vecImpulseParam[5]))))) *
      (vecImpulseParam[4] + (vecImpulseParam[3] - vecImpulseParam[4]) * (1/(1 +
      np.exp(vecImpulseParam[1] * (t - vecImpulseParam[6])))))

def ImpulseEvaluate(vecTimepoints, vecImpulseParam):
    vecImpulseValue = [ImpulseFunction(i, vecImpulseParam) for i in vecTimepoints]
    vecImpulseOutput = [i if i > 10^(-10) else 10^(-10) for i in vecImpulseValue]
    return vecImpulseOutput
```

**JASPAR2020 database**
For RGT-HINT analyses, the software expects each motif as a simple matrix representing position weights at each base for A, T, G, and C. This format is not readily available for the JASPAR2020 database, which utilizes the JASPAR format. We downloaded the full JASPAR2020 CORE non-redundant database, split the database into individual JASPAR format files for each transcription factor, and then converted that JASPAR format tile into a plain matrix file using the “universalmotif” R package. We also utilized the metadata extracted from this package to generate an .mtf file as required by RGT-HINT. All files for this type of analysis will be available on GEO at time of publication.
